# Spatial Single-Cell Atlas Reveals KSHV-Driven Broad Cellular Reprogramming, Progenitor Expansion, Immune and Vascular Remodeling in Kaposi’s Sarcoma

**DOI:** 10.1101/2025.09.01.673567

**Authors:** Wen Meng, Arun Das, Harsh Sinha, Rana Naous, Paige M. Bracci, Mike McGrath, Yufei Huang, Shou-Jiang Gao

**Author notes:** Correspondence: Yufei Huang, Cancer Virology Program, University of Pittsburgh Medical Center Hillman Cancer Center, Pittsburgh, PA 15213, USA., Shou-Jiang Gao, Cancer Virology Program, University of Pittsburgh Medical Center Hillman Cancer Center, Pittsburgh, PA 15213, USA. Wen Meng and Arun Das contributed equally to this work.

## Abstract

Kaposi’s sarcoma (KS) is a highly inflammatory, angiogenic tumor driven by Kaposi’s sarcoma-associated herpesvirus (KSHV), yet the origins of tumor cells and mechanisms of progression remain unclear. Here, we present the first spatial single-cell atlas of KS, profiling 256 samples across patch, plaque, and nodular lesions and normal controls. We identify CD34⁺ progenitor lymphatic endothelial cells (LECs) as the primary targets of KSHV, whose clonal expansion drives tumor growth. KSHV infection induces widespread cellular reprogramming across the tumor microenvironment, including LECs, vascular endothelial cells, fibroblasts, and macrophages, generating hybrid phenotypes that support angiogenesis, inflammation, and immune modulation. KSHV⁺ macrophages are enriched in tumor-proximal niches, further promoting a proangiogenic, immunosuppressive environment. Spatial analysis reveals evolving tumor-associated niches, with a core-to-periphery gradient correlating with infection, immune modulation, and cellular remodeling. We identify disease progression predictive signatures, offering mechanistic insights into KS pathogenesis and potential new therapeutic strategies by reprogramming the tumor microenvironment.

**Highlights:**

➣ CD34+ progenitor lymphatic endothelial cells are the primary KSHV target cells, with their clonal expansion driving Kaposi’s sarcoma growth.
➣ KSHV infection reprograms broad cell types into hybrid identities, and drives tumor-specific niche and vascular remodeling, endothelial plasticity, and immune modulation.
➣ KSHV-reprogrammed macrophages drive inflammation, angiogenesis, and immune modulation.
➣ Spatially resolved molecular and cellular signatures predict Kaposi’s sarcoma progression, offering novel therapeutic strategies targeting the tumor microenvironment.

## Introduction

Kaposi’s sarcoma (KS) is a highly vascularized and inflammatory tumor caused by infection with Kaposi’s sarcoma-associated herpesvirus (KSHV), also known as human herpesvirus 8 (HHV-8).^1^ KS is primarily recognized as an AIDS-related malignancy, but it also affects HIV-negative individuals, manifesting as Classic KS in elderly populations, Iatrogenic KS in immunosuppressive organ transplantation recipients, and Endemic KS in sub-Saharan Africa.^2^ Regardless of its diverse clinical contexts, KS pathology is universally characterized by chronic inflammation, dysregulated angiogenesis, and a complex tumor microenvironment (TME) rich in heterogeneous cell types.^3^ Central to this pathology is KSHV’s unique ability to manipulate the host immune response, reprogram infected cells, and orchestrate a proinflammatory, proangiogenic microenvironment that supports tumor initiation and progression.^4^

KSHV infection initiates inflammation and angiogenesis through a variety of mechanisms. Viral entry is mediated by receptors including integrins and ephrin receptor A2 (EphA2), which activate host signaling cascades resulting in the expression of key cytokines and angiogenic factors such as interleukin-6 (IL6), vascular endothelial growth factor (VEGF), and tumor necrosis factor-alpha (TNF-α).^5,6^ Furthermore, numerous latent and lytic KSHV genes and their products directly regulate proinflammatory and proangiogenic signaling pathways, or indirectly modulate immune cells, including macrophages (Mφs), neutrophils, natural killer cells (NKs), and T cells, altering the TME that fuel tumor growth.^7–9^ In vitro studies confirm that KSHV infection of endothelial cells and mesenchymal stem cells directly induces inflammatory and angiogenic responses, highlighting its capacity to reprogram host cells into pro-tumorigenic states critical for KS pathogenesis.^10–14^

The KS tumor comprises a mosaic of multiple cell types, including lymphatic endothelial cells (LECs), vascular endothelial cells (VECs), macrophages, fibroblasts (Fbs), mesenchymal stem cells, and progenitor cells.^15,16^ This cellular heterogeneity reflects the KSHV’s ability to induce phenotypic plasticity, driving the formation of hybrid or mixed cell identities that contribute to abnormal vasculature and chronic inflammation within lesions.^17^ For example, infection of mesenchymal stem cells induces expression of both VEC and LEC markers, such as CD31, VEGFR, Podoplanin (PDPN), and PROX1, implying that these cells may serve as progenitors for diverse endothelial cell lineages.^11,18–20^ Similarly, infection of VECs causes them to express LEC and progenitor cell markers, while LECs infected by KSHV also express markers of VEC and progenitor cells.^21–23^ This phenomenon underscores the dynamic cellular reprogramming that underlies KS tumor complexity, where cells can adopt mixed or hybrid phenotypes, contributing to the tumor’s abnormal vasculature and inflammatory milieu.

The cellular heterogeneity and plasticity have historically obscured the precise origin of KS tumor cells and the mechanism of KS progression.^24–27^ Recent advances in single-cell spatially resolved transcriptomics (scSRT) allow high-resolution mapping of gene expression in situ while preserving tissue architecture, crucial for understanding cellular interactions in the TME. Using the Xenium scSRT platform, we profiled 256 tissue samples from 43 KS patients across all clinical stages (patch, plaque, and nodular) alongside normal controls. This approach enabled simultaneous quantification of cellular and viral gene expression, illuminating spatial relationships among KSHV-infected cells, stromal elements, and immune infiltrates.

Here, we present the first comprehensive spatial atlas of KS at single-cell resolution, integrating viral and host transcriptional landscapes across disease stages. We reveal how KSHV-infected and uninfected cells convert into distinct lineages with spatially defined niches that orchestrate inflammation, angiogenesis, and microenvironmental remodeling during tumor progression. We have identified clonal expansion of progenitor-like CD34^+^LECs as a central driver of tumor growth, spatially constraining viral dissemination while inducing broad cross-lineage reprogramming of surrounding stromal and immune populations. Our data highlight macrophages as key effectors in KS pathogenesis, secreting inflammatory and angiogenic mediators, including IL6, TNF-α, and VEGF, that reinforce tumor-promoting inflammation and immune modulation. Together, these findings provide novel insights into the cellular programs and viral mechanisms underpinning KS, revealing potential therapeutic targets to disrupt the supportive TME and halt tumor progression. More broadly, our study offers a conceptual framework for understanding how oncogenic viruses exploit cellular plasticity, inflammatory signaling, and vascular remodeling to drive tumor evolution, with implications for other virus-associated cancers.

## Results

### Cell Composition and Spatial Dynamics Associated with KS Progression

We performed scSRT on tissue microarrays (TMAs) comprising normal skin tissues and KS lesions spanning patch, plaque, and nodular stages (Figure 1A). Using the Xenium platform (10X Genomics) and a custom designed panel of 306 genes targeting key major skin and immune cell markers, inflammatory and angiogenic regulators, tumor-associated genes, and representative KSHV latent and lytic genes (Table S1), we analyzed 256 cores of 2 mm^2^ arranged across 16 TMAs. These included 13 normal skin, 20 patch, 23 plaque, and 200 nodular samples (Table S2).

**Figure 1.**
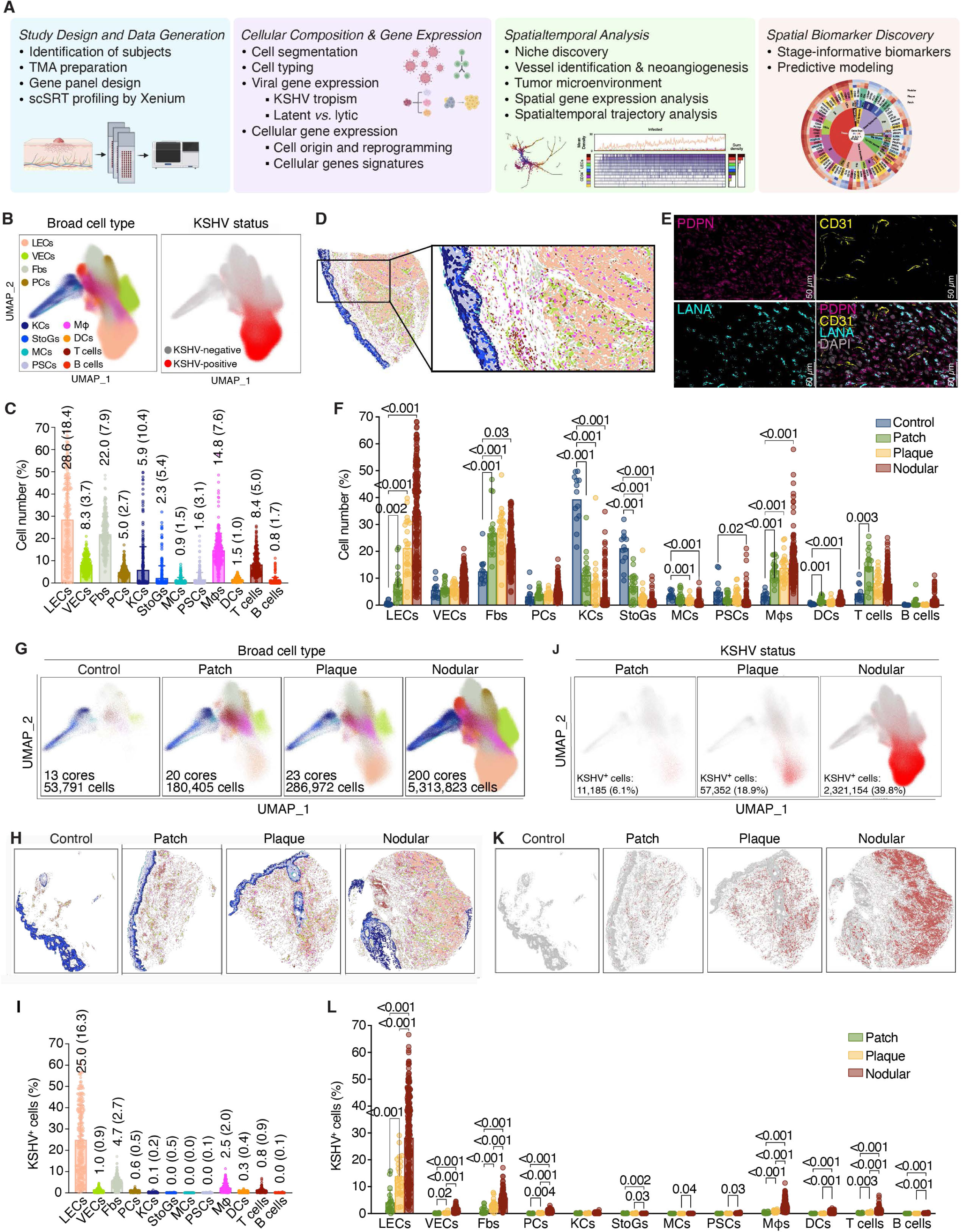
Cell composition, spatial dynamics, and KSHV cell tropism and transcriptional program associated with Kaposi’s sarcoma (KS) progression revealed by single-cell spatially resolved transcriptomics (scSRT) analyses. (A) Schematic illustration of scSRT pipeline and analysis workflow. scSRT data were generated from 256 cores of 2 mm^2^ arranged across 16 TMAs, including 13 normal skin, 20 patch, 23 plaque, and 200 nodular samples from 3 control subjects and 43 KS patients. Analysis workflow included cell segmentation, cell typing, viral and cellular gene expression, niche discovery, vessel identification, tumor microenvironment and trajectory analysis, disease stage-informative biomarker discovery, and predictive modeling. (B) The UMAP projection of the gene expressions of segmented cells colored based on cell type (left panel) and KSHV infection (right panel). (C) Scatter bar plot showing the percentages of identified cell types across all KS samples, highlighting lymphatic endothelial cells (LECs), vascular endothelial cells (VECs), fibroblasts (Fbs), and macrophages (Mφs) as the major cell types. Bars represent mean percentages and standard deviations (SD). (D) Spatial visualization of identified cell types in a nodular KS lesion. (E) Representative immunofluorescence staining images of PDPN (magenta), CD31 (yellow), LANA (cyan), and nuclei (grey) in a nodular KS lesion. (F) Scatter bar plot showing the percentages of identified cell types by KS stage, highlighting increases of LECs, fibroblasts and macrophages, and decreases skin structured cells keratinocytes (KCs) and spinous to granular cells (StoGs). Bars indicate the mean percentages with SD. Statistical significance was determined using mixed-effects model and Dunnett’s multiple comparisons test. P < 0.05 indicates significance. (G) The UMAP projection of different cell types by KS stage. (H) Spatial visualization of identified cell types by KS stage. (I) Scatter bar plot showing KSHV-infected percentages in identified cell types across all KS samples, highlighting LECs as the major infected cell type. Bars represent mean percentages with SD. (J) The UMAP projection of KSHV infection in different cell types by KS stage shown in (G). (K) Spatial visualization of KSHV infection in different cell types by KS stage shown in (H). (L) Scatter bar plot showing the percentages of KSHV-infected cells across identified cell types at different KS stages, highlighting marked increases of KSHV infection in LECs, fibroblasts, and macrophages along disease progression. Bars represent mean percentages with SD. Statistical significance was determined using RM two-way ANOVA model and Turkey’s multiple comparisons test. P < 0.05 indicates significance. Colors of different cell types shown in (B) are used in all other panels.

We developed a Density-Based Spatial Clustering of Applications with Noise (DBSCAN)-based algorithm to segment individual cores and assign spatial coordinates, ensuring consistency across all TMAs (see Methods). The analysis separated all 256 cores accurately and captured a total of 684,171,943 transcripts with a median of 117 transcript counts per cell, providing robust coverage for cell type annotation and spatial analysis.

To accurately annotate the cell types, we adapted a marker-based cell typing algorithm to assign cell types to cell clusters (Table S3) (see Methods). We identified 12 major cell types, including pilosebaceous cells (PSCs, 1.6%), keratinocytes (KCs, 5.9%), spinous to granular cells (StoGs, 2.3%), and melanocytes (MCs, 0.9%) as cells defining the skin layers; T cells (8.4%), macrophages (14.8%), B cells (0.8%), and dendritic cells (DCs, 1.5%) representing the immune landscape; pericytes (PCs, 5.0%), VECs (8.3%), and LECs (28.6%) defining the blood and lymphatic vessels; and fibroblasts (Fbs, 22.0%) representing the stromal compartment (Figure S1; Figure 1B and 1C). Further subtyping revealed five distinct fibroblast subtypes (secretory-reticular fibroblasts, SRFbs, 7.2%; proinflammatory fibroblasts, piFbs, 3.8%; myofibroblasts, myoFbs, 3.7%; secretory-papillary fibroblasts, spFbs, 2.4%; and mesenchymal fibroblasts, mFbs, 0.6%), three T-cell subtypes (exhausted CD8^+^ T cells, Tex, 3.5%; CD4^+^ T helper cells, Th, 1.7%; and regulatory CD4^+^ T cells, Treg, 0.1%), differentiated KCs (diffKCs, 2.5%), and proliferating LECs (ProlifLECs, 3.2%) (Figure S2A-S2F).

The UMAP projections of gene expression demonstrated distinct clusters of cell types, clearly delineating normal skin, endothelial cells of lymphatic and vascular origins, immune cells, and fibroblasts (Figure 1B). LECs and VECs form unique clusters, with VECs clustering closely with PCs, reflecting their common functions in supporting blood vessels. Interestingly, while macrophages clustered with immune cells (DCs, B cells, and T cells), a distinct subset clustered with LECs, indicating distinct functional roles in KS tumors. Spatial mapping of the cell types further highlighted the well-defined skin layers (KCs, StoGs, PSCs, and MCs), blood vessels (VECs and PCs), stromal fibroblasts, and tumor regions dominated by LECs and co-localized stromal and immune cells (Figure 1D). Immunofluorescence staining confirmed that PDPN^+^LECs, expressing KSHV latent protein LANA and endothelial marker CD31 were the main tumor cells (Figure 1E).

The cell composition analysis across KS stages revealed significant shifts in cellular proportions, primarily driven by a dramatic increase of LECs from 0.7% in normal skin to 33.3% in nodular lesions (Figure 1F-1H). VECs exhibited a modest increase from 6.6% in patch to 8.9% in nodular lesions, reflecting increased angiogenic activity. Macrophages increased moderately, comprising 15.8% in nodular lesions, a 2.8% increase from the patch stage, pointing to their potential role in tumor-associated inflammation and immune modulation. T cells declined from 14.4% in patch lesions to 8.1% in nodular lesions while the overall prevalence of B cells was low (<1%), indicating immune evasion. Fibroblasts, including both myoFbs and piFbs, increased by 3%-4%, reflecting extracellular matrix remodeling and tumor-stromal interactions. Conversely, KCs, StoGs and MCs, dominant in normal skin, were sharply depleted, dropping from 39.5% to 3.0%, from 21.2% to 0.5%, and from 3.7% to 0.5% in the nodular stage, respectively, indicative of disrupted skin architecture in the dermal and epidermal layers.

Taken together, these results mirror the pathological manifestations of KS tumors,^28,29^ revealing the cell composition and spatial dynamics of KS lesions. This includes the evolving cellular heterogeneity, characterized by a shift from a balanced tissue architecture to a tumor-dominant environment marked by tumor-associated cell proliferation, stromal remodeling, immune dysregulation, and disrupted normal tissue architecture.

To understand the molecular mechanisms underlying the cellular and spatial dynamics, we identified differentially expressed genes along KS progression by comparing them with control tissues. As expected, there was a distinct population of cells exhibiting a robust endothelial gene expression signature (Figure S3). Notably, we observed a significant upregulation of key genes associated with endothelial identity and function, including PROX1, LYVE1, KDR (VEGFR2), CD34, RAMP2, CLDN5, FABP4, and PLVAP. These genes are critical regulators of angiogenesis, vascular permeability, and lymphatic specification, suggesting active vascular differentiation or activation.^30,31^ The elevated expression of CXCR4, ITGB3, CD93, and MARCKSL1, further signifies endothelial migration, signaling, and cytoskeletal remodeling.

Specifically, PROX1 and LYVE1 are hallmarks of LECs, whereas KDR, CD34, and RAMP2 are prominent in blood VECs,^30,31^ indicating a hybrid or transitional endothelial phenotype. The enrichment of these genes implies active vascular remodeling and endothelial plasticity in response to KSHV infection and the altered TME.

In contrast, genes associated with mesenchymal, fibroblastic, or mural cell identities are significantly downregulated, including LUM, COL6A2, COL6A3, PDGFRB, and POSTN (Figure S3), all of which are involved in extracellular matrix organization, fibrosis, and stromal support.^30,31^ The reduction in CXCL12 and IFITM1 expression suggests an inhibitory mechanism for chemokine-mediated immune cell recruitment and interferon signaling.^32,33^ Additionally, the repression of IL3RA and TAGLN, which are linked to myeloid and smooth muscle lineages, further highlights the loss of non-endothelial characteristics during transition to an endothelial phenotype.^34,35^

Taken together, these findings reveal a transcriptional program consistent with endothelial reprogramming and vascular specialization, coupled with a shift toward mesenchymal and inflammatory signatures. This suggests the emergence or expansion of a distinct endothelial cell population, potentially involved in tissue remodeling, angiogenesis, or KS progression, including tumor-associated vasculature or lymphangiogenesis. These changes are accompanied by evasion of both innate and adaptive immunity, which may facilitate KS tumor progression.

### Broad KSHV Cell Tropism and KSHV-Driven Cell Proliferation in KS Progression

KSHV infection was predominantly detected in LECs (77.7%), followed by fibroblasts (23.5%), macrophages (17.8%), and VECs (13.1%), which together comprising over 90% of the infected cells, revealing broad KSHV cell tropism (Figure 1I). The infection dynamics showed a stage-specific progression, with the proportion of KSHV^+^ cells increasing from 6.1% in patch lesions to 39.8% in nodular lesions (Figure 1J). LECs had the highest infection rates, increasing from 44.2% in patch lesions to 82.6% in nodular lesions (Figure 1H and 1J-1L). VECs, fibroblasts, and macrophages also showed modest increases in infection rates, rising from 1.4% to 15.3%, 3.5% to 27.0% and from 2.1% to 20.7%, respectively. These results suggest a stage-specific shift in KSHV infection dynamics, where VECs, fibroblasts, and macrophages likely facilitate early viral dissemination, and LECs become the dominant infected cell population in later stages, driving tumor progression. UMAP projections also revealed these infection dynamics, with KSHV^+^ cells predominantly clustering within the LEC population and showing a notable increase in density in the nodular stage (Figure 1G and 1J). Spatial analysis across stages revealed a transition from small, dispersed regions of infected cells in patch lesions to densely packed tumor regions in nodular lesions (Figure 1H and 1K). The expansion of the LEC population, accompanied by secondary infection of immune cells and vasculature, underscores the critical role of KSHV infection and the remodeling of the TME in driving KS progression.

### KSHV Replication Program and KSHV-Encoded vIL6 in KS Progression

While KSHV latent infection is critical for KS progression by sustaining cell proliferation, survival, and immune evasion, a small subset of infected cells have been found to express viral lytic genes in KS tumors.^36^ These lytic cells, which undergo either full or abortive replication, produce virus-encoded cytokines and induce cellular proinflammatory and proangiogenic factors, promoting KS progression through both autocrine and paracrine mechanisms. We found that 46.5% of KSHV^+^ cells were in latent phase, expressing latent transcripts such as ORF71, ORF72 and ORF73, while 16.1% of the cells were in the lytic phase, expressing viral lytic transcripts ORF50 (RTA), ORF57, ORF59, ORF-K9, and ORF65 (Figure 2A). Surprisingly, 37.4% of KSHV^+^ cells expressed viral interleukin-6 (vIL6), encoded by a KSHV early gene ORF-K2 implicated in proinflammatory and proangiogenic functions and immune modulation.^37^ Among these vIL6^+^ cells, 28.6% also expressed viral lytic genes, indicating that vIL6 expression may be regulated by both the viral transactivator RTA and other mechanisms.^38–40^ This pattern of KSHV gene expression persisted across all KSHV^+^ cell types (Figure S4). While the number of lytic cells did not vary significantly across KS stages, the proportion of vIL6^+^ cells progressively increased from 18.2% in patch lesions to 41.5% in nodular lesions (Figure 2B), predominantly in tumor regions, with LECs being the major vIL6^+^ cells at 40.2% and increased along with disease progression from 17.0% in patch lesions to 44.8% in nodular lesions (Figure 2C and 2D; Figure S5). Immunofluorescence staining confirmed the presence of phosphorylated gp130 (p-gp130) and phosphorylated STAT3 (p-STAT3) in KS tumors, indicating active vIL6 signaling, particularly in KSHV^+^LECs (Figure 2E and 2F). These results suggest that KSHV lytic replication, particularly vIL6, plays a critical role in KS progression by driving inflammation and angiogenesis through autocrine and paracrine mechanisms.

**Figure 2.**
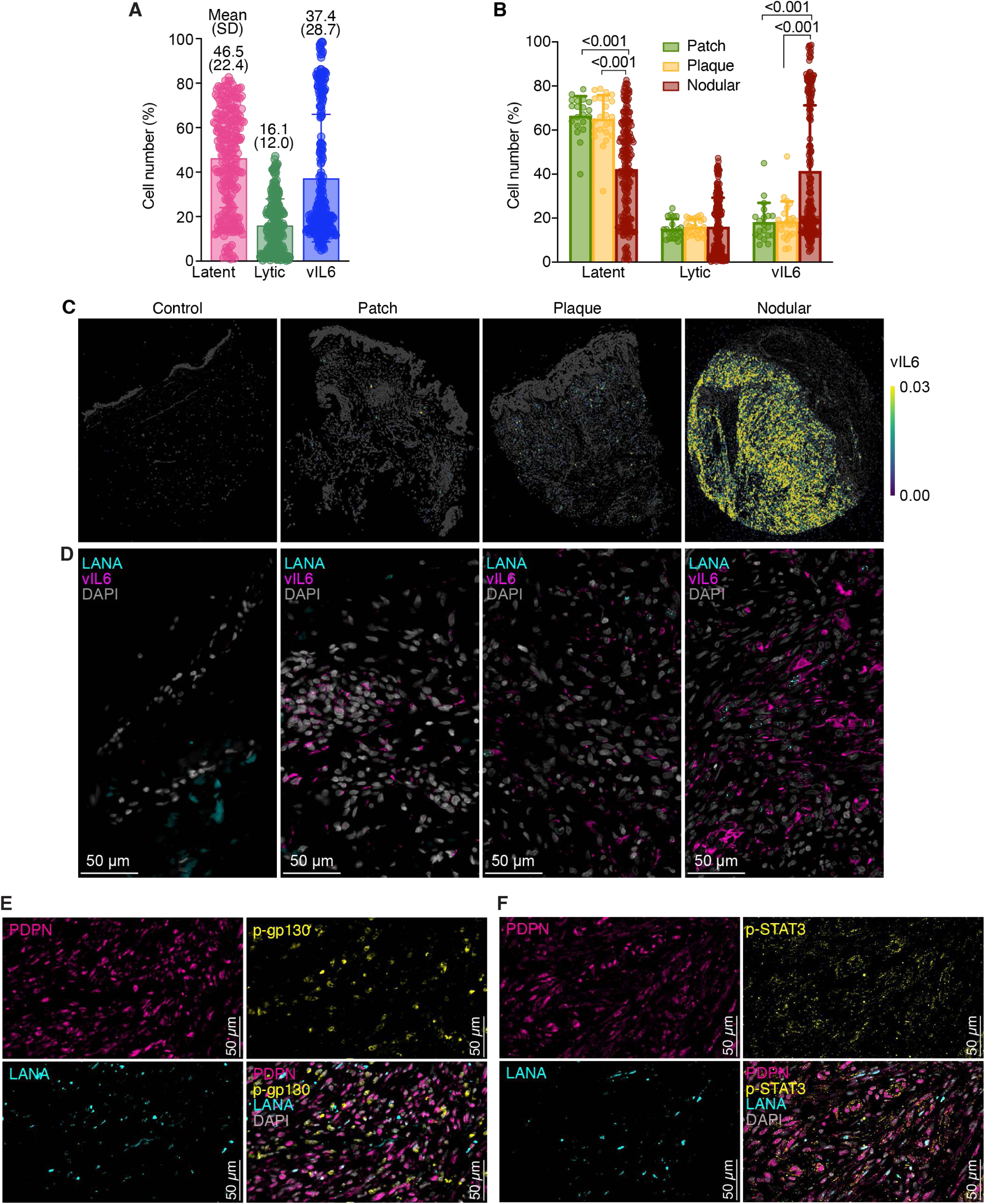
KSHV replication program and KSHV-encoded vIL6 in Kaposi’s sarcoma (KS) progression. (A) Scatter bar plot showing the percentages of KSHV latent, lytic and vIL6-expressing cells across all KS samples, highlighting the abundance of KSHV lytic gene expression, particularly vIL6 in KS tumors. Bars represent mean percentages and standard deviations (SD). (B) Scatter bar plot showing the percentages of KSHV latent, lytic and vIL6-expressing cells by KS stage, highlighting the increased vIL6 expressing cells at the advanced disease stage. Bars represent mean percentages with SD. Statistical significance was determined using RM two-way ANOVA model and Turkey’s multiple comparisons test. P < 0.05 indicates significance. (C) Spatial visualization of vIL6 transcript in different stages of KS lesions. (D) Representative immunofluorescence staining images of LANA (cyan), vIL6 (magenta) and nuclei (grey) in different stages of KS lesions. (E-F) Representative immunofluorescence staining images of p-gp130 (yellow, E) or PDPN (yellow, F) together with LANA (cyan), PDPN (magenta) and nuclei (grey) in a nodular KS lesion.

### Spatial Analysis Reveals Tumor-Driven Niches and Immune Shifts in KS Progression

Spatial analysis of KS tissues identified 11 distinct cellular niches, which were grouped into four main categories based on functional or anatomical characteristics: Peripheral skin-associated (Basal epidermis and Differentiated epidermis, Diff epidermis); Vascular-associated (Tumor-associated VEC stroma, TA VEC stroma; VEC stroma; and Stroma), Immune-enriched (T-cell immune stroma; Macrophage immune stroma; and Immune stroma), and Tumor-associated (Tumor core; Tumor; and Tumor boundary) niches (Figure 3A-3C). Differential expression analysis revealed niche-specific gene patterns corresponding to the respective niche categories (Figure S6).

**Figure 3.**
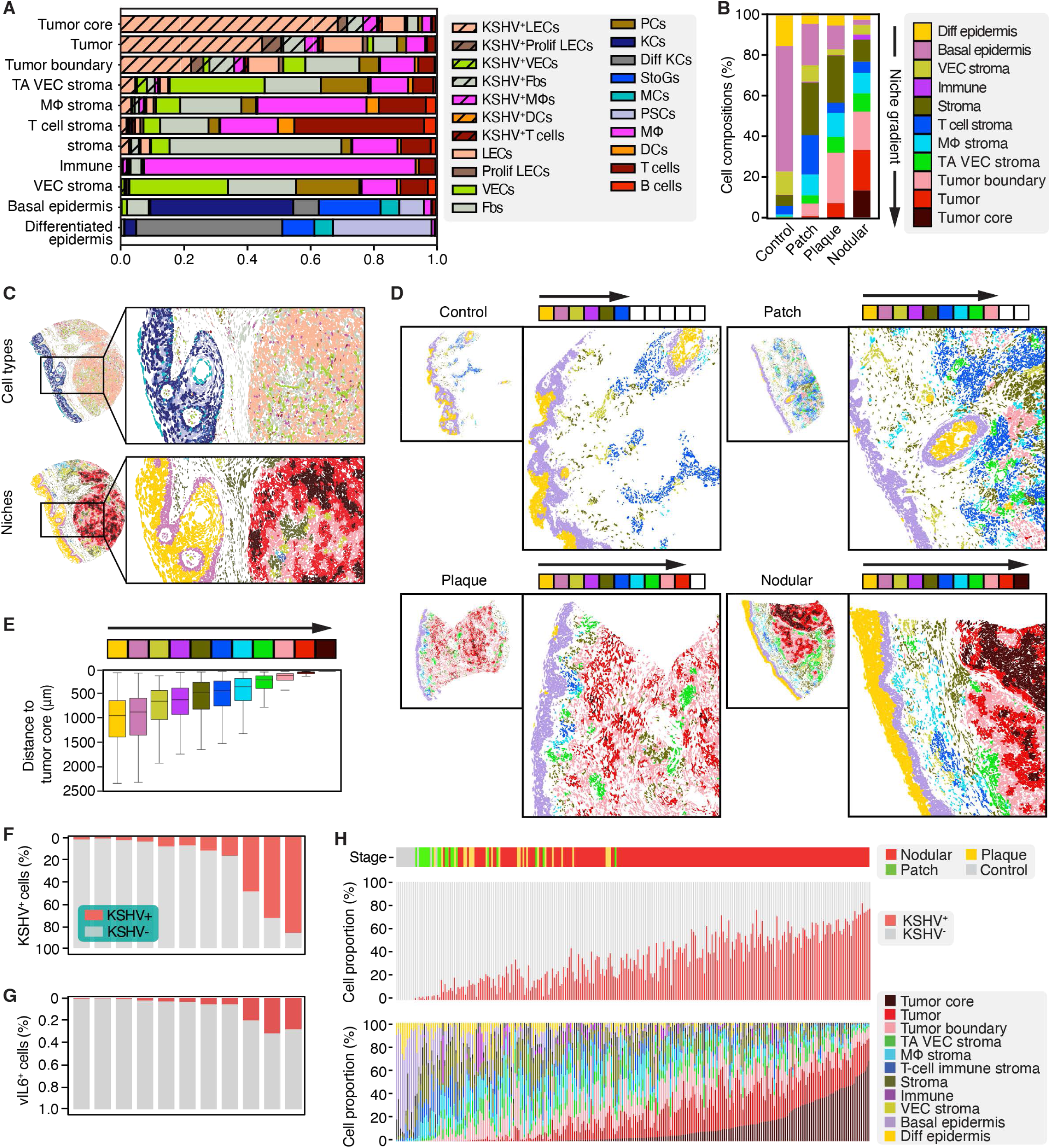
Identification of tumor-driven niches in Kaposi’s sarcoma (KS) progression by spatial analysis. (A) Cell type composition of 11 distinct cellular niches identified in KS lesions. They were grouped into four groups based on functional or anatomical characteristics. Peripheral skin-associated niches include Basal epidermis and Differentiated epidermis (Diff epidermis). Vascular-associated niches include Tumor-associated stroma (TA VEC stroma), VEC stroma, and Stroma. Immune-enriched niches include T-cell immune stroma, Macrophage immune stroma, and Immune stroma. Tumor-associated niches include Tumor core, Tumor and Tumor boundary. (B) Compositions of the identified niches by KS stage. (C) Spatial visualization of the identified niches in a nodular KS lesion (lower panel) and the matched spatial map of cell types (top panel). (D) Spatial visualization of the identified niches by KS stage revealing dynamic changes of niches from the epidermis to the Tumor core correlated with KS progression. (E) Structured gradient of niches revealed by distance-to-tumor-core analysis in KS lesions. Bars represent median distances to Tumor core with interquartile ranges (top 75% and lower 25%) and extended whiskers were created using Tukey method. (F-G) Correlation of niche gradient with KSHV infection rate (F) and vIL6 expression (G). Bars represent median percentages at different niches. (H) Dynamic changes of niche compositions by KS stage showing increases of Tumor-associated niches, and declines of Tumor-associated stroma niches, Immune-enriched niches, and Peripheral skin-associated niches along KS progression. Individual KS lesions were ordered based on the percentages of cells in Tumor core niche.

The Peripheral skin-associated niches play key roles in maintaining skin integrity, such as epidermis development and intermediate filament organization. The Basal epidermis niche was rich in KCs (45.3%), StoGs (19.6%), and fibroblasts (6.9%), indicating a mixture of epidermal and dermal components (Figure 3A and 3B). The Differentiated epidermis niche was dominated by differentiated KCs (46.3%) and PSCs (31.2%), reflecting its role in skin appendages. These niches were predominant in control tissues but declined significantly in advanced stages of KS, being replaced by Tumor- and Vascular-associated niches (Figure 3B). Spatial plots of the niches further revealed the dominant skin niches in the control samples as well as early patch lesions (Figure 3D). Tumor-associated niches, such as the Tumor core and Tumor, emerged during later disease stages, with the Tumor core dominated by KSHV^+^LECs (71.8%), expanding from 71.6% in patch lesions to 73.6% in nodular lesions (Figure 3B). The Tumor boundary, containing KSHV^+^LECs (26.4%) and fibroblasts (24.9%), supported stromal remodeling and immune modulation, reinforcing the spatial heterogeneity of KS lesions, especially in the plaque and nodular lesions (Figure 3D). The complex TME of the Tumor-associated niches revealed enriched pathways of regulation of endothelial cell apoptotic process, vascular wound healing, regulation of angiogenesis, positive regulation of vasculogenesis, angiogenesis involved in wound healing, and endothelial cell differentiation, further confirming the pro-tumoral and angiogenic nature of the TME (Figure S6).

Vascular-associated niches presented enriched pathways of epithelial mesenchymal transition (EMT) and positive regulation of vascular associated smooth muscle cell proliferation, etc., confirming their roles in vascularization (Figure S6).

Tumor-associated vascular stroma, which is closer to the Tumor-associated niches, expanded from 4.3% in patch to 8.4% in nodular lesions, facilitating tumor growth through angiogenesis, whereas Vascular stroma declined from 8.6% to 4.3%, indicating a shift toward tumor-driven vascularization (Figure 3B).

Immune-enriched niches were enriched with pathways of EMT, inflammatory response and TNF-α signaling via NF-κB, and apoptosis, revealing a complex immune microenvironment that supported tumor suppression and dysregulated immune responses (Figure S6). Interestingly, these niches showed opposing trends within each other, with T-cell immune stroma peaking at 19.7% in patch stages but declining to 5.6% in nodular lesions, suggesting weakening immune defenses (Figure 3B).

Meanwhile, Macrophage immune stroma increased from 3.5% to 7.5%, highlighting the increasing role of macrophages in later KS stages. The Immune niche, dense with macrophages, was most pronounced in nodular lesions (9.4%) compared to plaque lesions (3.7%), emphasizing the shifting landscape from immune response to tumor promotion during KS progression (Figure 3B).

Our distance-to-tumor-core analysis quantitatively mapped the spatial arrangement of niches within KS tissues, revealing a strikingly structured gradient of niches extending from the Tumor core outward to the peripheral compartments (Figure 3E). This gradient correlated with KSHV infection rates and vIL6 expression, with Tumor-associated niches exhibiting the highest levels and Peripheral skin-associated niches having the lowest (Figure 3F and 3G). As expected, Tumor core, Tumor and Tumor boundary niches were positioned at the center, with minimal distances to the Tumor core, reinforcing their direct involvement in tumor growth and KSHV-driven oncogenesis. In contrast, Vascular-associated (Tumor-associated, VEC stroma, and Stroma), and Immune-enriched (T-cell immune stroma, Macrophage immune stroma, and Immune stroma) niches occupied intermediate distances, indicating their peripheral roles in TME (Figure 3E). The most distant niches were the Peripheral skin-associated compartments, highlighting the progressive spatial expansion of lesions into the epidermis. This structured organization suggests that KS progression follows a defined spatial trajectory, where tumor-associated and angiogenic niches remain central, while immune and stromal components were positioned at increasing distances, possibly reflecting their protective or reactive roles against tumor invasion or their reprogrammed inflammatory roles that favor tumor progression. This well-defined tumor-core-to-periphery gradient across different KS stages (Figure 3D), suggests a conserved spatial architecture, which might be crucial for tumor development and immune evasion.

To assess how KSHV infection and lytic activity distributed across the spatial niche gradient, we quantified both KSHV-infected cells and the expression of vIL6 across the 11 defined niches. Infection was strongly enriched in the Tumor-associated niches (Tumor Core, Tumor, and Tumor Boundary) with high infection rates reaching 83% in the Tumor Core (Figure 3F). In contrast, the Immune and Skin-associated niches showed minimal infection, suggesting a spatially restricted infection within TME where KSHV dissemination is largely confined to the tumor-permissive environments. vIL6 expression mirrored the tumor niche dominance and the infection pattern along the niche gradient, remaining largely absent in the non-tumoral niches (Figure 3G). These findings suggests that vIL6 is spatially restricted to the niches with high tumor burden and viral load.

By ordering individual KS tissues based on the percentages of cells in Tumor core niche, a clear progression of niche dominance with lesion stages emerges (Figure 3H). We observed that advanced nodular lesions showed a pronounced enrichment of Tumor core, Tumor, and Tumor boundary niches, whereas early patch lesions (green) exhibited a higher abundance of T-cell immune stroma, Macrophage immune stroma, and baseline Vascular niches (Figure 3H). This trend suggests that as KS advanced, Tumor-associated niches become increasingly dominant, while Immune-enriched and Vascular-associated niches decline. The inverse relationship between tumor burden and immune presence highlights a potential shift from an immunosuppressive to a proinflammatory and angiogenic state, which might facilitate tumor progression. These compositional differences across KS stages suggest that the ratio of Tumor-associated to Immune-enriched niches could serve as a biomarker for disease progression and aggressiveness. The gradual depletion of immune niches in advanced lesions might indicate immune evasion by KSHV^+^ tumor cells. These proinflammatory and immunity shifts could be potentially exploited for novel therapeutic strategies aimed at reprogramming the TME or restoring immune activity in KS tumors.

### KSHV Infection Induces Transcriptional Reprogramming in Diverse Cell Types

KS tumor cells are known to express a mix of cellular markers, including those for LECs, VECs, macrophages and fibroblasts, as well as progenitor and mesenchymal markers.^41–43^ We investigated the transcriptional changes in various cell types, confirming that KSHV infection reprograms LECs, VECs, fibroblasts, and macrophages. Both KSHV^+^LECs and KSHV^-^LECs expressed VEC, fibroblast, and macrophage markers, with infected cells showing higher expression rates for these markers compared to uninfected counterparts (74.3% *vs.* 61.2%, p<0.001 for VEC marker; 39.1% *vs.* 32.1%, p<0.001 for fibroblast marker; and 28.7% *vs.* 21.9%, p<0.001 for macrophage marker) (Figure 4A), suggesting cellular reprogramming driven by both direct viral infection and paracrine signaling. Furthermore, the expression of VEC, macrophage, and fibroblast markers in KSHV^+^LECs remained largely stable, while KSHV^-^LECs exhibited increases as the tumor progressed (Figure 4B), reaffirming the paracrine effect in reprogramming during tumor progression.

**Figure 4.**
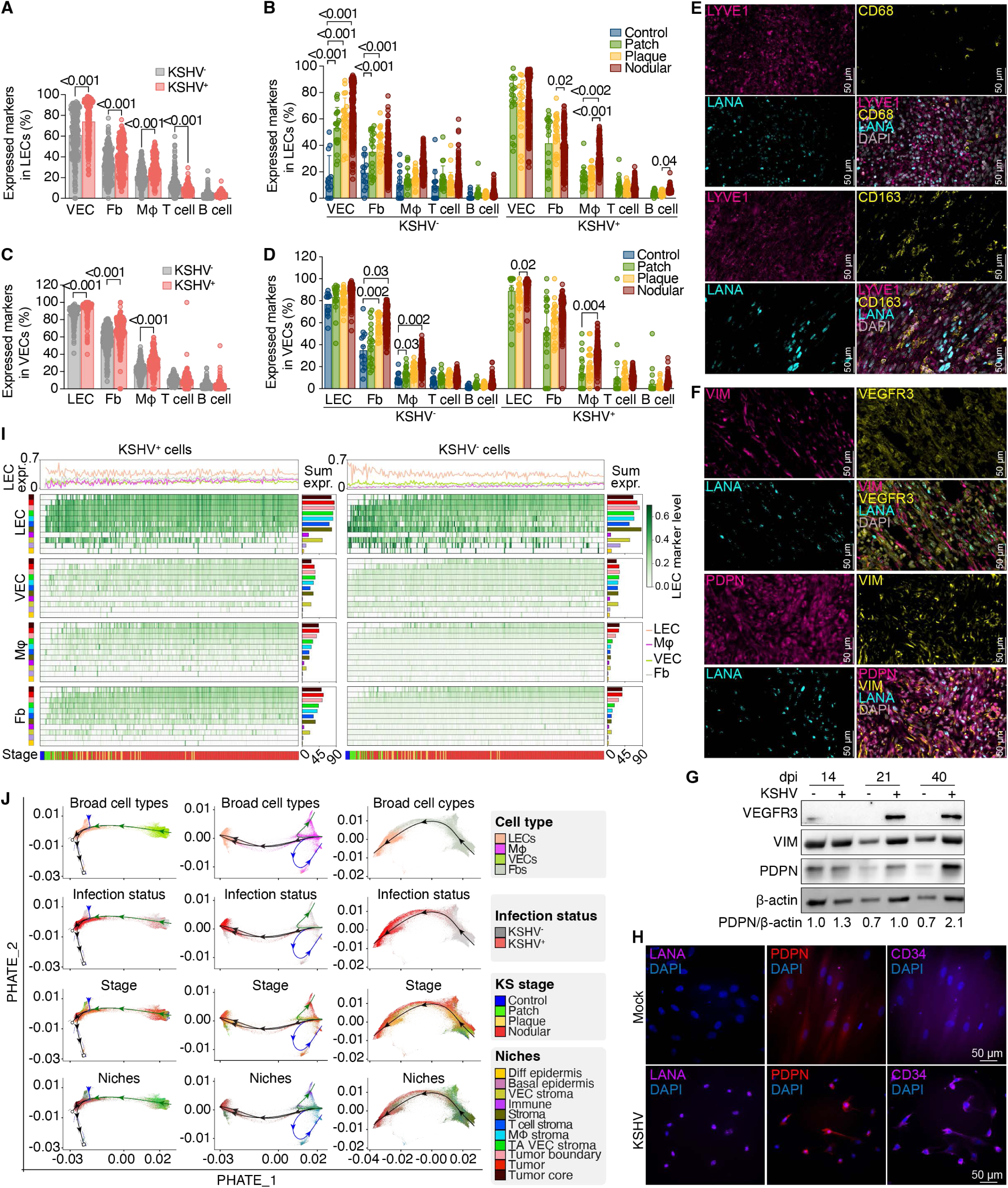
Induction of transcriptional reprogramming in diverse cell types in Kaposi’s sarcoma (KS) by KSHV infection. (A) Scatter bar plot showing the percentages of lymphatic endothelial cells (LECs) expressing markers of different cell types, including vascular endothelial cells (VECs), fibroblasts (Fbs), macrophages (Mφs), T cells, and B cells, across all KS samples, highlighting the heterogeneity and plasticity of LECs in KS lesions. Bars represent mean percentages and standard deviations (SD). (B) Scatter bar plot showing the percentages of LECs expressing markers of different cell types by KS stage, highlighting the increases of uninfected VECs and Fbs along disease progression. Bars represent mean percentages with SD. (C) Scatter bar plot showing the percentages of VECs expressing markers of different cell types, including LECs, fibroblasts, macrophages, T cells, and B cells, across all KS samples, highlighting the mixed identities and plasticity of VECs in KS lesions. Bars represent mean percentages with SD. (D) Scatter bar plot showing the percentage of VECs expressing markers of different cell types by KS stage along disease progression. Bars represent mean percentages with SD. (E) Representative images of immunofluorescence staining of LIVE1 (magenta), LANA (cyan), and nuclei (grey) together with CD68 (yellow, upper panels) and CD163 (yellow, lower panels) in a nodular KS lesion. (F) Representative images of immunofluorescence staining of a nodular KS lesion with VEGFR3 (yellow), VIM (magenta), LANA (cyan), and nuclei (grey, upper panels); and PDPN (magenta), VIM (yellow), LANA (cyan), and nuclei (grey, lower panels) in a nodular KS lesion. (G) Western-blots showing KSHV infection of primary human dermal fibroblasts (HDFa) induces the expression of VEGFR3, PDPN and VIM. β-actin served as the loading control. (H) Immunofluorescence staining images showing KSHV infection of HDFa induces the expression of PDPN and CD34. KSHV-infected cells are identified by LANA staining. Nuclei were stained with DAPI. (I) Space-time heatmaps of LEC marker expression in LECs, VECs, macrophages and fibroblasts across KS tissue niches and stages. KSHV^+^LECs maintain strong lymphatic identity within Tumor and Stromal niches, while KSHV-infected VECs, macrophages, and fibroblasts progressively acquire LEC-like features, primarily in Tumor-associated niches. KSHV^-^LECs show high marker expression in controls and early lesions that modestly decline with disease progression, and uninfected stromal cells exhibit low, niche-restricted LEC marker expression. Niches with colors described in Figure 3B are shown on both sides of the panels with the sum of LEC marker expression (Sum expr.) in different niches in the right panels, and in different disease stages in the top panel (LEC expr.). (J) Trajectory analysis using PHATE reveals KSHV-driven reprogramming of VECs, macrophages, and fibroblasts toward a LEC-like, KSHV^+^ state enriched in Tumor-associated regions. Macrophages follow two distinct trajectories, giving rise to proinflammatory cells in peripheral immune niches and immunomodulatory cells at the tumor margin. Statistical significance was determined using mixed-effects model and Dunnett’s multiple comparisons test (A-D). P < 0.05 indicates significance.

VECs are known to express LEC markers,^44^ and indeed, 90% VECs expressed these markers (Figure 4C). Interestingly, 62% of VECs also expressed fibroblast markers, and 25% expressed macrophage markers. However, the proportions of KSHV^+^VECs and KSHV^-^VECs expressing fibroblast and macrophage markers were similar (Figure 4C), with an increase in fibroblast marker expression in KSHV^-^VECs in advanced tumor stages (Figure 4D), confirming the role of stromal remodeling in KS progression.

Immunofluorescence imaging confirmed the co-expression of multiple markers in both LANA^+^ and LANA^-^ cells, including LEC marker LYVE1 together with macrophage and monocyte markers CD68 and CD163 (Figure 4E), as well as LEC markers VEGFR3 and PDPN with the mesenchymal marker vimentin (VIM) (Figure 4F). Western-blotting and immunofluorescence imaging further detected increased levels of VEGFR3, PDPN, CD34, and VIM in primary human dermal fibroblasts following KSHV infection (Figure 4G and 4H). These results confirm the extensive reprograming of different cell types by KSHV infection.

To gain deeper insight into the spatial and temporal dynamics of different cell types in KS progression, we developed a novel space-time representation framework that depicts diverse cellular and molecular features along a continuous niche gradient and stage-aligned pseudotime (see Methods), enabling high-resolution tracking of KSHV infection and cell-state transitions across tissue niches and disease stages.

Space-time heatmaps of LEC marker expression confirmed sustained LEC marker expression in KSHV^+^LECs within Tumor and Stromal niches, while KSHV^+^VECs, KSHV^+^ macrophages, and KSHV^+^ fibroblasts progressively acquired lymphatic identity, primarily within Tumor-associated niches (Figure 4I, left panel). KSHV^-^LECs showed strong lymphatic marker expression in control and early-stage lesions, which declined modestly as disease progressed, likely due to suppression by tumor-induced remodeling (Figure 4I, right panel). In contrast, uninfected stromal cells (VECs, macrophages, and fibroblasts) showed low levels of LEC marker expression, which was spatially restricted to Tumor niches, further suggesting paracrine signaling from adjacent infected cells.

Trajectory inference-based gene expression analysis further demonstrated the expansion of KSHV^+^LECs supported by transcriptional reprogramming of multiple stromal lineages. PHATE analysis revealed that the trajectories of VECs, macrophages, and fibroblasts converged toward a LEC-like, KSHV-infected state enriched in Tumor-associated regions (Figure 4J). These transitions reflect KSHV-driven reprogramming of vascular, immune, and mesenchymal compartments toward a lymphatic endothelial-like identity. Additionally, macrophages exhibited two transcriptionally and spatially distinct trajectories: one leading to proinflammatory macrophages localized to peripheral immune niches and the other to immune-modulatory macrophages at the tumor margin (Figure 4J). These findings highlight both direct and indirect routes through which KSHV drives the expansion of the LEC-like population and reshapes the TME, orchestrating niche-specific transcriptional remodeling to support tumorigenesis.

### Progenitor-Like LECs Are the Origins of KSHV-Targeted Cells that Drive KS Tumorigenesis Through Clonal Expansion and Niche Remodeling

KS tumor cells express the CD34 progenitor cell marker in addition to LEC markers.^41^ Our results have shown the expansion of KSHV^+^LECs and reprogramming of different cell types along KS tumor progression. These findings solidify KSHV infection as the main driving force behind KS tumor progression and suggest that the origin of KS tumor cells might involve progenitor cells in addition to other differentiated cells, such as LECs, VECs, fibroblasts, and macrophages, which could be induced to partially differentiate or dedifferentiate upon KSHV infection. Previous studies have shown that KSHV can reprogram progenitor cells, such as mesenchymal stem cells.^11,18–20^ Given that LECs represent the most abundant tumor cells (Figure 1F), we examined CD34 expression in these cells. Across all TMA cores, KSHV^+^LECs showed a significant enrichment of CD34 expression compared to KSHV^-^LECs (69.4% *vs.* 18.2%, P < 0.0001) (Figure 5A). Hence, KSHV⁺CD34⁺LECs comprised the dominant population, while KSHV⁺CD34⁻LECs were rare, suggesting that KSHV infection either targets CD34⁺ progenitor cells or induces CD34 expression upon infection of LECs. In contrast, VECs represented only 8.4% of KS lesions, with <1% infected by KSHV (Figure 1C and 1I) even though over >89.0% of VECs expressed CD34 (Figure 5B). Importantly, KSHV⁺CD34⁺LECs increased from 39.0% in patch lesions to 52.0% in plaque lesions and 79.0% in nodular lesions, ultimately comprising the dominant population in advanced disease (Figure 5C). Meanwhile, KSHV^-^CD34⁺LECs declined, and CD34⁻LECs remained at negligible level and essentially virus-free as tumor progressed (Figure 5C). Interestingly, KSHV⁺CD34⁺VECs also showed a slight increase, from 1.3% in patch lesions to 3.6% in plaque lesions and 14.5% in nodular lesions, suggesting viral infection spread from KSHV⁺CD34⁺LECs, which causes a minor expansion of KSHV⁺CD34⁺VECs (Figure 5D). CD34⁻VECs and KSHV⁻VECs remained unchanged throughout tumor progression (Figure 5D). These findings support a model in which CD34⁺LECs served as the initial KSHV primary target cells, which persist as a proliferative, angiogenic reservoir in KS tumors. Furthermore, KSHV-associated tumor expansion is tightly coupled to CD34⁺ progenitor-like identity.

**Figure 5.**
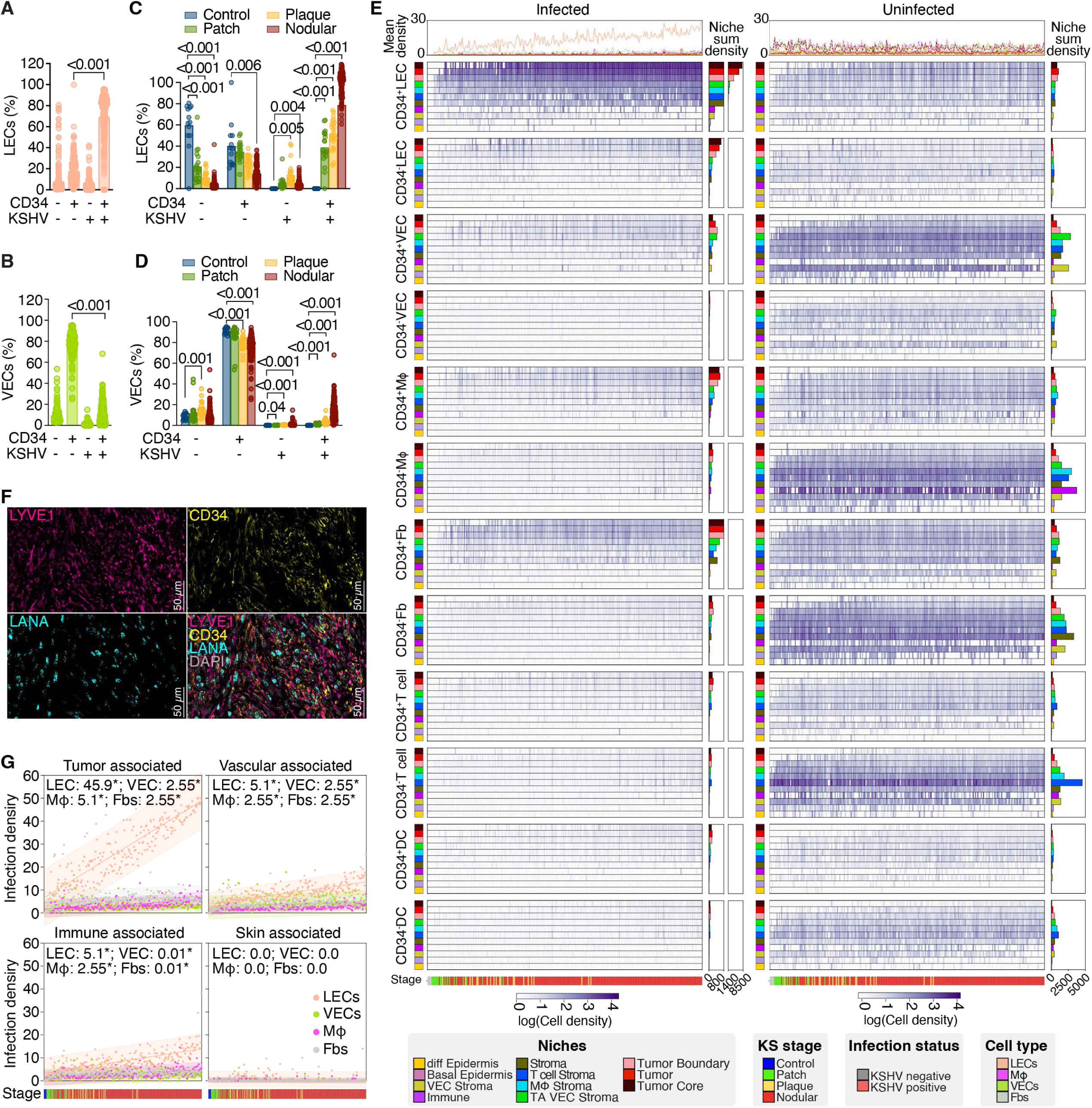
Progenitor-like lymphatic endothelial cells (LECs) as the origins of KSHV-targeted cells, driving Kaposi’s sarcoma (KS) tumorigenesis through clonal expansion and niche remodeling. (A) Scatter bar plot showing the percentages of LECs expressing CD34 and infected by KSHV, highlighting that most LECs in KS lesions are both KSHV-infected and CD34-positive. Bars represent mean percentages and standard deviations (SD). (B) Scatter bar plot showing the percentages of LECs expressing CD34 and infected by KSHV by KS stage, highlighting the increase of KSHV^+^CD34^+^LECs and the decrease of KSHV^-^CD34^-^LECs along disease progression. Bars indicate the mean percentages with SD. (C) Scatter bar plot showing the percentages of vascular endothelial cells (VECs) expressing CD34 and infected by KSHV, highlighting that most VECs in KS lesions are CD34-positive but KSHV-negative. Bars indicate the mean percentages with SD. (D) Scatter bar plot showing the percentages of VECs expressing CD34 and infected by KSHV by KS stage. Bars indicate the mean percentages with SD. (E) Space-time heatmaps of cell density in different types of cells stratified by CD34 expression and infection status, highlighting that KSHV⁺CD34⁺LECs appear early in disease development and act as the primary viral amplifiers in KS lesions, driving secondary infection in VECs, fibroblasts, and macrophages in a niche-specific manner, while uninfected stromal and immune cells are spatially reorganized in response to LEC expansion. The sum of cell density in different niches (Niche sum density) are shown in the right panels, and in different disease stages in the top panel (Mean density). (F) Representative immunofluorescence staining images of LIVE1 (magenta), CD34 (yellow), LANA (cyan), and nuclei (grey) in a nodular KS lesion. (G) Scatterplots of infected cell density by KS stage with fitted line in different niche regions, showing that KSHV⁺ cell densities increase predominantly in Tumor-associated niches, driven by LEC expansion, with smaller or moderate increases in other Stromal and Immune niches, while Skin-associated niches remain largely unaffected. The slopes of the fitted lines, denoting increase in infected cells per μm^2^ across KS progression from control to nodular, are shown on the top of the plots. Statistical significance was determined using ordinary one-way ANOVA model and Turkey’s multiple comparisons test (A-D). P < 0.05 indicates significance.

By tracking KSHV infection and cell-state transitions across tissue niches and disease stages, we observed that KSHV infection of CD34⁺LECs clearly preceded that CD34⁻LECs (Figure 5E, left panel; Figure S7). In patch lesions, KSHV⁺LECs were exclusively CD34^+^, while CD34⁻LECs remained essentially virus-free, reinforcing the idea that CD34⁺ cells are preferentially infected and likely serve as the tumor-initiating population (Figure 5E, left panel; Figure S7). With tumor progression, the majority of infection density gains arose from the CD34⁺ compartment, with negligible contributions from CD34^-^LECs. Infected CD34⁺LECs progressively accumulated along pseudotime and localized to Tumor-associated niches, while uninfected CD34⁺LECs and CD34⁻LECs persisted without expansion (Figure 5E; Figure S7), highlighting the selective expansion and spatial dominance of KSHV⁺CD34⁺LECs. Immunofluorescence imaging validated these findings, confirming abundant LANA⁺ spindle-shaped cells that co-expressed CD34 and LYVE1 (Figure 5F). Together, these results supported a model in which KSHV selectively infects CD34⁺LECs, transforming them into a clonally expanding, tumor-propagating population.

We further examined whether other cell types, including VECs, fibroblasts, and macrophages, acquired the CD34 marker upon KSHV infection, and how KSHV⁺LECs influenced viral dissemination across these lineages and reshaped surrounding niches. A substantial proportion of these cells also expressed CD34, both in uninfected and KSHV-infected cells (Figure 5E; Figure S7 and S8). However, CD34⁺LECs preceded other infected cells, followed by CD34⁺ and CD34⁻ VECs, fibroblasts, and macrophages (Figure 5E, left panel). In patch lesions, similar to CD34⁻LECs, both CD34⁺ and CD34⁻ VECs, fibroblasts and macrophages remained almost virus-free, reaffirming that infection of CD34⁺LECs were the tumor-initiating population. Space-time density heatmaps of infected cells confirmed LECs as the dominant infected population across all stages with secondary infection waves emerging in VECs, fibroblasts, and macrophages. Interestingly, KSHV⁺CD34⁺ macrophages and KSHV⁺CD34⁺ fibroblasts were predominantly localized in the Tumor-associated niches, suggested their direct impact on tumor progression, while KSHV⁺CD34⁻ macrophages, KSHV⁺CD34⁻ fibroblasts, and KSHV⁺VECs were primarily localized within Stroma niches (Figure 5E, left panel; Figure S7). As expected, both CD34⁺ and CD34⁻ uninfected VECs, fibroblasts, and macrophages were located in Stroma niches (Figure 5E, right panel; Figure S7). Scatterplots further showed that infected cell densities increased markedly in Tumor-associated niches, predominantly driven by LECs (slope = 45.9 cells/μm^2^/τ, which denotes increase in infected cells per μm^2^ across KS progression from control to nodular), while only smaller increases were observed in VECs (2.55 cells/μm^2^/τ), fibroblasts (2.55 cells/μm^2^/τ), and macrophages (5.1 cells/μm^2^/τ), which were primarily CD34⁺ cells (Figure 5G). Moderate increases were observed in Vascular-associated (LEC slope = 10.2 cells/μm^2^/τ) and Immune-associated niches (LEC slope = 10.2 cells/μm^2^/τ), confirming secondary dissemination from expanding LEC populations. In contrast, skin-associated niches showed minimal to no increase across all cell types, highlighting niche-specific constraints on viral spread. These results confirm that KSHV⁺LECs serve as local viral amplifiers, selectively remodeling surrounding niches via spatially restricted viral dissemination. In parallel, density maps of uninfected cells revealed niche-level reorganization in response to the expansion of KSHV⁺LECs (Figure 5E, right panel; Figure S7). KSHV⁻LECs persist alongside their infected counterparts within Tumor-associated niches, while uninfected VECs and fibroblasts shifted outward toward peri-vascular and dermal stromal regions, and uninfected macrophages were localized within Immune-associated niches. These findings reaffirm that KSHV⁺LECs not only drive clonal expansion but also reshape the TME, facilitating secondary spread to adjacent stromal lineages, and creating infection-permissive tissue compartments that support sustained tumor growth.

### Integrative Spatial Analysis Reveals Neoangiogenesis as a Driver of KS Progression

Neovascularization is a hallmark of KS, playing a critical role in tumor progression and viral propagation.^45^ To systematically examine the formation and expansion of new blood vessels in KS lesions, we integrated vessel segmentation, immunofluorescence validation, and gene expression profiling across different tumor stages and spatial niches.

We developed an algorithm to identify and segment blood vessels (Figure 6A). This process involves detecting VECs, PCs, and luminal areas, followed by density-based clustering to demarcate individual vessels (see Methods). Our approach achieves an overall Dice score of 0.93 against expert pathologist annotations, indicating robust and accurate vessel segmentation. By overlaying expert pathologist annotations (white) with immunofluorescence staining images for CD31 (green) and DAPI (blue), we validated the accuracy of the vessel detection algorithm (Figure 6B, left panel).

**Figure 6.**
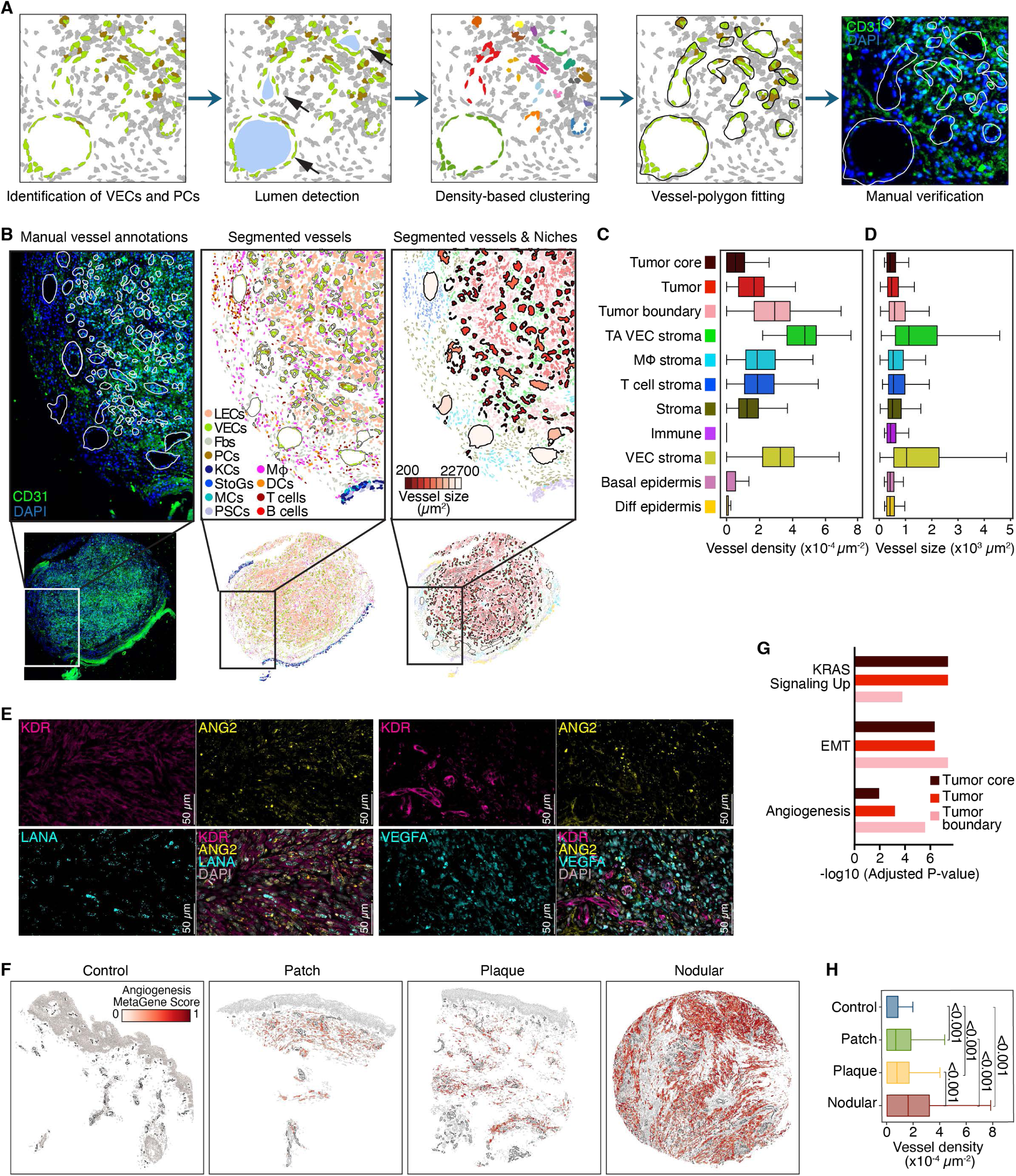
Neoangiogenesis as a driver of Kaposi’s sarcoma (KS) progression revealed by integrative spatial analysis. (A) Pipeline for vessel segmentation, which achieved high segmentation accuracy (Dice score = 0.93). (B) Manual annotation of vessels (white) overlying on CD31 (green) and DAPI (blue) staining images (left) and segmented vessels mapped to cell types and niches (middle, right), showing good agreement between segmented vessels and manual annotation and revealing dense vasculature near tumor boundaries, particularly within LEC-rich regions. (C) Quantification of vessel density across KS niches showing highest densities in Tumor core and Tumor boundary regions, with moderate densities in Immune-enriched and distal Stromal areas. Bars represent median vessel densities with interquartile ranges (top 75% and lower 25%) and extended whiskers were created using Tukey method. (D) Quantification of vessel size (vessel cross-sectional area) across KS niches showing small-lumen capillaries in the Tumor core gradually enlarging toward Tumor-associated stroma and boundary, indicating progressive vessel maturation. Bars represent median vessel sizes with interquartile ranges (top 75% and lower 25%) and extended whiskers were created using Tukey method. (E) Representative immunofluorescence staining images of KDR (magenta), ANG2 (yellow), and nuclei (grey) together with LANA (cyan, left) or VEGFA (cyan, right) in a nodular KS lesion. (F) Spatial maps of angiogenesis meta-gene expression across KS stages showing localized high-expression zones in patch lesions that expand and interconnect in plaque lesions, reaching widespread activation in nodular lesions. (G) Differentially expressed genes in cells within or contacting Tumor-associated vessels are enriched for Angiogenesis, KRAS signaling, EMT, and other cancer-related pathways, highlighting active vascular remodeling in KS. (H) Quantification of vessel density across KS stages shows moderate increases in patch and plaque lesions, with nodular lesions exhibiting markedly higher vessel densities. Bars represent median vessel densities and standard deviations.

Furthermore, we overlaid segmented vessels with annotated cell types and niches (Figure 6B, middle and right panels), revealing that regions near the Tumor boundary consistently contain high concentrations of blood vessels, particularly within LEC-rich areas, compared to nearby stromal regions. This clustering of endothelial cells with pronounced CD31 staining underscores the abundant vasculature that defines KS tumors.

To quantify the spatial features of the identified vasculature, we analyzed changes in vessel density and size across the niche gradient. Vessel density was calculated as the number of vessels within a defined hexagonal area (42 μm on each side) and normalizing it to a hexagonal area of 4,582.69 μm². Vessel size was determined by calculating the cross-sectional area of the polygon enclosing the vessel. These quantitative measurements revealed that the Tumor core and immediate Tumor boundary regions exhibited higher vessel densities compared to more distal stromal or epidermal areas (Figure 6C). Vessel density peaked in the Tumor-associated VEC stroma and Tumor boundary regions, indicating hotspots of active endothelial proliferation and remodeling, likely driven by both VEC activation and infected tumor cells, while the Immune-enriched niches surrounding the tumor regions showed moderate vessel densities (Figure 6C). This spatial density profile mirrored the increasing infection gradient towards the Tumor core (Figure 3F). Interestingly, the Tumor core harbors small-lumen capillaries, which gradually transition into larger vessels as they move outward into the Tumor-associated stroma (Figure 6D). These increasing lumen cross-sectional areas likely reflect progressive vessel maturation and remodeling, with smaller capillaries coalescing into larger vascular channels. This outward gradient suggests that initial angiogenic sprouting and smaller vessel formation occurs within Tumor niches, driven by autocrine and paracrine factors such as ANG2, VEGF, and vIL6 secreted by tumor cells, while vessels at the Tumor boundary become more mature and expanded. Immune-enriched and general stroma areas showed smaller vessel diameters, likely reflecting either earlier stages of angiogenic sprouting or narrower capillary beds (Figure 6D). As expected, epidermal compartments (basal or differentiated layers) showed the smallest lumen areas, consistent with the sparse vascular supply in normal or less-transformed epidermal layers.

To confirm the molecular drivers of these vascular changes, we performed immunofluorescence staining for ANG2, KDR and LANA as well as ANG2, KDR and VEGFA (Figure 6E). The expression of ANG2, a marker for neoangiogenesis, and VEGFA, a marker for blood vessel sprouting and stability, were closely associated with KDR-positive endothelial cells, consistent with active vessel remodeling. The proximity of LANA⁺ spindle-shaped tumor cells to ANG2⁺ and KDR⁺ vascular regions further highlights the role of KSHV in shaping a proangiogenic TME.^46–48^ This spatial juxtaposition reinforces the role of viral infection in driving neoangiogenesis and vasculature remodeling in KS tumor development.

To assess whether these angiogenic changes intensify with tumor progression, we analyzed an angiogenesis hallmark meta-gene signature across control, patch, plaque, and nodular lesions (Figure 6F). We defined the metagene based on the average expression of the MSigDB Angiogenesis Hallmark Genes COL5A2, PDGFA, ITGAV, and POSTN, excluding the cellular marker genes and KSHV genes (see Methods). Early patch lesions showed spatially scattered and localized high-expression zones, which became more extensive and interconnected in plaque lesions (Figure 6F). By the nodular stage, nearly the entire lesion displayed elevated expression of angiogenic genes, reflecting the growing demand for vascular proliferation and remodeling to support rapid tumor expansion. This spatial signature aligned with immunofluorescence staining results (Figure 6E) and vessel detection results (Figure 6A-6D), pointing to a TME that becomes progressively more vascularized and angiogenic as KS advances from early patch to nodular lesions.

Furthermore, we identified differentially expressed genes in cells within or contacting the identified vessels in a niche versus the rest of the niches. The top differentially expressed genes in the Tumor niches were enriched by the Angiogenesis pathway in the MSigDB Cancer Hallmark (Figure 6G), along with pathways such as KRAS Signaling Up, EMT and several other pathways (Figure S9). The vessels associated with Tumor niches showed enriched pathways such as G2-M Checkpoint, E2F Targets, Mitotic Spindle, and EMT, while the normal vessels were only enriched in the EMT and Myogenesis Pathways (Figure S9). These results reinforce the trend of angiogenesis across niche gradients and tumor stages, particularly highlighting the direct correlation between high vessel density and advanced KS lesions (Figure 6H).

Specifically, patch and plaque lesions showed moderately increased vascularization compared to control tissues, whereas nodular lesions exhibited strikingly higher vessel densities. This trend suggests that as KS progresses, the tumor increasingly relies on newly formed blood vessel networks to meet its demands for nutrients, enhanced metabolism, oxygen consumption, waste removal, and potentially the dissemination of tumor cells and virus.

### KSHV Infection Reprograms Macrophages and Shapes Their Spatial Organization in the Tumor Proximity and Microenvironment during KS Progression

Building on our observation of macrophage expansion and T cell decline during KS progression (Figure 1F), we investigated the role of macrophages in the TME throughout KS progression. We analyzed macrophage abundance, spatial organization, infection status, and transcriptional program across tumor stages and tissue niches.

Among immune populations, macrophages were consistently the most enriched in tumor-adjacent niches, including the Tumor boundary, Tumor-associated VEC stroma, and Macrophage immune stroma, suggesting niche-specific recruitment (Figure 7A).

**Figure 7.**
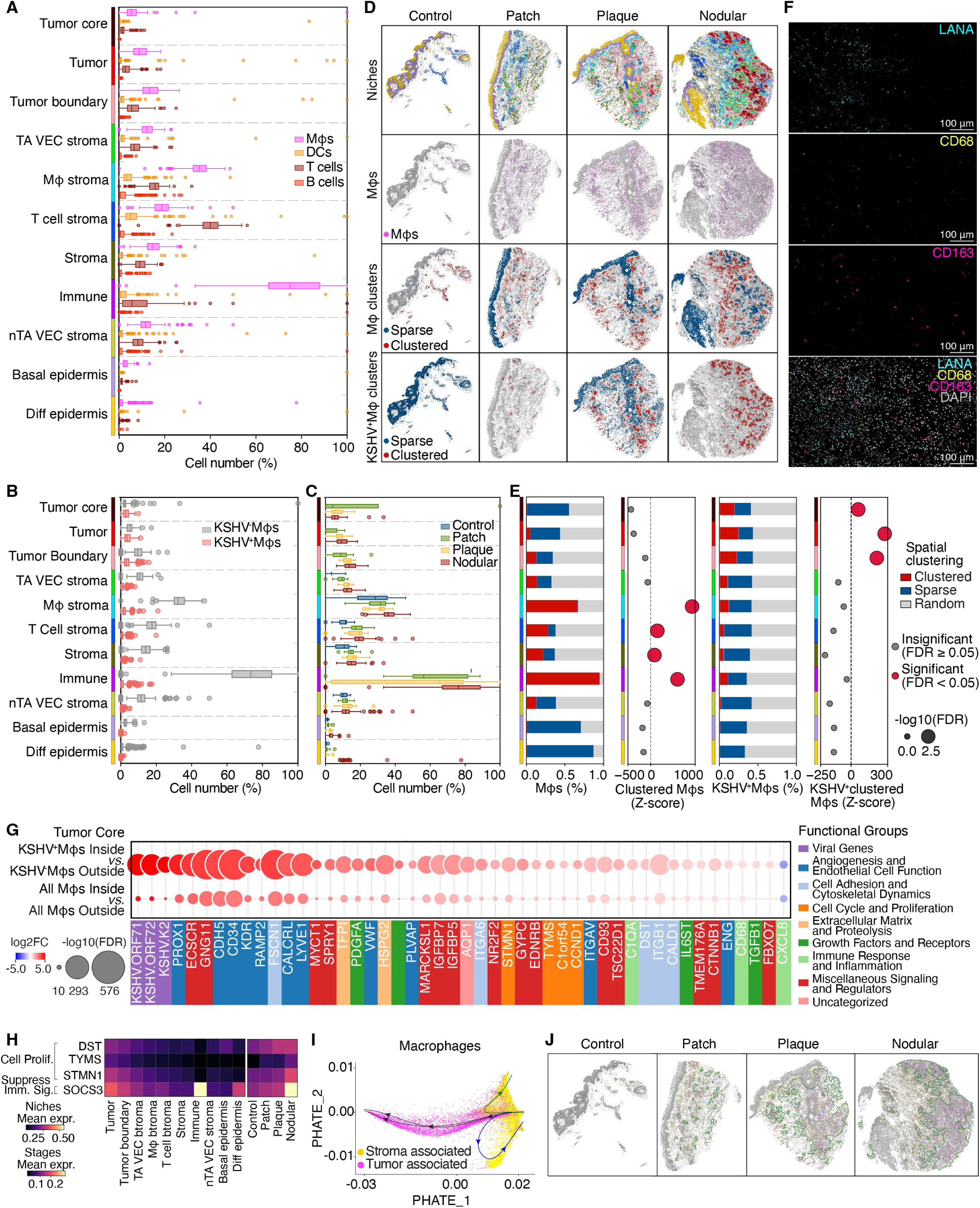
KSHV-driven macrophage reprograming and spatial reorganization in the tumor proximity microenvironment during Kaposi’s sarcoma (KS) progression. (A) Box-and-whiskers plots of major immune cells, including macrophages (Mφs), T cells, B cells, and dendritic cells (DCs) across different niches, showing enrichment of macrophages in tumor-proximal niches, including Tumor boundary, Tumor-associated vascular endothelial cell (VEC) stroma, and Macrophage immune stroma. Bars represent median percentages with interquartile ranges (top 75% and lower 25%) and extended whiskers were created using Tukey method. (B) Box-and-whiskers plots of KSHV-infected and -uninfected macrophages, showing that most macrophages remain uninfected across niches, while KSHV⁺ macrophages are largely restricted to Tumor and Tumor boundary compartments. Bars represent median percentages with interquartile ranges (top 75% and lower 25%) and extended whiskers were created using Tukey method. (C) Box-and-whiskers plots of macrophages by KS stage, showing that KSHV⁺ macrophages are largely restricted to Tumor and Tumor boundary compartments. Bars represent median percentages with interquartile ranges (top 75% and lower 25%) and extended whiskers were created using Tukey method. (D) Spatial maps of macrophages across KS stages showing two organizational patterns: dense clusters in stromal and immune compartments and sparse cells along tumor margins. KSHV⁺ macrophages form tight clusters in Tumor and Tumor boundary regions. (E) Quantification of macrophage spatial organization using Moran’s I and permutation Z-scores. Clustered macrophages are enriched in Immune and Macrophage stroma niches, while sparse KSHV⁺ macrophages are enriched within Tumor core and Tumor boundary regions. (F) Representative immunofluorescence staining images of LANA (cyan), CD68 (yellow), CD163 (magenta), and nuclei (grey) in a nodular KS lesion. (G) Differential gene expression in macrophages inside versus outside tumor compartments, stratified by KSHV infection. Tumor-localized macrophages upregulate lymphatic- and vascular-associated genes, while KSHV⁺ macrophages show broader reprogramming, including viral, proliferation, inflammatory, and matrix-remodeling genes. (H) Mean expressions of immune suppression and proliferation markers across niches and KS stages, demonstrating a progressive increase towards tumor niches and with advancing KS progression. (I) PHATE projection of macrophages scored by a tumor-infiltrating metagene, highlighting two transcriptional trajectories: Stromal-associated macrophages (SAMs) distributed broadly, and Tumor-associated macrophages (TAMs) localized to tumor compartments. (J) Spatial maps of SAMs and TAMs across KS stages showing a progressive shift from Stromal-associated SAMs in early lesions to dense, tumor-localized TAM domains in nodular lesions.

Despite their abundance, most macrophages remained uninfected (Figure 7B). KSHV⁺ macrophages were largely restricted to the Tumor and Tumor boundary compartments, with negligible presence in stromal or immune niches. This suggests that macrophage accumulation is driven by niche-specific cues, while KSHV infection selectively marks a small subset localized within tumors, highlighting the importance of both spatial location and infection status in defining distinct functional macrophage states in the TME.

To investigate whether spatial localization dictates macrophage functional heterogeneity, we examined their spatial distribution across KS stages. In control and patch tissues, macrophages were broadly dispersed throughout the dermis, but as the lesions advanced, they progressively concentrated around tumor compartments, often extending into perivascular regions (Figure 7C). Spatial plots revealed two distinct organizational patterns: macrophages either formed dense clusters in stromal and immune compartments or were distributed as sparse, scattered cells along tumor margins (Figure 7D, second row). Moran’s I autocorrelation analysis (see Methods) classified macrophages as either clustered or sparse (Figure 7D, third row and 7E, first column). In plaque and nodular lesions, clustered macrophages (red) dominated dermal and perivascular zones, especially in the Macrophage stroma, Tumor-associated VEC stroma, and Immune-enriched niches, whereas sparse macrophages (blue) were enriched within tumor compartments (Figure 7D, third row). This spatial dichotomy became more pronounced in KSHV⁺ macrophages, which formed tight microclusters confined to the Tumor core and Tumor boundary, with negligible presence in peripheral niches (Figure 7D, bottom row and 7E, third column). Permutation-derived Z-scores confirmed this structure: clustered macrophages were significantly enriched in Immune (Z ≈ 1000, P<0.05) and macrophage stroma (Z ≈ 900, P<0.05), while sparse KSHV⁺ macrophages were specifically enriched in the Tumor core (Z ≈ 300, P<0.05) and Tumor (Z ≈ 250, P<0.05) regions (Figure 7E, second and fourth columns).

Immunofluorescence imaging confirmed the infiltration of KSHV⁺ macrophages in Tumor core and Tumor regions (Figure 7F). These results reveal two spatially and likely functionally distinct macrophage populations in KS tumors: (1) uninfected, clustered macrophages concentrated in stromal and immune zones, and (2) infected, sparse macrophages infiltrated into tumor compartments. This spatial compartmentalization suggests distinct transcriptional programs between the two macrophage subsets.

We performed differential expression analysis comparing macrophages inside and outside tumor compartments, stratified by infection status. Even in the absence of infection, tumor-localized macrophages had upregulated expression of lymphatic and vascular-associated genes, including PROX1, CD34, LYVE1, and RAMP2, indicating that the TME in tumor proximity alone was sufficient to induce partial reprogramming of macrophages toward a lymphatic-progenitor-angiogenic state (Figure 7G, bottom row). However, KSHV⁺ macrophages displayed broader and more pronounced transcriptional remodeling (Figure 7G, top row). Viral genes such as ORF71, ORF72, and vIL6 were strongly upregulated (log₂FC > 4), alongside proliferation markers STMN1, TYMS, and CCND1; inflammatory mediators CXCL8, C1QA, and CD68; and matrix remodeling factors DST, ITGB1, and CALD1 (Figure 7G, top row, 7H). Notably, KSHV⁺ macrophages also upregulated TGFB1, a canonical immunosuppressive cytokine, alongside CXCL8 and IL6ST, which are typically associated with proinflammatory signaling. Furthermore, we observed stage and niche specific upregulation of SOCS3, an immune suppression marker, as well as DST, TYMS, and STMN1, proliferation-associated genes, with expression progressively increasing towards tumor niches and as disease progresses (Figure 7H). This combination suggests that KSHV⁺ macrophages adopt a transcriptionally inflamed but functionally immunosuppressive hybrid phenotype. Rather than mounting effective anti-tumor immunity, these macrophages likely contribute to immune evasion, stromal remodeling, and angiogenesis, consistent with the behavior of tumor-associated macrophages (TAMs) described in virally driven tumors.^49,50^ These findings support a two-step model: first, tumor proximity TME primes macrophages toward a lymphatic-progenitor-angiogenic state; and second, KSHV infection amplifies this transition, producing a highly reprogrammed, tumor-permissive macrophage state with both inflammation and immune suppression features.

To define the transcriptional states underlying this spatial and infection-linked macrophage bifurcation, we computed a metagene score based on the top 20 genes upregulated in tumor-infiltrating macrophages from our differential expression analysis (Figure 7G). Projecting this score onto the same PHATE manifold as in Figure 4J revealed a clear split within the macrophage population into two distinct transcriptional trajectories (Figure 7I). One trajectory contains macrophages with low metagene scores, corresponding to those in peripheral stromal and immune niches. The other trajectory comprised macrophages with high scores, expressing genes linked to angiogenesis, proliferation, and immune activation. Based on their spatial localization and gene expression programs, we termed these two states Stromal-associated macrophages (SAMs) and TAMs, respectively. SAMs are broadly distributed, consistently uninfected, and maintain a transcriptional program aligned with immune and dermal identity. In contrast, TAMs are localized to tumor compartments, frequently infected, and exhibit a reprogrammed transcriptional state indicative of tumor-permissive activity, including angiogenesis, matrix remodeling and functional immunosuppression, despite the expression of inflammatory mediators. Spatial mapping of these states across tissue sections revealed a stage-dependent transformation (Figure 7J). In patch lesions, SAMs predominated throughout both tumor and adjacent zones, while TAMs were rare. In contrast, TAMs became abundant in nodular lesions, forming a dense, spatially coherent domain within the Tumor core and Tumor boundary niches, displacing the earlier stromal distribution of SAMs. These findings demonstrate that KS tumors orchestrate a hierarchical, spatially structured transformation of macrophages, where tumor proximity initiates partial reprogramming and KSHV infection drives full activation of a hybrid transcriptional state that supports tumor growth, vascular remodeling, and immune evasion.

### Spatial Single-Cell Features Accurately Predict KS Progression and Reveal Stage-Informative Cellular Programs

To systematically characterize the spatial cellular and molecular determinants of KS progression, and assess their predictive power, we derived three classes of gene expression features from scSRT analyses: (1) bulk features, representing aggregate gene expression across all tissue cores; (2) single-cell features, preserving cell-type-specific expression without spatial context; and (3) spatial single-cell features, which capture gene expression of individual cell types within defined tissue niches, offering a microenvironment-specific view of transcriptional changes linked to disease stage.

We trained three classifiers (LASSO, Ridge, and Extreme Gradient Boosting, XGBoost) on each feature class and evaluated their ability to discriminate patch, plaque, and nodular lesions (see Methods). Models based on spatial single-cell features consistently outperformed those based on single-cell or bulk features, achieving the highest ROC AUC scores across models (Figure 8A). The improvement observed in linear models (LASSO, Ridge) highlights the added value of spatial resolution even in simpler frameworks. In contrast, the performance gap narrowed in XGBoost, suggesting that non-linear models can partially recover spatial dependencies but still benefit from explicit niche-specific encoding. Together, these findings underscore the importance of spatial context in capturing the biology of KS progression.

**Figure 8.**
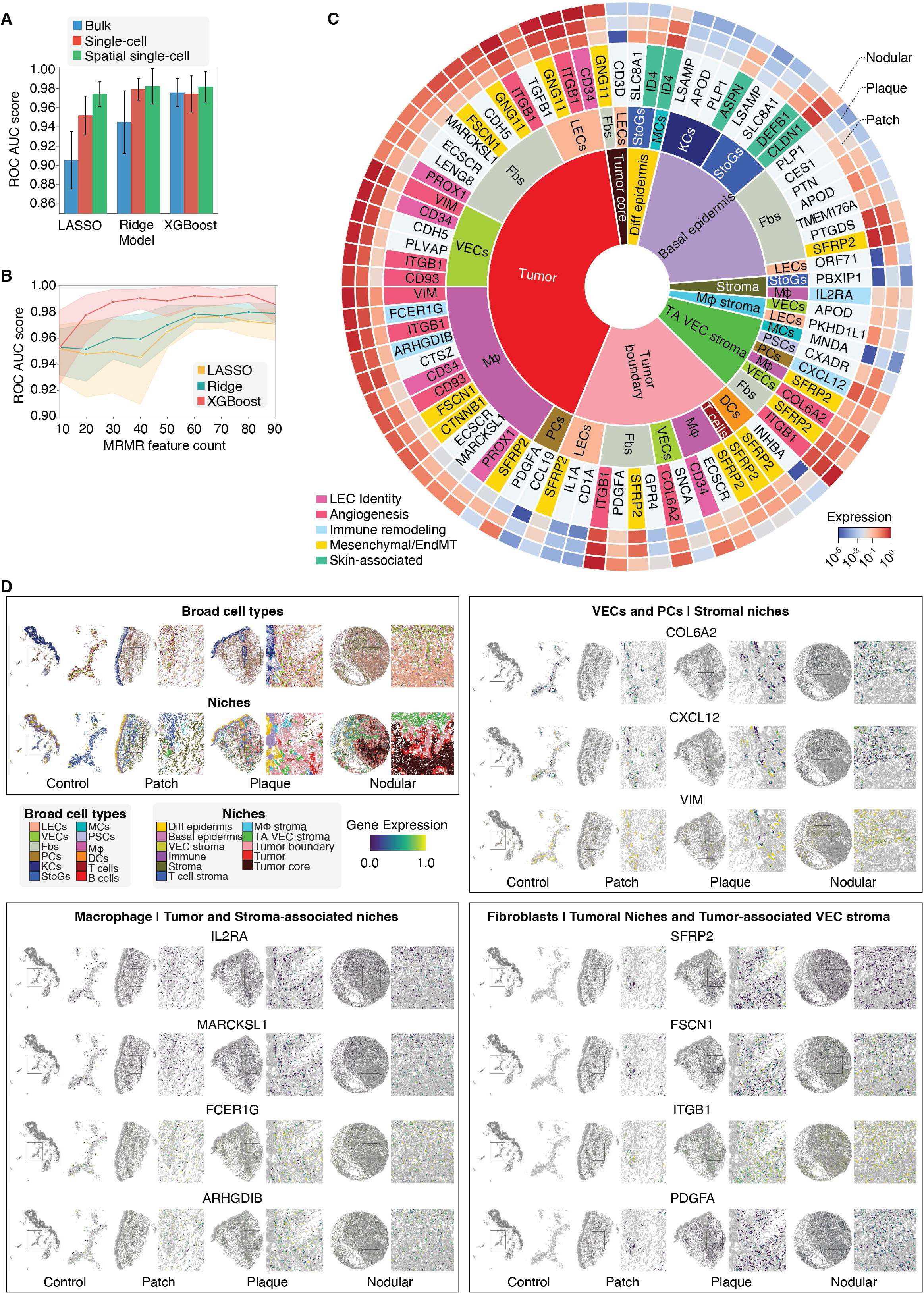
Spatial single-cell features accurately predict Kaposi’s sarcoma (KS) progression and reveal stage-specific cellular programs. (A) ROC AUC comparison of classifiers trained on bulk, single-cell, and spatial single-cell features, showing superior predictive performance of spatially resolved features for discriminating patch, plaque, and nodular KS lesions. Bars represent median scores and standard deviations. (B) Feature selection using MRMR reveals the top spatial single-cell features driving stage prediction, with ROC AUC plateauing at ∼80 features, highlighting niche- and cell type-specific programs underlying KS progression. (C) Expression of top 80 spatial single cell features (genes of a cell type within a niche), highlighting KSHV-driven reprogramming of lymphatic endothelial cells (LECs), fibroblasts, and Tumor-adjacent stromal cells, with upregulation of lymphatic, progenitor, and tumor-supportive programs along disease progression. (D) Spatial maps of selected top features, showing stage-specific macrophage, vascular, and matrix features with polarization toward immunosuppressive, proangiogenic, and endothelial-to-mesenchymal states, reflecting niche-specific contributions to tumor progression and TME remodeling.

To identify the most informative features for stage prediction, we applied the Minimum Redundancy Maximum Relevance (MRMR) algorithm to select the top spatial single-cell features across three models.^51^ ROC AUC improved steadily with features count, plateauing at 0.99 around 80 features, which were selected to balance performance and interpretability (Figure 8B). These features, derived from the best-performing XGBoost model, revealed a spatial coordinated program of niche remodeling, immune suppression, and endothelial plasticity across KS stages (Figure 8C; Table S4). The most predictive signals originated from Tumor-associated niches, including Tumor core, Tumor, and Tumor boundary, with key contributions from LECs, fibroblasts, macrophages, and VECs (Figure 8C). Notable stage-informative genes included PROX1, ITGB1, SFRP2, CD34, CD93, and FSCN1, which are involved in lymphatic specification, angiogenesis, matrix remodeling, and tumor invasion.

Consistent with earlier findings, LECs were the most informative and reprogrammed population, expressing viral genes (e.g., ORF71), inflammatory cytokines (e.g., IL1A), and LEC markers (e.g., CD1A and PKHD1L1), associated with tumor progression (Figure 8C; Table S4). These features reflect the dual roles of LECs as primary KSHV targets and tumor-initiating spindle cells. Importantly, LEC-associated genes like PROX1 and progenitor genes such as CD34 were also upregulated in infected macrophages, fibroblasts, and VECs located in tumor-adjacent niches, reinforcing our observations of KSHV-driven cross-lineage reprogramming toward a tumor-supportive mixed lymphatic- and progenitor-like identity (Figure 8C; Table S4).

For instance, spatial plots show high expression of GNG11 and ITGB1 in LECs within Tumor core and Tumor, respectively, and expression of ARHGDIB and FSCN1 in infected macrophages and fibroblasts, respectively (Figure S10).

Macrophages also contributed strongly to the predictive signature. Enriched genes, including ARHGDIB, CD93, IL2RA, MARCKSL1, and FCER1G and reflect their polarization toward an immunosuppressive and proangiogenic state, particularly in plaque and nodular lesions (Figure 8D, bottom left box; Table S4). Stromal VECs and PCs upregulated COL6A2, CXCL12, and VIM, markers of endothelial-to-mesenchymal transition and vessel remodeling (Figure 8D, top right box). In contrast, skin-associated niches, such as the Basal and Differentiated epidermis, were enriched for early-stage epithelial markers like DEFB1, CLDN1, ID4, and ASPN, which progressively declined from patch to nodular stages, reflecting skin barrier breakdown and tumor infiltration (Figure S10; Table S4). Fibroblasts within tumor-adjacent and stromal niches exhibited a distinct transcriptional program consistent with cancer-associated fibroblasts (CAFs). Notably, SFRP2, a secreted Wnt modulator and hallmark CAF marker, was among the most stage-informative features, expressed prominently in fibroblasts, macrophages, and VECs near tumor regions (Figure 8D, bottom right box). Other fibroblast markers such as FSCN1, ITGB1, and PDGFA were enriched in tumor and perivascular compartments, indicating their supporting roles in matrix remodeling, cell migration, and angiogenesis (Figure 8D, bottom right box). These patterns highlight a tumor-associated fibroblast program that co-opts mesenchymal and vascular functions to reinforce tumor growth.

Although vIL6 spatial features did not rank among the top 80 MRMR predictors, their highest-ranking instances were all restricted to tumor niches (Tumor LECs: 3381, Tumor fibroblasts: 3657, Tumor boundary LECs: 4018, Tumor core LECs: 4220, and Tumor boundary fibroblasts: 4413), suggesting their lower ranks likely reflect redundancy with top ranked features. This observation is confirmed by correlation analysis, showing that vIL6 features were strongly associated with top 80 stromal and vascular features, including COL6A2 in Tumor boundary and Tumor-associated stroma VECs (Pearson correlation r = 0.39-0.44 across niches), GPR4 in Tumor boundary fibroblasts (r = 0.41-0.43), PDGFA in Tumor pericytes (r = 0.43), and CD34 in Tumor macrophages (r = 0.39) (Figure S11). These associations reinforce our earlier findings that vIL6 is a central driver of KS angiogenesis, inflammation, and niche remodeling and further suggesting that vIL6 acts a viral amplifier by enhancing fibroblast activation, extracellular matrix and vessel remodeling, and progenitor-like reprogramming within TME through paracrine viral cytokine signaling.

Together, these results demonstrate that spatial single-cell features not only improve stage prediction but also uncover biologically meaningful patterns of cellular reprogramming, niche remodeling, and transcriptional transitions that drive KS pathogenesis. To facilitate broader use of these findings, we have provided the top-performing XGBoost model based on spatial single-cell features as a public resource for accurate KS stage prediction, enabling streamlined application and further extension of our work.

## Discussion

KS, a highly inflammatory and angiogenic tumor caused by KSHV infection, presents significant challenges in clinical management and therapy.^52^ The multifactorial nature of KS involves a complex interplay between viral infection, cellular reprogramming, immune modulation, and TME remodeling.^3^ Our study, representing the first single-cell spatial transcriptomic analysis of KS tumors to date, not only expands our understanding of KS pathogenesis but also uncovers pivotal mechanisms that drive tumor initiation, progression, and immune evasion. By analyzing 43 KS patients across different tumor stages with 256 tissue cores, we reveal critical findings that can transform how we view KS biology and suggest novel therapeutic strategies.

A key discovery of our study is the broad KSHV tropism, which infects a diverse range of cell types beyond the traditionally recognized LECs and LECs. We show that KSHV infects LECs, VECs, fibroblasts, and macrophages, with these infected cells acquiring hybrid identities that combine endothelial, mesenchymal, and immune signatures. These results of transcriptional reprogramming recapitulate the hallmark of KS progression, which are the co-expression of cellular markers in tumor cells typically found in both endothelial and mesenchymal lineages, indicative of a mixed cellular identity that supports tumor growth and immune evasion.^41–43^

The extent to which KSHV induces endothelial plasticity is remarkable, as infected cells not only maintain their vascular identity but also gain properties associated with progenitor/mesenchymal cells, facilitating angiogenesis and immune modulation. This endothelial-to-mesenchymal transition (EndMT) underscores the dynamic reprogramming that KSHV drives within the TME, allowing the virus to orchestrate a supportive environment for tumor development. As a result, one of the most striking findings is the KSHV induction of a profound endothelial plasticity that contributes to both vascular remodeling and niche reorganization. KSHV-infected endothelial cells, particularly in LECs and VECs, exhibit a transition toward hybrid endothelial-mesenchymal phenotypes, characterized by upregulation of both endothelial and mesenchymal markers. This transition is critical for the formation of new blood vessels in the tumor, a process known as neoangiogenesis, which is essential for sustaining tumor growth and facilitating viral propagation.^45^

Our results show that KSHV infection drives the expression of key angiogenic markers such as PROX1, LYVE1, and KDR, indicating that infected LECs play a central role in lymphatic remodeling and tumor vasculature formation. Beyond KSHV-induced vascular remodeling, spatial analysis uncovered dynamic but discrete niches in KS lesions along disease progression. Tumor-associated niches including the Tumor core, Tumor, Tumor boundary, and the adjacent stroma emerged as dominant sites of infection and clonal expansion, while Immune- and Skin-associated niches exhibited restricted or declining infection. As lesion advanced from patch to nodular stages, these niches were progressively restructured, where infected LECs accumulated, while fibroblasts and VECs shifted into tumor-adjacent compartments, and uninfected immune cells were displaced to peripheral zones. This niche remodeling creates infection-permissive microenvironments that sustain viral persistence, foster angiogenesis, and suppress immune clearance. Together, these findings underscore KSHV infection not only reprograms individual cell types but also orchestrates tissue-level architecture and the broader TME to promote tumor progression and viral spread.

One of the most novel and insightful findings in our study is the identification of CD34⁺LECs as the origin of KS tumor initiation. We show that KSHV preferentially targets CD34⁺LECs, transforming them into tumor-propagating cells that contribute to both the lymphatic and vascular components of KS tumors. This finding significantly changes our understanding of KS tumor initiation, suggesting that CD34⁺ progenitor cells, including those with mesenchymal stem cell-like characteristics, are the primary targets for KSHV infection and tumor initiation. Indeed, previous works have demonstrated that KSHV infected progenitor cells acquire both lymphatic, vascular and mesenchymal markers while maintaining precursor identity.^11,18–20^ Importantly, we observed that CD34⁺LECs remain the dominant infected population throughout disease progression, with a progressive increase in their frequency in advanced stages of KS, further solidifying their role in tumor initiation and clonal expansion.

The persistence of CD34⁺LECs as the major infected population in KS tumors suggests that they not only serve as the initial site of viral infection but also contribute to the amplification of the tumor through clonal expansion. The high expression of CD34 in KSHV⁺LECs, particularly in the nodular lesions, marks them as the tumor-initiating population, which sustains tumor growth through both vascular and immune-modulatory functions. This discovery adds a new dimension to our understanding of KS pathogenesis, indicating that KSHV infection of progenitor cells might be a critical driver of tumorigenesis.

We reveal a critical role of KSHV lytic replication in KS progression. Although latent infection plays a central role in maintaining cell survival and immune evasion,^36^ our analysis shows a high level of lytic activity in KS tumors. A significant proportion of KSHV-positive cells in the tumor express viral lytic genes, including RTA and ORF57, which drive the activation of viral replication. Previous works have also detected KSHV lytic replication and the expression of lytic genes in KS tumors despite at the lower rates.^53^ More importantly, we observed that vIL6 is expressed at high levels in tumor cells, particularly in the Tumor boundary and Tumor core regions. vIL6 is known to induce blood to lymphatic endothelial cell differentiation, and contribute to inflammation and angiogenesis, two key features of KS.^54–58^ The high level of vIL6 expression in the tumor correlates with increased vascularization, suggesting that KSHV-driven lytic replication and the secretion of viral cytokines like vIL6 are pivotal for tumor progression. This finding also reinforces the importance of KSHV’s ability to evade immune response during both latent and lytic replication, driving both inflammatory and angiogenic pathways that support tumor survival.

Our study provides new insights into how KSHV drives macrophage reprogramming, revealing the existence of two distinct macrophage populations in the KS TME. While macrophages were previously detected in KS lesions, their role remains obscure.^59^ We identified TAMs that are spatially and functionally distinct from SAMs.

TAMs, which are predominantly found in Tumor core and Tumor boundary regions, are infected with KSHV and exhibit a transcriptional program associated with inflammation, angiogenesis, and immune suppression. These KSHV⁺TAMs likely promote tumor progression by supporting the tumor vasculature and suppressing effective anti-tumor immunity, thus contributing to the immune evasion seen in KS.

Interestingly, our data suggest that the macrophage polarization toward a pro-tumoral, immunosuppressive phenotype is driven by KSHV infection, which amplifies the expression of immune-suppressive cytokines such as TGFB1 and vIL6, alongside proinflammatory mediators like CXCL8. This reprogramming of macrophages highlights the role of immune modulation in KS, where KSHV infection of immune cells plays a central role in immune suppression, angiogenesis, and tumor progression.

The hallmark of KS is its remarkable remodeling of the TME.^3^ In this study, we provide extensive evidence that KSHV infection drives immune modulation, angiogenesis, and stromal remodeling. These processes are facilitated by the tumor-associated niches that emerge during KS progression. Tumor core, boundary, and adjacent vascular niches are enriched for angiogenic and inflammatory signals, promoting tumor growth and immune evasion. The spatial organization of these niches, and the dynamic shifts in their cellular composition, suggest that niche remodeling is a critical driver of KS progression.

The angiogenic signatures, including ANG2, VEGF, COL5A2, PDGFA, ITGAV, POSTN, and vIL6, we identified are particularly relevant, as they highlight the close relationship between KSHV infection and the formation of new blood vessels within the tumor. These findings underscore the central role of endothelial cells in both the viral life cycle and in tumorigenesis, offering new avenues for therapeutic interventions aimed at targeting the vasculature and restoring immune function within the KS TME.

An important contribution of our study is the identification of spatial single-cell features that can predict disease progression in KS. By integrating spatially resolved transcriptomics with machine learning classifiers, we demonstrated that spatial features, specifically those associated with Tumor-associated niches, are highly predictive of KS progression. This finding is critical because it highlights the importance of the spatial arrangement of cells in understanding tumor progression. Spatial single-cell features, such as the expression of endothelial, mesenchymal, and immune-related genes, correlate with disease stage, and can be used as biomarkers for predicting tumor aggressiveness. This predictive capability provides a powerful tool for understanding KS biology and could lead to the development of more accurate prognostic tools.

Finally, this study reveals key cellular mechanisms and tumor progression pathways that can be targeted for therapeutic intervention in KS. Given the essential roles of endothelial plasticity, macrophage reprogramming, and neoangiogenesis, we propose that targeting KSHV-driven vascular remodeling and immune modulation could offer novel therapeutic strategies for KS. By modulating progenitor cell reprogramming, macrophage polarization, and vascular remodeling, it may be possible to halt or slow down KS progression, particularly in the advanced nodular stages, where tumor burden is high and immune evasion is most pronounced.

## Methods

### Study approval

The University of Pittsburgh Institutional Review Board (IRB) determined that all specimens obtained do not meet the definition of living individuals per the Code of Federal Regulations (45CFR 46.102) and the study was not research involving human subjects as defined by the US Department of Health and Human Services and the US Food and Drug Administration regulations, and therefore waived it of ethical oversight (STUDY20050085).

### Skin tissue samples

A total of 43 anonymized cutaneous Kaposi’s sarcoma lesions and 3 normal skin tissues without any known types of cancer were acquired from Pitt Biospecimen Core. All tissues are repository samples and are blinded without linking to the study subjects. All tissues have been reviewed by a pathologist for TMA coring. Clinicopathologic findings are summarized in Table S1.

### Singe-cell spatially resolved transcriptomics (scSRT)

Xenium Slides & Sample Prep Reagents (P/N 1000460, 10x Genomics), Xenium Human Skin Gene Expression Panel Kit (P/N 1000643, 10x Genomics), Xenium Add-on Custom 1 to 50 Gene Panel (P/N 1000464, 10x Genomics), and Xenium Cell Segmentation Add-on Kit (P/N 1000662, 10x Genomics) were used to prepare the library according to the manufacturer’s instructions. Once the library was prepared, the slides were loaded onto a Xenium Analyzer (10x Genomics) for in-situ barcode sequencing, imaging, and data processing using Xenium Decoding Reagents (P/N 1000461, 10x Genomics).

### Hematoxylin-eosin staining (H&E) and whole slide scanning

H&E was carried out with the same slides used for scSRT using the Hematoxylin & Eosin Stain Kit (H-3502, Vector Laboratories) according to the manufacturer’s instructions. The slides were then scanned with Olympus IX83 Microscope with cellSens Software.

### Immunofluorescence assay

Immunofluorescence assay with paraffin-embedded sections was carried out as previously described.^60^ Primary antibodies to KSHV LANA (1:200, ab4103; Abcam), KSHV vIL-6 (1:200, 251352; Abbiotec), PDPN (1:500, BMS1105; Invitrogen), CoraLite®594-conjugated CD31 (1:200, CL594-11265; Proteintech), p-gp130 (S782) (1:200, PA5-105780; Invitrogen), p-STAT3 (S705) (1:20, 710093; Invitrogen), LYVE1 (1:200, ab33682, Abcam), CD68 (1:200, 14-0688-82; Invitrogen), CD163 (1:200, TA506380; Origene), VEGFR3 (1:100, PA5-16871; Invitrogen), VIM (1:500, sc-6260; Santa Cruz), VIM (1:2000, 80232-1-RR; Proteintech), CD34 (1:200, MA1-10202; Invitrogen), KDR/VEGFR2 (1:200, 26415-1-AP; Proteintech), VEGFA (1:50, ab1316; Abcam), and ANG-2 (1:100, AF623; R&D Systems) were used. Secondary antibodies included Goat anti-Rat IgG (H+L) Highly Cross-Adsorbed Secondary Antibody, Alexa Fluor™ Plus 488 (1:400, A48262; Invitrogen), Goat anti-Rabbit IgG (H+L) Highly Cross-Adsorbed Secondary Antibody, Alexa Fluor™ Plus 555 (1:400, A32732; Invitrogen), and Goat anti-Mouse IgG (H+L) Highly Cross-Adsorbed Secondary Antibody, Alexa Fluor™ Plus 647 (1:400, A32728; Invitrogen). Images were acquired using an Olympus IX83 Microscope with cellSens Software.

Immunofluorescence assay with culture cells was performed as previously described,^61^ using antibodies against LANA (1:500, ab4103; Abcam), VIM (1:500, sc-6260, Santa Cruz), PDPN (1:500, MA5-32617; Invitrogen), and CD34 (1:500, MA1-10202; Invitrogen). Signal was detected with an Alexa555-, or Alexa647-conjugated secondary antibody (Invitrogen). Images were acquired using an Olympus FV2000 Confocal Microscope with cellSens Software.

### Cell culture

Primary human dermal fibroblasts (HDFa) (PCS-201-012; ATCC) were cultured in Fibroblast Basal Medium (PCS-201-030; ATCC) supplied with Fibroblast Growth Kit-Low serum (PCS-201-041; ATCC). KSHV-infected iSLK (iSLK-BAC16) cells were cultured as previously described.^62^

### Virus preparation, titration, and infection

KSHV was prepared from iSLK-BAC16 cells as previously described.^62^ The iSLK-BAC16 cells were replaced with fibroblast medium two days after induction. The supernatant containing KSHV infectious virions was collected and titrated with HDFa. HDFa was infected with KSHV at two infectious units per cell.^61^

### Western-blotting

Western-blotting was performed as previously described,^63^ using primary antibodies against VEGFR3 (1:1000, PA5-16871; Invitrogen), VIM (1:2000, sc-6260; Santa Cruz), PDPN (1:1000, MA5-32617; Invitrogen), and β-actin (1:2000, sc-47778; Santa Cruz).

### TMA core identification

To split individual TMA core and assign the correct pathology and patient metadata to them, we developed an algorithm to automatically identify the TMA cores from the whole-slide tissues based on the identified cell locations using the Density-Based Spatial Clustering of Applications with Noise (DBSCAN) method.^64^ Traditionally, TMA core identification relies on imaging modalities. However, since the variations in experiments can generate different imaging channels based on variable staining requirements and every scSRT datasets generate the spatial locations of individual cells, we opted for a method that directly operates on spatial coordinates of the cells, eliminating dependency on imaging modalities to identify the cores.

We merged spatially proximal clusters based on pairwise distances of cells that falls into a user-defined distance threshold based on the whole-slide image and tissue size. For under-segmented or densely packed clusters, iterative splitting using the DBSCAN algorithm with modified parameters allows improved separation. Together, these functionalities enable tailored refinement of TMA core segmentation, even in complex datasets, with less user inputs or parameter tuning required. After assigning a TMA core identifier to each core in each slide, individual cells were assigned local spatial coordinates for the individual cores.

### Quality control, UMAP, and clustering

To ensure robust and accurate downstream analyses, stringent quality control measures were applied to the scSRT data. Cells exhibiting fewer than 14 detected genes or 15 total counts were filtered out, effectively removing low-quality cells and potential artifacts from subsequent analyses. Following filtering, data normalization was conducted to correct for differences in sequencing depth across cells. This involved scaling total counts per cell to a uniform sequencing depth, followed by log-transformation using a natural logarithmic scale (log1p). This normalization step stabilized the variance across cells and reduced skewness resulting from highly expressed genes.

Next, Principal Component Analysis (PCA) was applied to the normalized expression data to reduce dimensionality and capture the primary variances in the dataset, providing a clearer framework for biological interpretation.^65^ Given the considerable dataset size, computational efficiency was enhanced through the parallelization capabilities offered by Rapids Single-Cell (NVIDIA).^66^ The neighborhood graph construction was subsequently carried out using the Compressed AGgregated nearest neighbor (CAGRA) algorithm, employing squared Euclidean distance metrics, the top 15 principal components, and 50 nearest neighbors.^67^ This approach facilitated efficient and accurate representation of local cellular relationships. Cells were then projected onto a two-dimensional space using Uniform Manifold Approximation and Projection (UMAP) and a Leiden clustering (resolution = 0.5) was carried out to identify 22 initial clusters.^68^

### Cell typing

To achieve precise cell-type annotation from the scSRT data, we adapted a quantitative approach leveraging the receiver operating characteristic area under the curve (ROC AUC).^69^ Marker gene lists specific to known cell types were compiled from the gene panel designed for the experiment (Table S3). For each cell type, meta-gene expression signatures were computed as aggregated marker gene expression values across individual cells.

ROC AUC scores were systematically calculated for every cluster-cell type pair using the meta-gene expressions. Specifically, binary labels representing membership within each cluster were derived, and ROC AUC scores were computed to quantify the correspondence between cluster expression profiles and the predefined cell type signatures. A parallelized computational strategy utilizing the Rapids Single Cell (NVIDIA) was employed to enhance performance across multiple clusters and cell types.

Clusters underwent hierarchical clustering based on their top ROC AUC scores using Euclidean distance and average linkage methods.^70^ A distance threshold, set by a flattening rate of 0.15 (fraction of the maximum dendrogram linkage distance), was applied to merge and consolidate biologically similar clusters. Clusters with maximum ROC AUC scores below 0.6 were annotated as ‘Unknown,’ whereas clusters exceeding 0.7 for multiple cell types were flagged for further sub-clustering. Furthermore, cell-type specificity was enhanced by incorporating mandatory expression of “must-have” marker genes for selected cell types, such as KRT2 for “Spinous to Granular Cells,” requiring a mean expression of at least 1.0 within candidate clusters.

For clusters necessitating additional resolution, recursive sub-clustering was executed with adjusted ROC AUC thresholds (lower threshold: 0.56, upper threshold: 0.7) and a hierarchical clustering flattening rate of 0.25. This was achieved by separating cells from candidate clusters and re-clustering them iteratively. ROC AUC computations and hierarchical merging were recursively applied until each resultant cluster could be uniquely assigned to a single cell type based on stringent ROC AUC and marker gene validation criteria.

### Niche analysis

To define spatial microenvironments, we quantified the local infection-resolved cellular composition by calculating the neighborhood cell type composition (NCTC) for each cell in TMA cores.^71^ Using the derived local spatial coordinates of each TMA core, we identified 30 nearest neighboring cells for each index cell via Euclidean distance. For each neighborhood, we computed the frequency of all surrounding cell types, resulting in a local count vector. Specifically, we stratified the LECs, VECs, fibroblasts, macrophages, dendritic cells, and T-cells by their KSHV infection status producing unique labels for both KSHV⁺ and KSHV⁻ populations. Hence, spatial remodeling and TME changes due to infection can be better resolved, which would otherwise be obscured in aggregated cell type profiles.

The resulting count vectors were clustered using K-Means (K=11) clustering algorithm, defining 11 spatial niches across the tissue landscape.^72^ This choice of 11 clusters was informed based on manual inspection of the spatial niche maps and comparison of their cell type distributions. Each cluster was annotated based on the dominant cell types and their infection status, revealing three niches enriched in KSHV⁺LECs, three Immune-infiltrated zones, and vascular and mostly uninfected stromal compartments, and two skin niches. By naming the most KSHV⁺LEC dominant cluster as the Tumor core and sorting the rest of the clusters based on the average spatial distance of each cell to the nearest cell from the Tumor core niche, we were able to find a niche ordering that further helped identify the intermediate Tumor and Tumor boundary niches. Hence, we were able to identify very high resolution and biologically meaningful spatial domains.

### Moran’s-I analysis of spatial patterns

To quantify spatial autocorrelation and identify spatial clustering of cell types across different niches and tumor proximities in KS tumors, Moran’s I statistic was computed using spatially resolved single-cell data.^71^ Spatial autocorrelation analyses were individually conducted for each cell type as well as bivariate analyses pairing each cell type with specific marker genes (e.g., vIL6).

Spatial coordinates of individual cells were extracted from annotated spatial transcriptomic datasets stored as AnnData objects. These spatial coordinates were converted into a GeoDataFrame using geopandas (v1.0.1). Queen contiguity spatial weights, representing spatial adjacency of cells, were computed using libpysal (v4.13.0) to define neighborhood relationships.^73^ These spatial weights were row-standardized to normalize for the varying neighbor counts across cells. Local Moran’s I statistics were computed using the esda package (v2.7.0). For univariate analyses, Moran_Local was calculated using cell-type-specific abundances. Similarly, Moran_Local_BV was calculated for bivariate analyses to assess spatial relationships between cell-type abundances and gene expression levels.

Cells were classified into significant spatial clusters, spatial outliers, or non-significant, based on a significance threshold (P-value < 0.05) established via conditional permutation testing. Resulting significant spatial clusters were visualized through spatial cluster maps, highlighting spatially dependent cell-type distributions within TME. Stacked bar plots summarizing proportions of each spatial cluster category per niche and tumor proximity were also generated.

### Trajectory analysis

Trajectory analysis was performed to investigate cellular transitions during KS progression. First, endothelial cell populations (LECs and VECs) were identified, and their proportions were calculated per sample and disease stage. TMA cores with a lymphatic-to-vascular endothelial cell ratio less than 5 were selected and categorized into discrete proportion bins, from which 8 balanced representative TMA cores were randomly sampled across KS stages (patch, plaque, and nodular) and 4 TMA cores from control. Subsequently, dimensionality reduction was executed using Potential of Heat-diffusion for Affinity-based Transition Embedding (PHATE) on selected endothelial populations with 20 nearest neighbors (K=20).^74^ Cellular clusters were identified via the Leiden algorithm using a resolution of 0.4 to inform pseudotime inference.^68^

Trajectory inference was conducted using the Slingshot algorithm, applied to the PHATE embeddings with Leiden-derived clusters guiding pseudotime initiation and termination points.^75^ Resulting pseudotime values and trajectory weights were extracted to define cell ordering. Differential gene expression along pseudotime was analyzed using the PyGAM package by fitting Linear Generalized Additive Models (GAMs) on the pseudotime, identifying genes significantly associated with progression trajectories.^76^ Heatmaps visualizing smoothed gene expression of top genes and modules of co-expressed genes were generated to elucidate dynamic molecular changes during KS cellular transitions.

### Infection density estimation and progression trends

To quantify absolute viral burden, we computed infection density as the number of infected cells per unit tissue area (cells per μm^2^) for each core, stratified by lineage and, for LECs, by CD34 status. Core area was estimated from the convex hull of all cellular coordinates in that core. Per-core densities were projected onto the same LEC-derived pseudotime order used above. We then fit linear models of density versus pseudotime using statsmodels.OLS. Regression slopes were reported as cells/μm^2^/τ, with two-sided P-values. Residuals were inspected for approximate normality and homoscedasticity, and fitted lines with 95 percent prediction intervals were overlaid on density scatter plots. This analysis yielded lineage-specific progression rates of infection burden and allowed direct comparison of the steep CD34⁺LEC-driven increase with more modest density gains in fibroblasts, macrophages, and VECs.

### Space-time representation of infection dynamics

We constructed a space-time framework to resolve how KSHV infection changes across niches and clinical stages. Analyses were implemented in Python (v3.11.9) with NumPy (v1.26.4), Pandas (v2.2.2), Matplotlib (v3.9.2), Seaborn (v0.13.2), Statsmodels (v0.14.4), and SciPy (v1.15.3). For each cell, we computed the fraction of neighboring cells that were KSHV infected for predefined lineages such as LECs, VECs, macrophages, and fibroblasts. Neighborhoods were defined as all cells within 10 μm Euclidean distance, converted to pixels using the Xenium resolution of 0.2125 μm per pixel.

Per TMA core, local infection proportions were aggregated by lineage and niche to form niche-by-core matrices scaled to 0 to 100 percent. Cores were ordered along a pseudotime axis defined by the mean density of infected LECs per core, which increases with pathological stage and preserves within-stage variability. Visualization comprised: (i) a niche color stripe, (ii) heatmaps of mean infection percentage for each lineage across cores, and (iii) bar plots of the summed infection percentage per niche. Trend quantification used ordinary least squares (statsmodels.OLS) on per-core lineage means. Cubic smoothing splines (SciPy UnivariateSpline) were applied in cases where noted in the respective figures. We report slopes as percentage points per pseudotime unit (%/τ), with two-sided P-values and 95 percent prediction intervals, and overlay fitted lines on scatter traces.

### Space-time representation for marker expression

We applied the same ordering and niche layout to quantify stage- and niche-resolved marker expression. Curated marker lists were supplied as a table with a column indicating the target lineage for each gene. For a given infection stratum (“infected” or “uninfected”), we divided cells into subsets, intersected the marker list with genes present in the panel, computed per-cell average marker expression by taking the mean across the selected genes, and then averaged by niche and core to generate niche-by-core expression matrices. Heatmaps were rendered with fixed scaling (vmin 0, vmax 0.7, linear scale) and a global colorbar, alongside the niche stripe and a bottom “stage” row derived from the modal stage annotation per core. We produced separate figures for infected and uninfected strata and for each lineage, preserving identical sample order, niche order, fonts, and layout to enable direct visual comparison with infection-percentage plots.

### Vessel segmentation and method validation

Blood vessels are composed of VECs, perivascular cells (e.g. PCs) and a central lumen through which blood flows. We developed an algorithm to segment blood vessels using the scSRT data, which includes four steps: (i) identification of VECs and PCs; (ii) detection of lumen regions; (iii) density-based clustering to localize individual vessels; and (iv) polygon fitting to delineate vessel boundaries.

In the first step, VECs and PCs were identified based on the expression of canonical markers. The scSRT data provided precise spatial coordinates and cell boundaries. Using the shapely package (v2.0.6, https://doi.org/10.5281/zenodo.5597138),^77^ we converted cell boundaries into spatial polygons. Lumen regions were identified by computing the spatial difference between the tissue core boundary and the union of all cellular polygons, followed by morphological operations (erosion and dilation) to eliminate small intercellular gaps. We then retained only those lumen regions that spatially intersected with at least five VECs or PCs.

To ensure consistent representation across vessels of varying sizes, we performed proportional point sampling based on the spatial area of both cellular and lumen regions. Sample points were distributed within each region proportional to its calculated area, enabling uniform downstream analysis. To localize individual vessels, we applied Hierarchical Density-Based Spatial Clustering of Applications with Noise (HDBSCAN; hdbscan v0.8.40; minimum cluster size = 200, minimum samples = 15, alpha = 3) to the combined set of sampled points.^78^ HDBSCAN was chosen for its robustness to variable density and noise, enabling accurate vessel identification as distinct spatial clusters. Finally, we applied concave hull fitting (alphashape v1.3.1) to each cluster to delineate vessel boundary.^79^

Vessel size was determined by calculating the cross-sectional area of the polygon enclosing the vessel (expressed in 10^3^ μm^2^). Vessel density was calculated by counting the number of vessels within a defined hexagonal area (42 μm on each side) and normalizing it to a hexagonal area of 4,582.69 μm².

The segmented vessels were validated against manually annotated vessels. Manual annotations were performed using Xenium Explorer (v3.1.1), where vessels were delineated based on morphological features observed in immunofluorescence images, including spatial arrangements of VECs and PCs. In total, 540 vessels were manually annotated across three tissue cores and used as the reference set for validation. Segmentation accuracy was quantified using the Dice score,^80^ a widely used metric for spatial overlap. It is defined as:

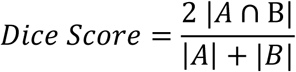

where A is the set of pixels in the predicted vessel and B is the set of pixels in the manually annotated vessel. The score ranges from 0 (no overlap) to 1 (perfect overlap).

### Differential expression analysis

Differential expression (DE) analyses were performed using Scanpy (v1.11.2) in Python (v3.11.9). For each comparison, up to 2,000 cells per group were randomly subsampled to control for group imbalance and computational cost. DE testing was repeated across 2-5 independent folds, and results were aggregated by computing the median log2 fold-change and interquartile range for each gene. KSHV genes were excluded from selected analyses to prevent bias. Visualization of DE results, including volcano plots and scatter maps, was performed using Seaborn (v0.13.2) and Matplotlib (v3.9.2). Effect size was encoded by log2 fold-change in a red-blue colormap, and significance by point size proportional to -log10 adjusted P-values.

### Model training and feature selection for KS stage prediction

We derived three classes of features from the scSRT data to characterize molecular determinants of KS progression: (i) bulk features, defined as aggregate gene expression across entire tissue cores; (ii) single-cell features, representing cell-type– specific expression averaged across all cells; and (iii) spatial single-cell features, capturing gene expression profiles of individual cell types within defined tissue niches.

To assess the predictive power of each feature class for KS stage classification, we trained three machine learning models: LASSO logistic regression, Ridge logistic regression, and XGBoost. All models were implemented using scikit-learn (v1.6.1) and XGBoost (v3.0.0) libraries.^81^ Prior to training, gene expression values were log-normalized. To control for class imbalance across stages, sample weights were applied during training. Model performance was evaluated using ROC AUC in a one-*vs*-rest framework, with five-fold cross-validation.

To identify the most stage-informative spatial single-cell features, we applied the MRMR algorithm (using mrmr-selection v0.2.8) to rank features by their relevance to KS stages (measured by ANOVA F-statistics) while minimizing inter-feature redundancy (based on correlation).^82^ The final XGBoost model was retrained using the top 80 ranked features, selected to optimize classification performance while maintaining model interpretability.

### Statistics and reproducibility

All statistical analyses were performed in Python (v3.11.9) unless otherwise noted. Results of DE analyses were tested using the two-sided Wilcoxon rank-sum test as implemented in Scanpy, with P-values adjusted for multiple testing using the Benjamini-Hochberg false discovery rate (FDR). Genes were considered significant if they satisfied both a log2 fold-change cutoff (≥1 for bulk analyses, ≥6 for stage-wise analyses) and an adjusted P-value threshold (-log10 FDR ≥20 or ≥25, depending on analysis).

To assess reproducibility, DE analyses were repeated across independent resampling folds, and significance calls were confirmed to be consistent across runs. For vessel density comparisons across stages, pairwise differences were evaluated using the two-sided Mann-Whitney U test, with Benjamini-Hochberg correction for multiple testing (α = 0.05). Sample information is reported in Table S2.

## Data and code availability

All raw and processed scSRT datasets generated in this study will be deposited to Zenodo and made available upon publication. This includes the full annotated h5ad object containing cell-level expression, spatial coordinates, infection status, niche annotations, and metadata, as well as the per-core summary tables used in pseudotime and infection density analyses.

All custom analysis code, including preprocessing pipelines, spatial neighborhood calculations, space-time plotting functions, regression models, and figure-generation scripts, will be made publicly available on GitHub upon publication. The repository will be fully documented and provides Jupyter notebooks reproducing each figure panel. Software dependencies are specified in an environment.yml file to enable exact replication.

Representative figures and source data tables are provided in Supplementary Data. All results in the manuscript can be regenerated directly from the Zenodo dataset using the GitHub codebase.

## Acknowledgments

We thank members of Drs. Shou-Jiang Gao and Yufei Huang laboratories for technical assistance and discussions. This study was supported by grants from the National Institutes of Health (CA096512, CA284554, CA278812, CA291244 and CA124332 to S.-J. Gao; U01CA279618 and R21GM155774 to Y. Huang), UPMC Hillman Cancer Center Startup Funds to S.-J. Gao and Y. Huang, and in part by award P30CA047904. This research was also supported in part by the University of Pittsburgh Center for Research Computing, RRID:SCR_022735. Specifically, this work used the HTC cluster, which is supported by S10OD028483.

## Author contributions

Conceptualization, S.-J.G.; methodology, W.M., A.D., H.S., R.N. and Y.H.; investigation, W.M., A.D., H.S., R.N., Y.H., S.-J.G.; writing – original draft, W.M., A.D., Y.H., S.-J.G.; writing – review & editing, W.M., A.D., H.S., R.N., P.M.B., M.M., Y.H., S.-J.G; funding acquisition, Y.H., S.-J.G.; supervision, Y.H., S.-J.G.

## Supplementary information

The online version contains supplementary material available at:

**Correspondence and requests for materials** should be addressed to Shou-Jiang Gao or Yufei Huang.

## Supplemental Information

### Supplemental Tables

**Table S1.** A panel of KSHV and cellular genes examined in Kaposi’s sarcoma tumors and normal control tissues by single-cell spatially resolved transcriptomics.

| Gene | Ensemble ID |
| --- | --- |
| ACER1 | ENSG00000167769 |
| ACTA2 | ENSG00000107796 |
| ADAM12 | ENSG00000148848 |
| AHNAK2 | ENSG00000185567 |
| AIF1 | ENSG00000204472 |
| AKR1C1 | ENSG00000187134 |
| ALOX5AP | ENSG00000132965 |
| APCDD1 | ENSG00000154856 |
| APOD | ENSG00000189058 |
| AQP1 | ENSG00000240583 |
| AQP3 | ENSG00000165272 |
| AREG | ENSG00000109321 |
| ARHGDIB | ENSG00000111348 |
| ASPN | ENSG00000106819 |
| ATF3 | ENSG00000162772 |
| AXL | ENSG00000167601 |
| BASP1 | ENSG00000176788 |
| BCAN | ENSG00000132692 |
| BHLHE41 | ENSG00000123095 |
| BIRC3 | ENSG00000023445 |
| BIRC5 | ENSG00000089685 |
| C11orf96 | ENSG00000187479 |
| C15orf48 | ENSG00000166920 |
| C1QA | ENSG00000173372 |
| C1orf54 | ENSG00000118292 |
| C5AR1 | ENSG00000197405 |
| CALCRL | ENSG00000064989 |
| CALD1 | ENSG00000122786 |
| CAPNS2 | ENSG00000256812 |
| CBFA2T3 | ENSG00000129993 |
| CCL19 | ENSG00000172724 |
| CCL22 | ENSG00000102962 |
| CCND1 | ENSG00000110092 |
| CCR7 | ENSG00000126353 |
| CD1A | ENSG00000158477 |
| CD1B | ENSG00000158485 |
| CD3D | ENSG00000167286 |
| CD3E | ENSG00000198851 |
| CD3G | ENSG00000160654 |
| CD40LG | ENSG00000102245 |
| CD52 | ENSG00000169442 |
| CD68 | ENSG00000129226 |
| CD69 | ENSG00000110848 |
| CD79A | ENSG00000105369 |
| CD83 | ENSG00000112149 |
| CD8A | ENSG00000153563 |
| CD93 | ENSG00000125810 |
| CDC20 | ENSG00000117399 |
| CDH5 | ENSG00000179776 |
| CDK1 | ENSG00000170312 |
| CENPF | ENSG00000117724 |
| CES1 | ENSG00000198848 |
| CLDN1 | ENSG00000163347 |
| CLDN4 | ENSG00000189143 |
| CLDN5 | ENSG00000184113 |
| CLEC10A | ENSG00000132514 |
| CLEC9A | ENSG00000197992 |
| COCH | ENSG00000100473 |
| COL17A1 | ENSG00000065618 |
| COL5A2 | ENSG00000204262 |
| COL6A1 | ENSG00000142156 |
| COL6A2 | ENSG00000142173 |
| COL6A3 | ENSG00000163359 |
| COTL1 | ENSG00000103187 |
| CPE | ENSG00000109472 |
| CPVL | ENSG00000106066 |
| CRABP2 | ENSG00000143320 |
| CRYAB | ENSG00000109846 |
| CTSW | ENSG00000172543 |
| CTSZ | ENSG00000101160 |
| CXADR | ENSG00000154639 |
| CXCL12 | ENSG00000107562 |
| CXCR4 | ENSG00000121966 |
| CYB561A3 | ENSG00000162144 |
| DCT | ENSG00000080166 |
| DEFB1 | ENSG00000164825 |
| DSC1 | ENSG00000134765 |
| DST | ENSG00000151914 |
| DUSP2 | ENSG00000158050 |
| ECSCR | ENSG00000249751 |
| EDNRB | ENSG00000136160 |
| ENAH | ENSG00000154380 |
| ENPP1 | ENSG00000197594 |
| FABP4 | ENSG00000170323 |
| FAM210B | ENSG00000124098 |
| FBLN1 | ENSG00000077942 |
| FBXO7 | ENSG00000100225 |
| FCER1A | ENSG00000179639 |
| FCER1G | ENSG00000158869 |
| FGFBP1 | ENSG00000137440 |
| FKBP8 | ENSG00000105701 |
| FRZB | ENSG00000162998 |
| FSCN1 | ENSG00000075618 |
| GATA2 | ENSG00000179348 |
| GEM | ENSG00000164949 |
| GIMAP7 | ENSG00000179144 |
| GNG11 | ENSG00000127920 |
| GPR157 | ENSG00000180758 |
| GPR183 | ENSG00000169508 |
| GPR4 | ENSG00000177464 |
| GYPC | ENSG00000136732 |
| GZMK | ENSG00000113088 |
| HAPLN1 | ENSG00000145681 |
| HBD | ENSG00000223609 |
| HES1 | ENSG00000114315 |
| HIST1H4C | ENSG00000197061 |
| HMGB2 | ENSG00000164104 |
| HMGN2 | ENSG00000198830 |
| HTRA1 | ENSG00000166033 |
| ID2 | ENSG00000115738 |
| ID4 | ENSG00000172201 |
| IDO1 | ENSG00000131203 |
| IFI27 | ENSG00000165949 |
| IFITM1 | ENSG00000185885 |
| IFNG | ENSG00000111537 |
| IGFBP2 | ENSG00000115457 |
| IGFBP3 | ENSG00000146674 |
| IGFBP5 | ENSG00000115461 |
| IGFBP7 | ENSG00000163453 |
| IL1B | ENSG00000125538 |
| IL2RA | ENSG00000134460 |
| IL32 | ENSG00000008517 |
| IL33 | ENSG00000137033 |
| IL3RA | ENSG00000185291 |
| IL4I1 | ENSG00000104951 |
| INHBA | ENSG00000122641 |
| INSIG1 | ENSG00000186480 |
| IRF4 | ENSG00000137265 |
| IRF8 | ENSG00000140968 |
| ITGA6 | ENSG00000091409 |
| ITGB1 | ENSG00000150093 |
| ITM2A | ENSG00000078596 |
| KIT | ENSG00000157404 |
| KLRB1 | ENSG00000111796 |
| KRT15 | ENSG00000171346 |
| KRT18 | ENSG00000111057 |
| KRT2 | ENSG00000172867 |
| KRT5 | ENSG00000186081 |
| KRTDAP | ENSG00000188508 |
| LAMB3 | ENSG00000196878 |
| LCK | ENSG00000182866 |
| LENG8 | ENSG00000167615 |
| LEPR | ENSG00000116678 |
| LGALS2 | ENSG00000100079 |
| LINC00636 | ENSG00000240423 |
| LSAMP | ENSG00000185565 |
| LST1 | ENSG00000204482 |
| LTB | ENSG00000227507 |
| LUM | ENSG00000139329 |
| LY6D | ENSG00000167656 |
| LYPD3 | ENSG00000124466 |
| LYVE1 | ENSG00000133800 |
| LYZ | ENSG00000090382 |
| MAL2 | ENSG00000147676 |
| MARCKSL1 | ENSG00000175130 |
| MEF2C | ENSG00000081189 |
| MFAP4 | ENSG00000166482 |
| MFAP5 | ENSG00000197614 |
| MGP | ENSG00000111341 |
| MIR205HG | ENSG00000230937 |
| MITF | ENSG00000187098 |
| MKI67 | ENSG00000148773 |
| MLANA | ENSG00000120215 |
| MMP2 | ENSG00000087245 |
| MMP27 | ENSG00000137675 |
| MMRN1 | ENSG00000138722 |
| MNDA | ENSG00000163563 |
| MT1F | ENSG00000198417 |
| MT1X | ENSG00000187193 |
| MYC | ENSG00000136997 |
| MYCT1 | ENSG00000120279 |
| MYH11 | ENSG00000133392 |
| MYL9 | ENSG00000101335 |
| MZB1 | ENSG00000170476 |
| NDUFA4L2 | ENSG00000185633 |
| NLGN4Y | ENSG00000165246 |
| NMB | ENSG00000197696 |
| NOTCH3 | ENSG00000074181 |
| NR2F2 | ENSG00000185551 |
| NR4A2 | ENSG00000153234 |
| NSG1 | ENSG00000168824 |
| NUSAP1 | ENSG00000137804 |
| PBXIP1 | ENSG00000163346 |
| PCDH7 | ENSG00000169851 |
| PDGFRA | ENSG00000134853 |
| PKHD1L1 | ENSG00000205038 |
| PKIB | ENSG00000135549 |
| PLAC9 | ENSG00000189129 |
| PLP1 | ENSG00000123560 |
| PLVAP | ENSG00000130300 |
| PMEL | ENSG00000185664 |
| POSTN | ENSG00000133110 |
| PROX1 | ENSG00000117707 |
| PTGDS | ENSG00000107317 |
| PTGER3 | ENSG00000050628 |
| PTGR1 | ENSG00000106853 |
| PTN | ENSG00000105894 |
| PTPRCAP | ENSG00000213402 |
| PTTG1 | ENSG00000164611 |
| QPCT | ENSG00000115828 |
| RAMP2 | ENSG00000131477 |
| RASSF8 | ENSG00000123094 |
| RGCC | ENSG00000102760 |
| RGS1 | ENSG00000090104 |
| RGS5 | ENSG00000143248 |
| RHOV | ENSG00000104140 |
| RNASE1 | ENSG00000129538 |
| RRM2 | ENSG00000171848 |
| S100A13 | ENSG00000189171 |
| S100A14 | ENSG00000189334 |
| S100B | ENSG00000160307 |
| SELE | ENSG00000007908 |
| SERPINB2 | ENSG00000197632 |
| SERPINB5 | ENSG00000206075 |
| SFRP1 | ENSG00000104332 |
| SFRP2 | ENSG00000145423 |
| SLC25A37 | ENSG00000147454 |
| SLC25A39 | ENSG00000013306 |
| SLC2A3 | ENSG00000059804 |
| SLC8A1 | ENSG00000183023 |
| SLPI | ENSG00000124107 |
| SNCA | ENSG00000145335 |
| SOD3 | ENSG00000109610 |
| SOSTDC1 | ENSG00000171243 |
| SOX15 | ENSG00000129194 |
| SOX17 | ENSG00000164736 |
| SPARCL1 | ENSG00000152583 |
| SPRY1 | ENSG00000164056 |
| STMN1 | ENSG00000117632 |
| STRADB | ENSG00000082146 |
| SYNPO2 | ENSG00000172403 |
| TACSTD2 | ENSG00000184292 |
| TAGLN | ENSG00000149591 |
| TFAP2A | ENSG00000137203 |
| TFAP2B | ENSG00000008196 |
| TFF3 | ENSG00000160180 |
| TFPI | ENSG00000003436 |
| TFPI2 | ENSG00000105825 |
| THBS2 | ENSG00000186340 |
| THY1 | ENSG00000154096 |
| TM4SF1 | ENSG00000169908 |
| TMEM150C | ENSG00000249242 |
| TMEM176A | ENSG00000002933 |
| TMEM45A | ENSG00000181458 |
| TNFRSF17 | ENSG00000048462 |
| TOP2A | ENSG00000131747 |
| TP63 | ENSG00000073282 |
| TRPM1 | ENSG00000134160 |
| TSC22D1 | ENSG00000102804 |
| TSPAN33 | ENSG00000158457 |
| TUBB2B | ENSG00000137285 |
| TYMS | ENSG00000176890 |
| TYR | ENSG00000077498 |
| TYROBP | ENSG00000011600 |
| TYRP1 | ENSG00000107165 |
| UBE2C | ENSG00000175063 |
| VIM | ENSG00000026025 |
| VWF | ENSG00000110799 |
| WDFY4 | ENSG00000128815 |
| ZNF667-AS1 | ENSG00000166770 |
| CTNNB1 | ENSG00000168036 |
| ICAM1 | ENSG00000090339 |
| VCAM1 | ENSG00000162692 |
| CD34 | ENSG00000174059 |
| NRP1 | ENSG00000099250 |
| CD209 | ENSG00000090659 |
| HSPG2 | ENSG00000142798 |
| EFNA2 | ENSG00000099617 |
| SLC7A11 | ENSG00000151012 |
| ITGA3 | ENSG00000005884 |
| ITGB3 | ENSG00000259207 |
| ITGAV | ENSG00000138448 |
| ITGB5 | ENSG00000082781 |
| CXCL10 | ENSG00000169245 |
| IL1R2 | ENSG00000115590 |
| CCL3 | ENSG00000277632 |
| SOCS3 | ENSG00000184557 |
| CCR8 | ENSG00000179934 |
| PTGS2 | ENSG00000073756 |
| ENG | ENSG00000106991 |
| VEGFA | ENSG00000112715 |
| KDR | ENSG00000128052 |
| VEGFC | ENSG00000150630 |
| IL1A | ENSG00000115008 |
| IL6 | ENSG00000136244 |
| TNF | ENSG00000232810 |
| FGF2 | ENSG00000138685 |
| PDGFRB | ENSG00000113721 |
| PDGFA | ENSG00000197461 |
| TGFB1 | ENSG00000105329 |
| CXCL8 | ENSG00000169429 |
| MMP9 | ENSG00000100985 |
| IL6R | ENSG00000160712 |
| IL6ST | ENSG00000134352 |
| IL18 | ENSG00000150782 |
| IL10 | ENSG00000136634 |
| IL12B | ENSG00000113302 |
| KSHV.K2 | HQ404500 |
| KSHV.ORF50 | HQ404500 |
| KSHV.ORF57 | HQ404500 |
| KSHV.ORF59 | HQ404500 |
| KSHV.ORF65 | HQ404500 |
| KSHV.ORF71 | HQ404500 |
| KSHV.ORF72 | HQ404500 |
| KSHV.ORF73 | HQ404500 |
| KSHV.K9 | HQ404500 |

**Table S2.**
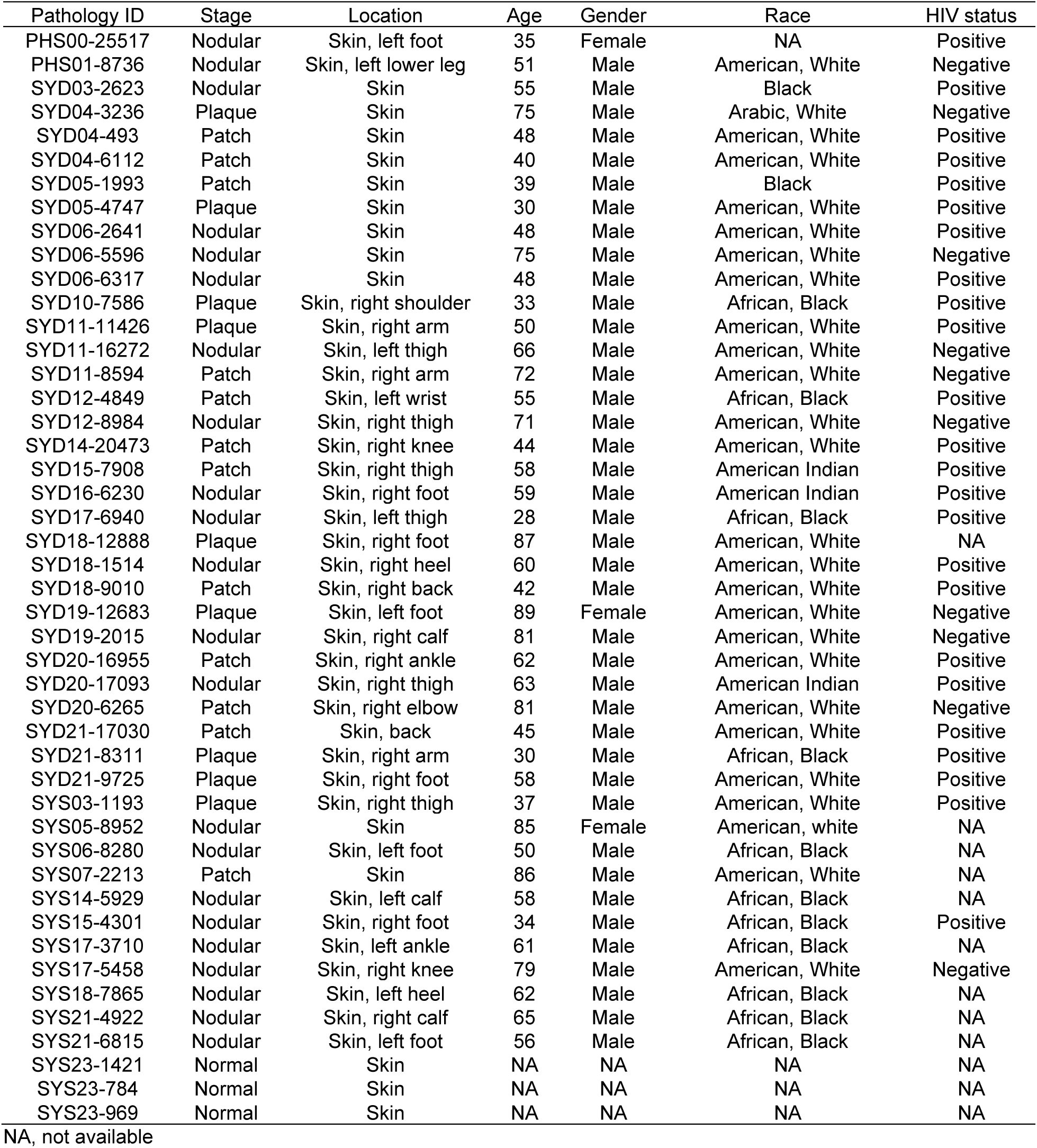
Kaposi’s sarcoma patients, control subjects, and their information examined by single-cell spatially resolved transcriptomics.

**Table S3.**
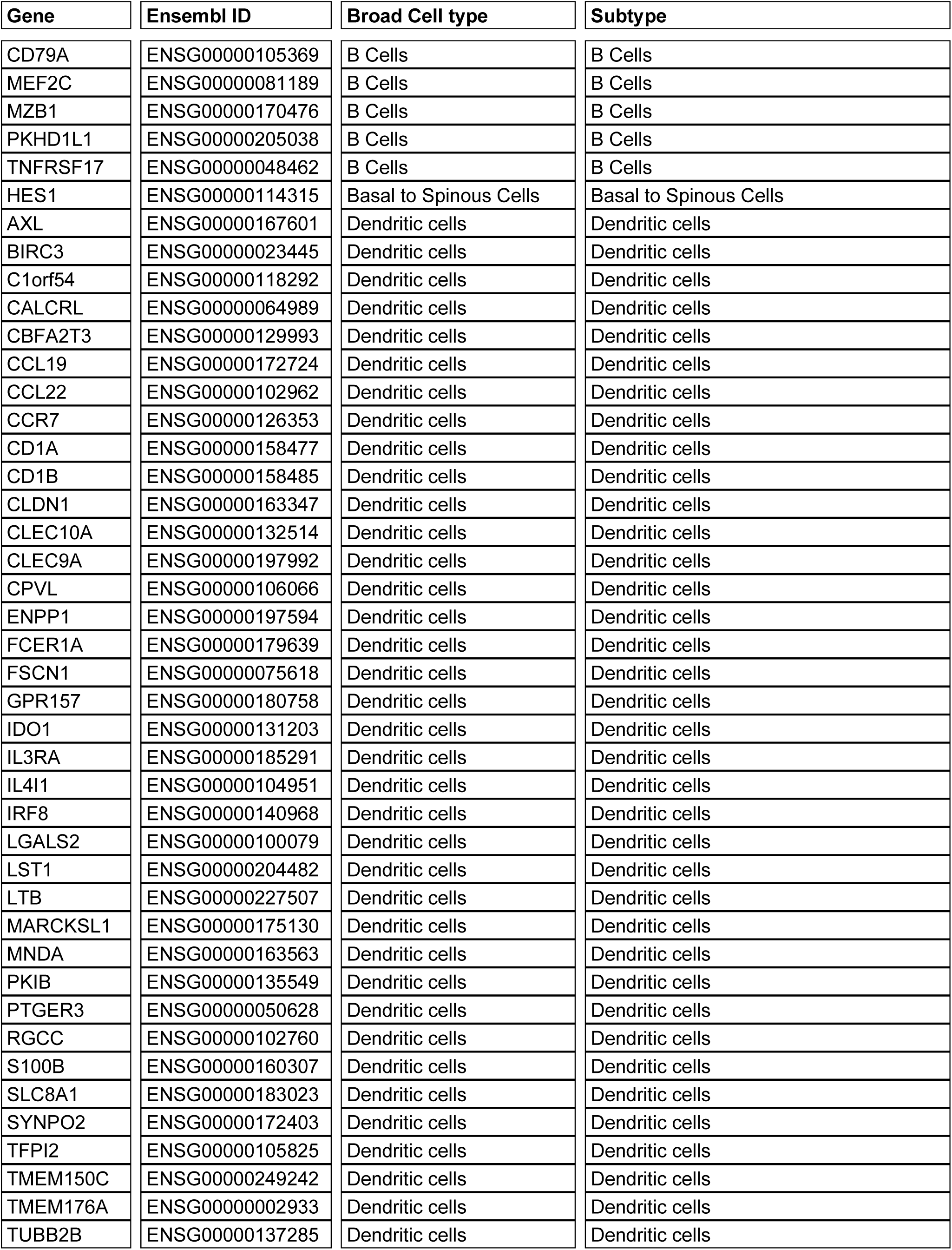
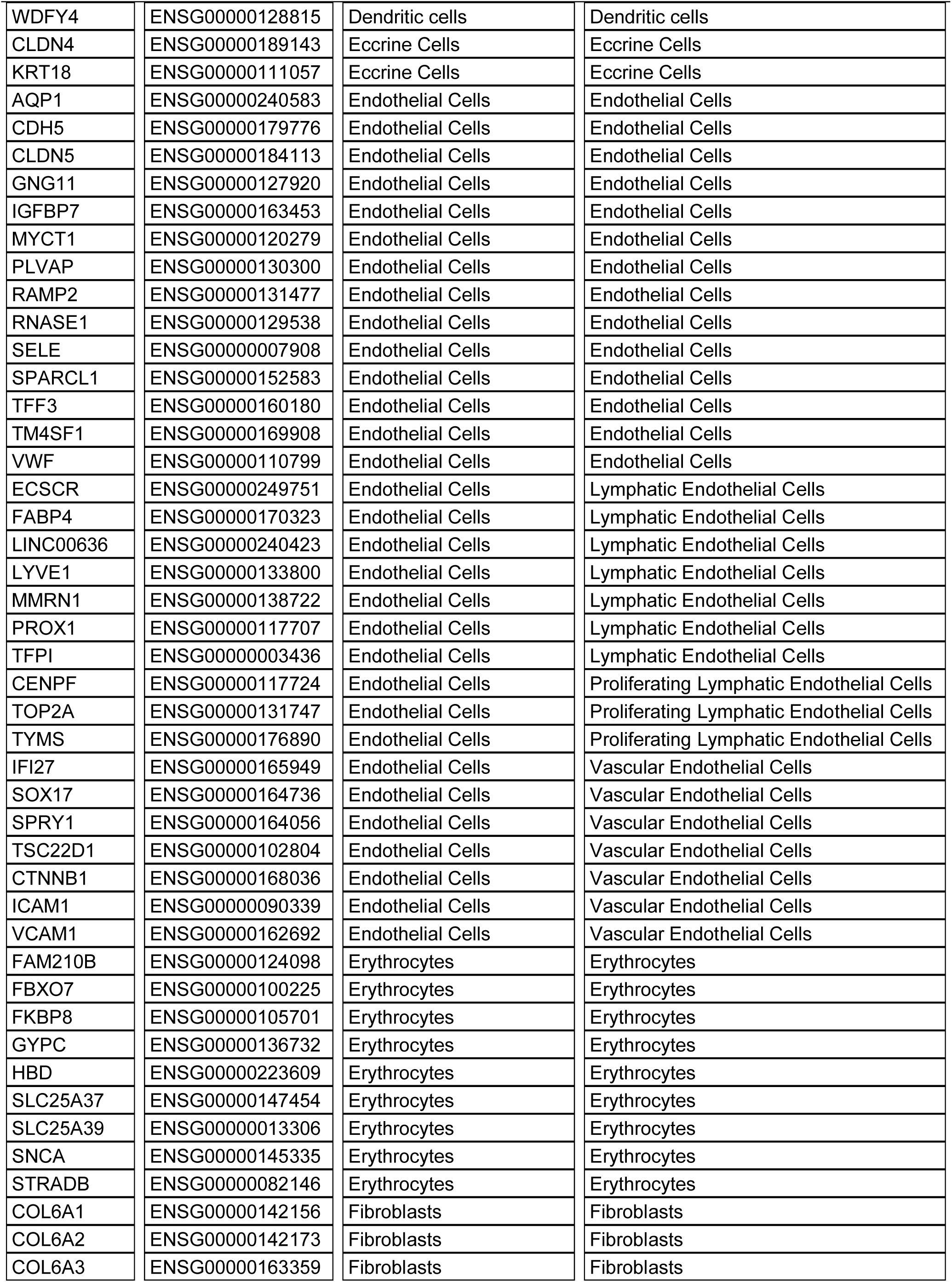
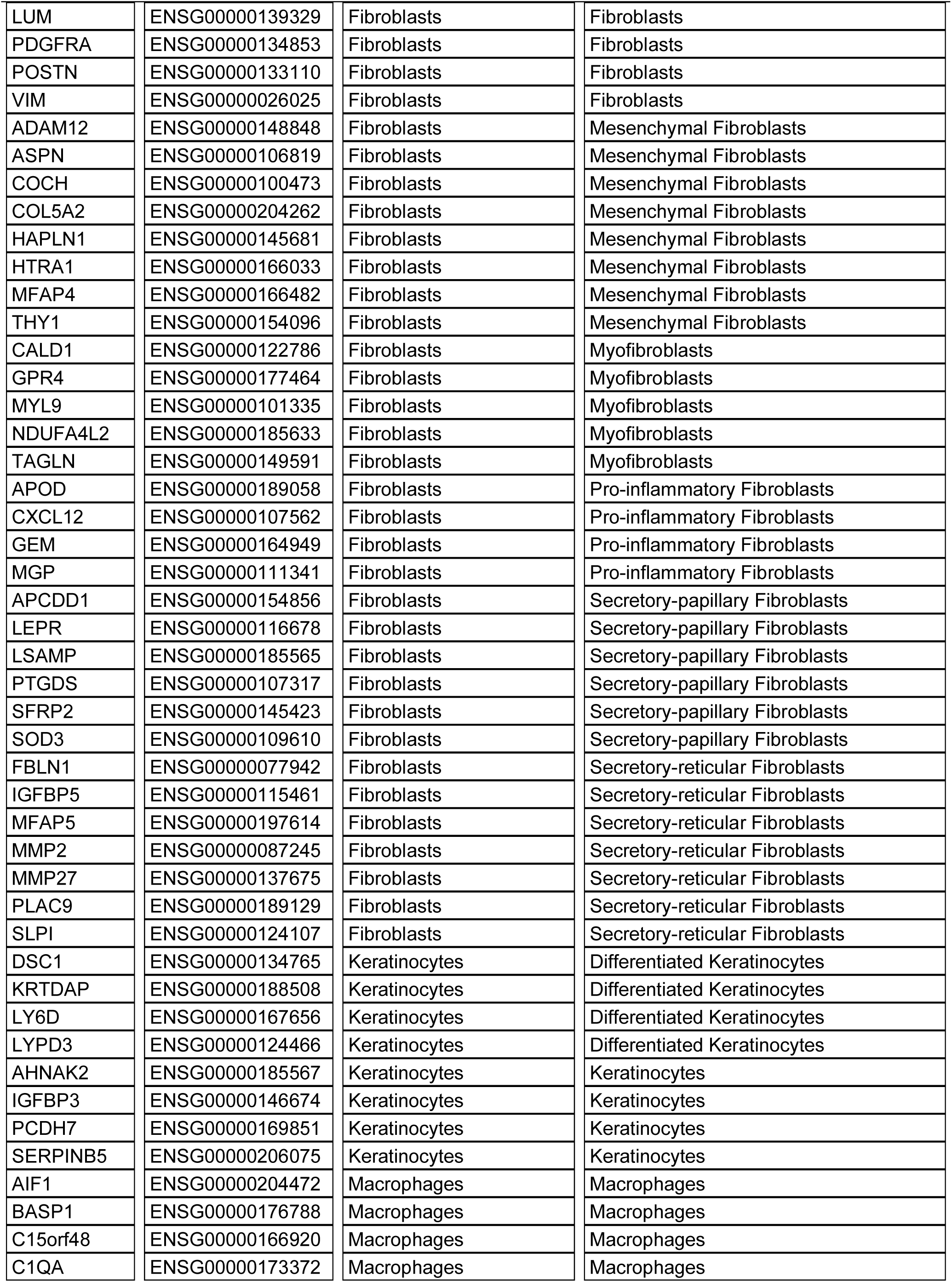
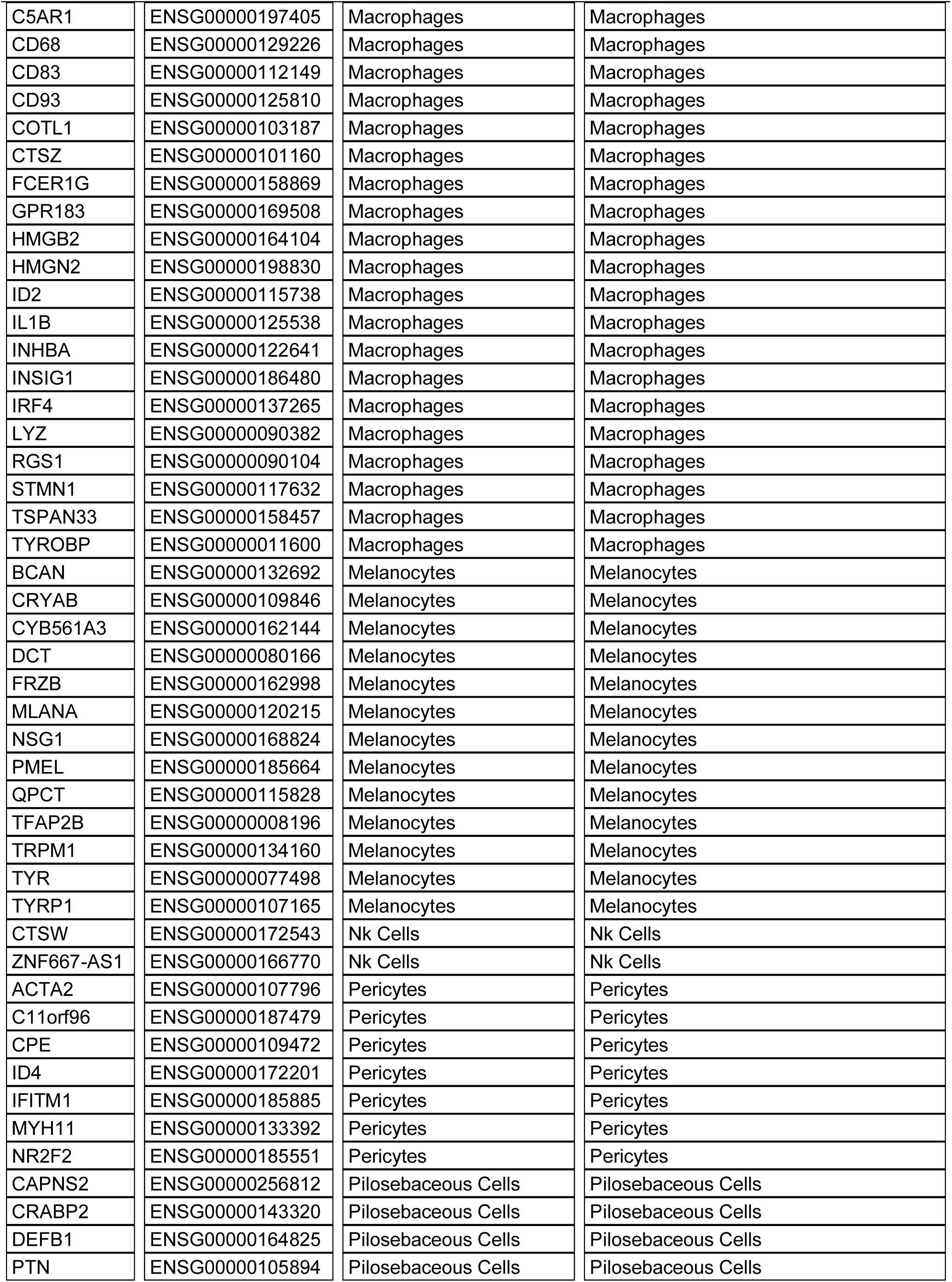
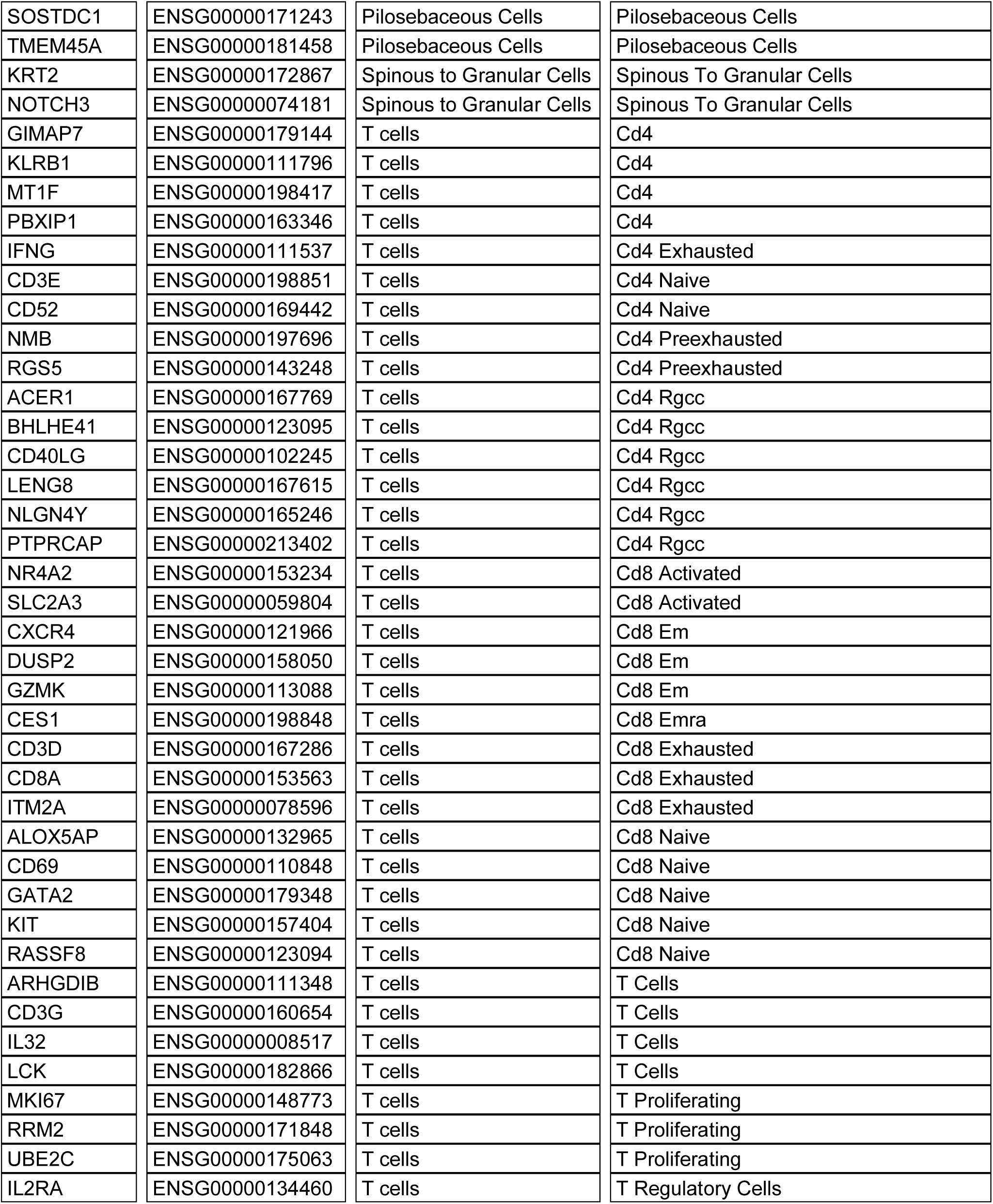
Cellular genes used for cell typing and subtyping.

**Table S4.**
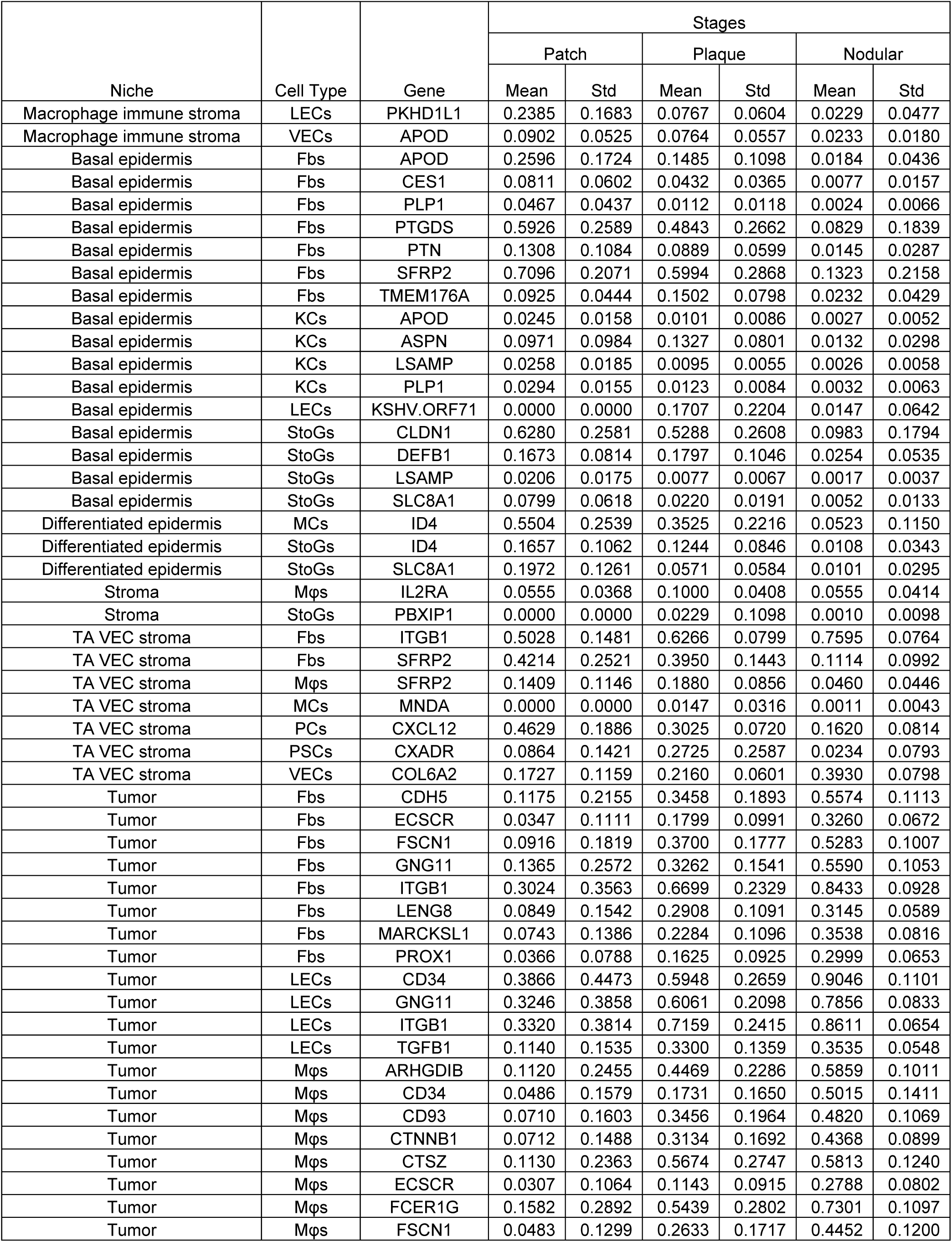
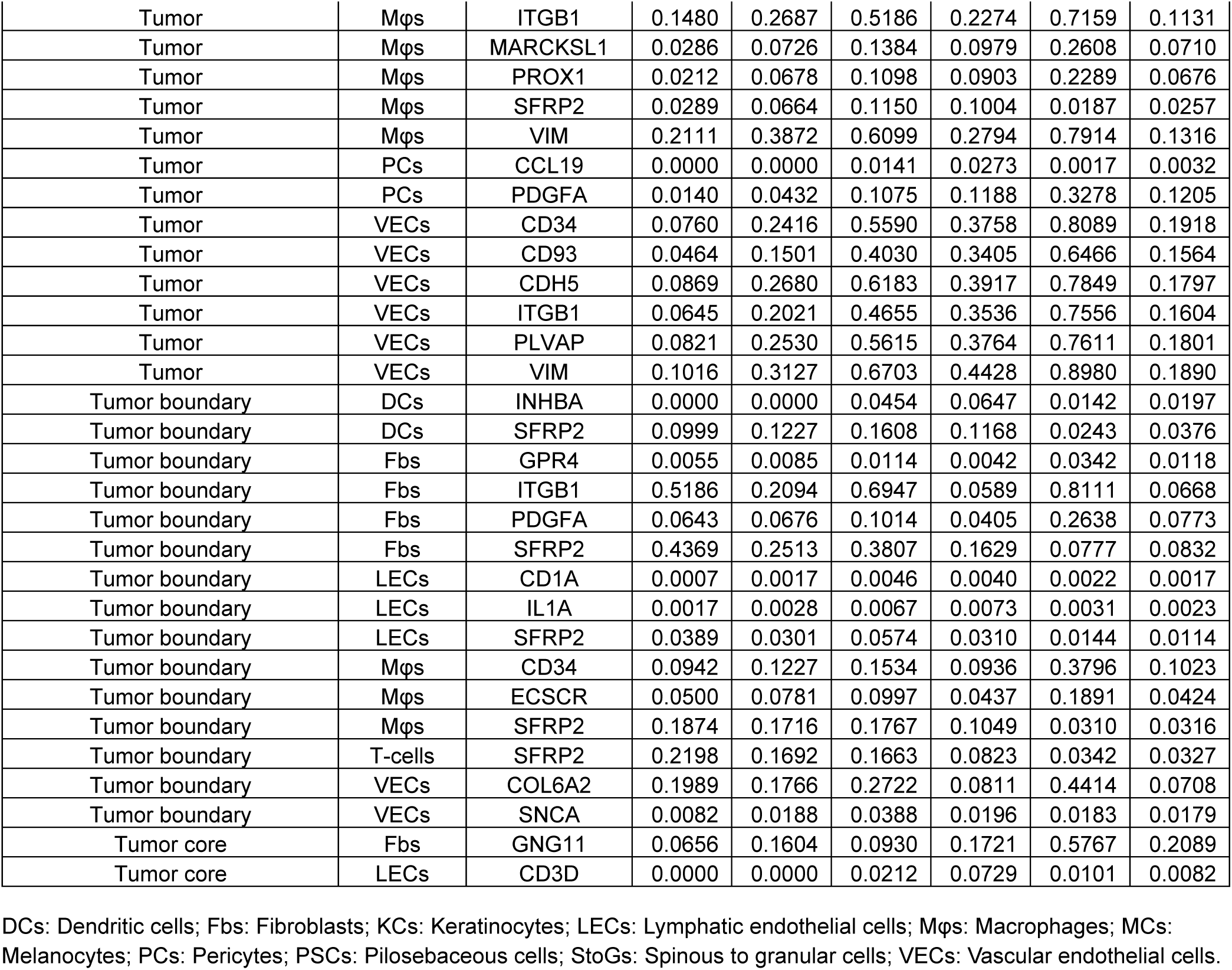
Top spatial single-cell features for predicting Kaposi’s sarcoma progression.

### Supplementary Figures

**Figure S1.**
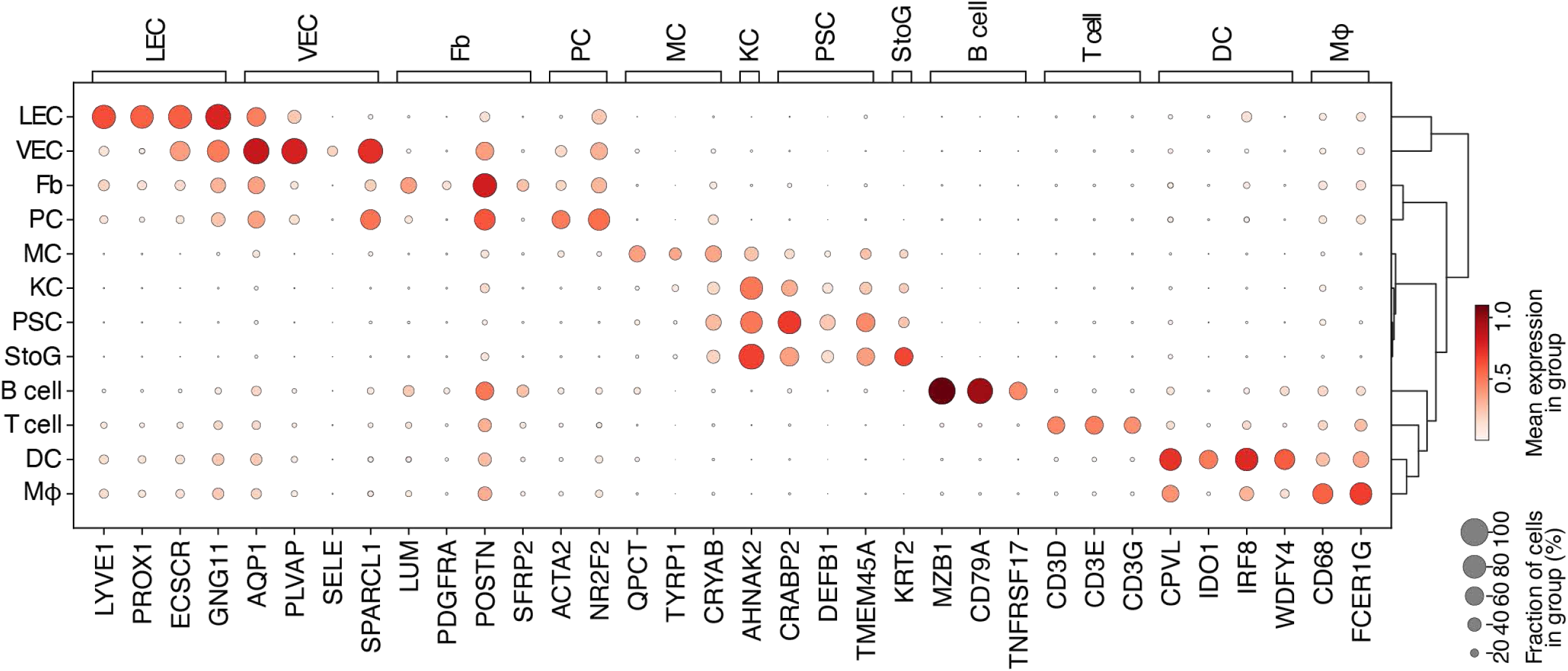
Bubble heat map showing annotation of major cell types in Kaposi’s sarcoma tissues using a marker-based algorithm.

**Figure S2.**
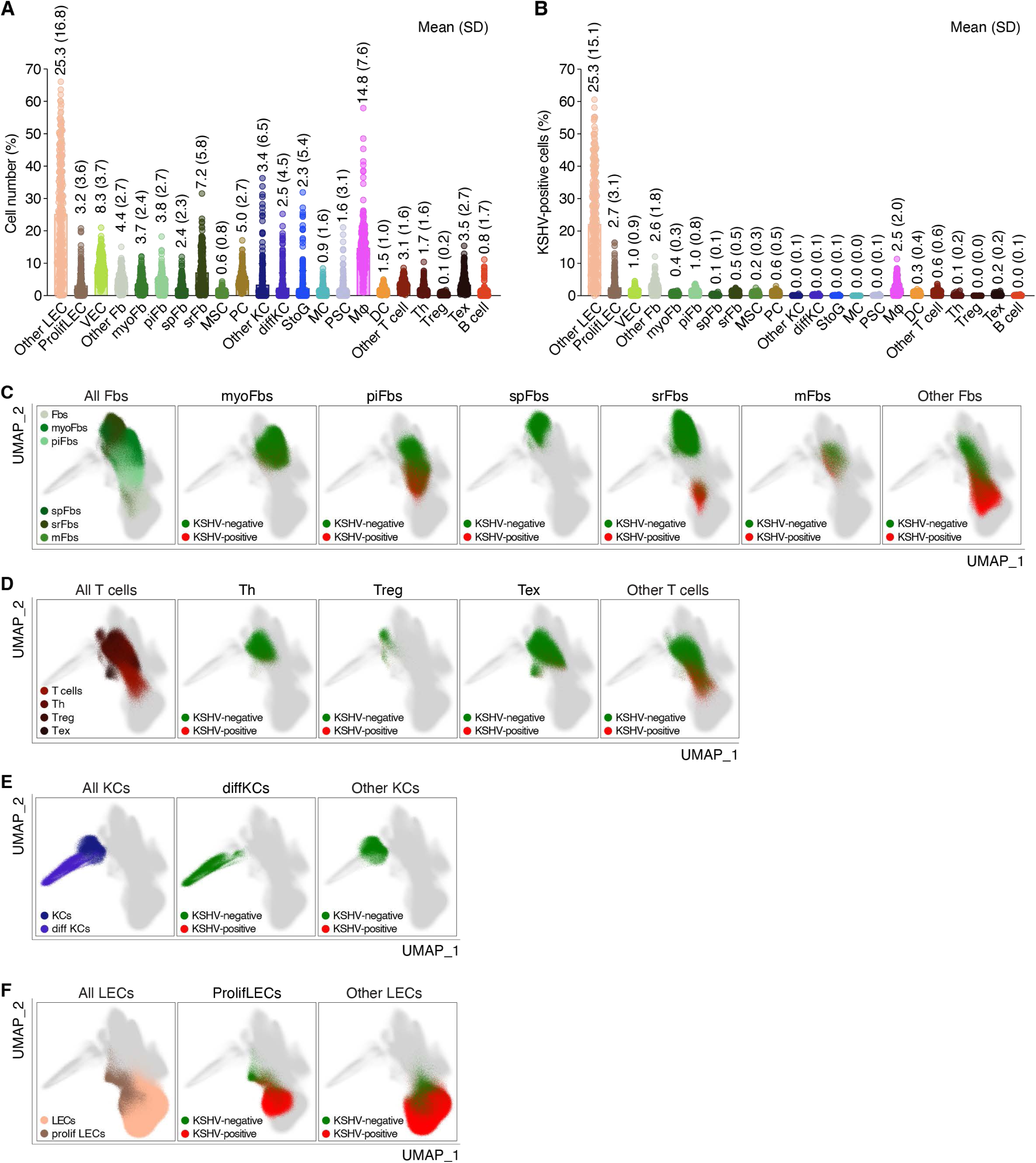
Identification of major cell types and their subtypes using a marker-based algorithm in Kaposi’s sarcoma (KS) (A) Scatter bar plot showing the percentages of identified cell types and their subtypes across all KS samples. Bars represent mean percentages and standard deviations (SD). (B) Scatter bar plot showing KSHV-infected percentages in identified cell types and their subtypes across all KS samples. Bars indicate the mean with SD. (C-F) The UMAP projection of the gene expression of cell subtypes with KSHV infection (red) and without KSHV infection (green) in fibroblasts (C), T-cells (D), Keratinocytes (KCs, E), and lymphatic endothelial cells (LECs, F).

**Figure S3.**
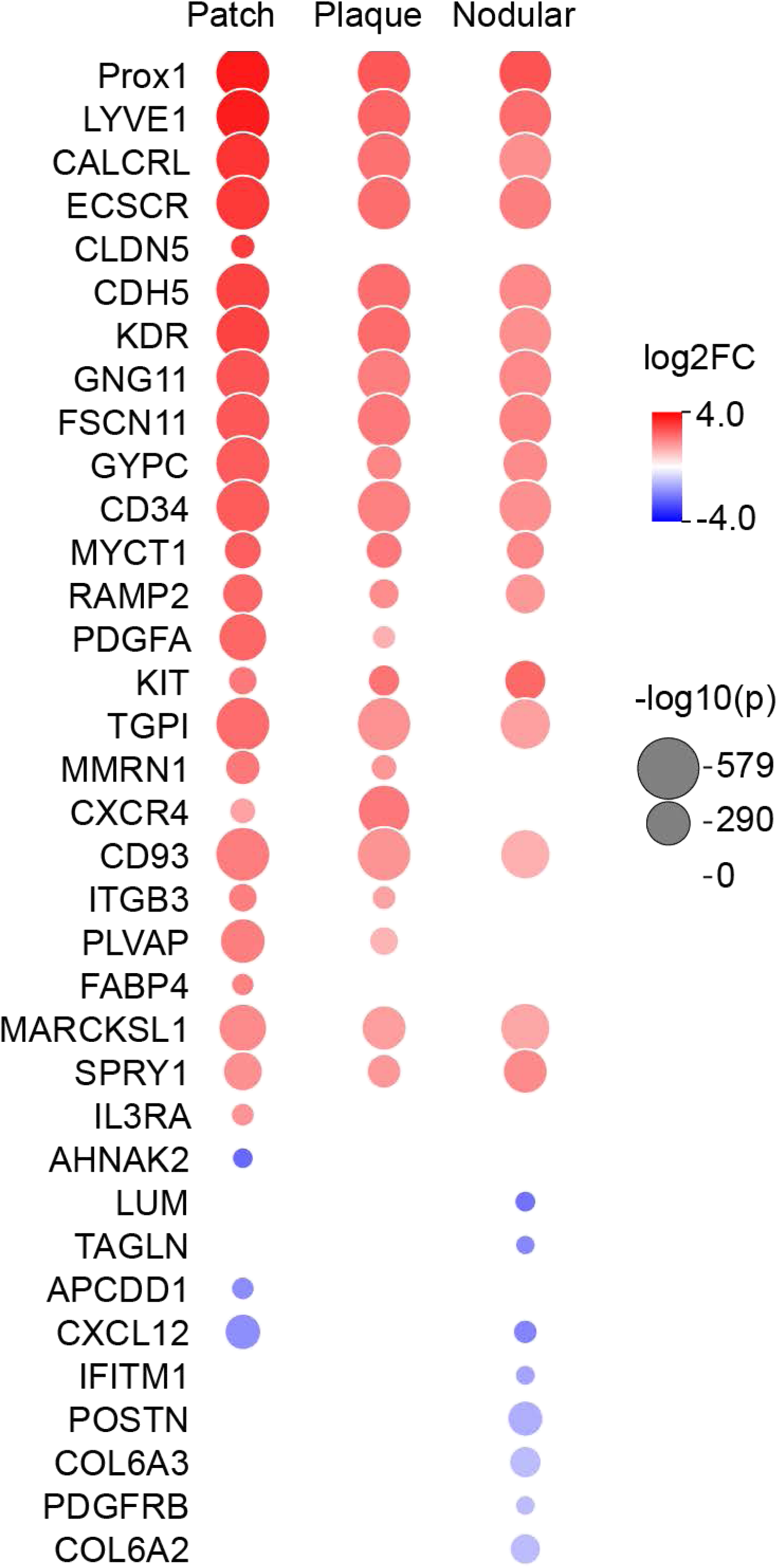
Bubble graph showing differential gene expression along Kaposi’s sarcoma progression.

**Figure S4.**
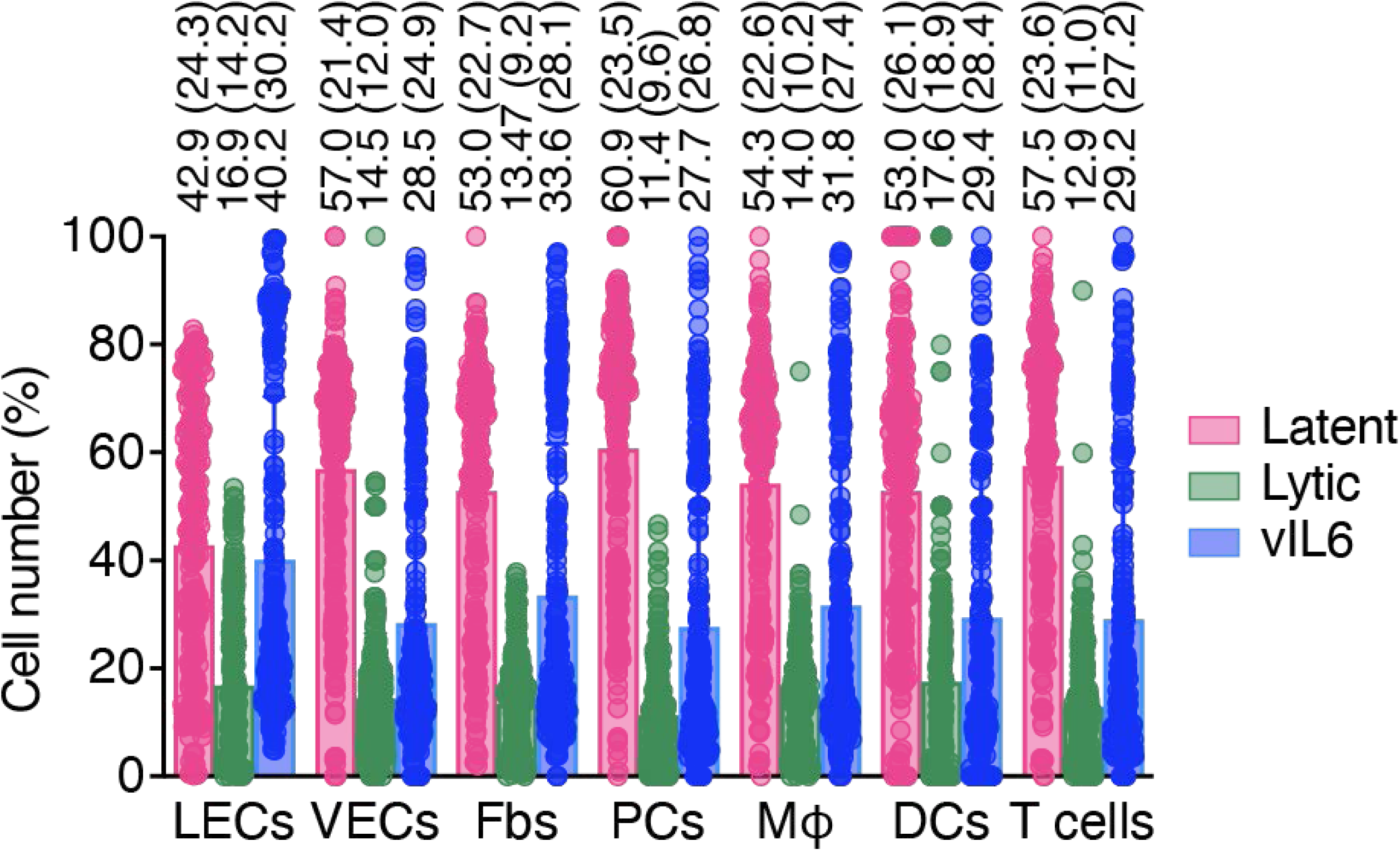
Scatter bar plot showing the percentages of KSHV latent, lytic and vIL6-expressing cells in different identified cell types in Kaposi’s sarcoma. Bars represent mean percentages and standard deviations.

**Figure S5.**
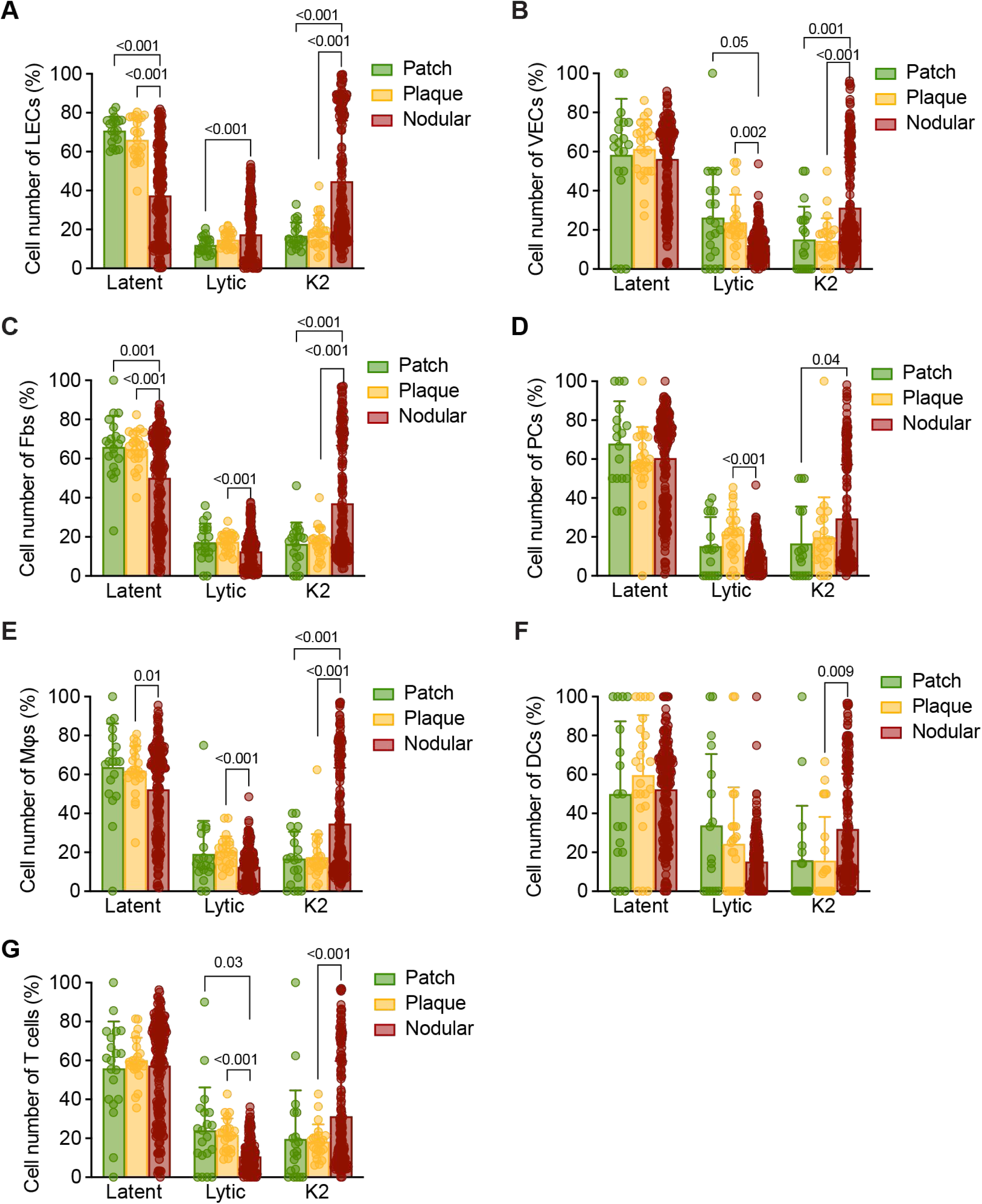
Scatter bar plots showing the percentages of KSHV latent, lytic and vIL6-expressing cells in different identified cell types by Kaposi’s sarcoma stage. (A-G) Cell types include lymphatic endothelial cells (LECs, A), vascular endothelial cells (VECs, B), fibroblasts (Fbs, C), Pericytes (PCs, D), macrophages (Mφs, E), dendritic cells (DCs, F), and T cells (G). Bars represent mean percentages and standard deviations. Statistical significance was determined using RM two-way ANOVA model and Turkey’s multiple comparisons test. P < 0.05 indicates significance.

**Figure S6.**
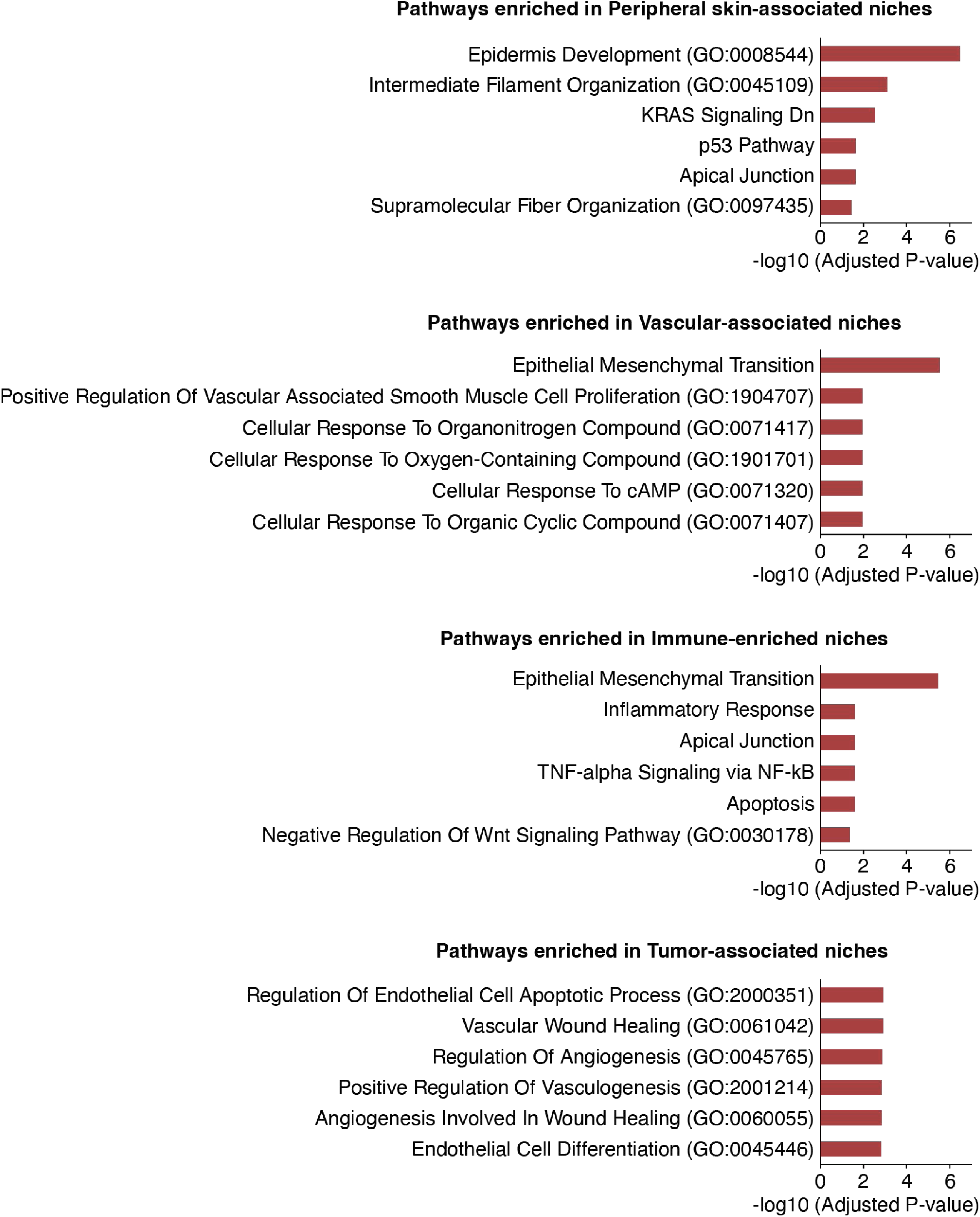
Enrichment pathways revealed in major niches in Kaposi’s sarcoma. (A-D) Pathway enrichment analysis showing enrichment of pathways in Skin-associated niches (A), Vascular-associated niches (B), Immune-enriched niches (C), and Tumor-associated niches (D).

**Figure S7.**
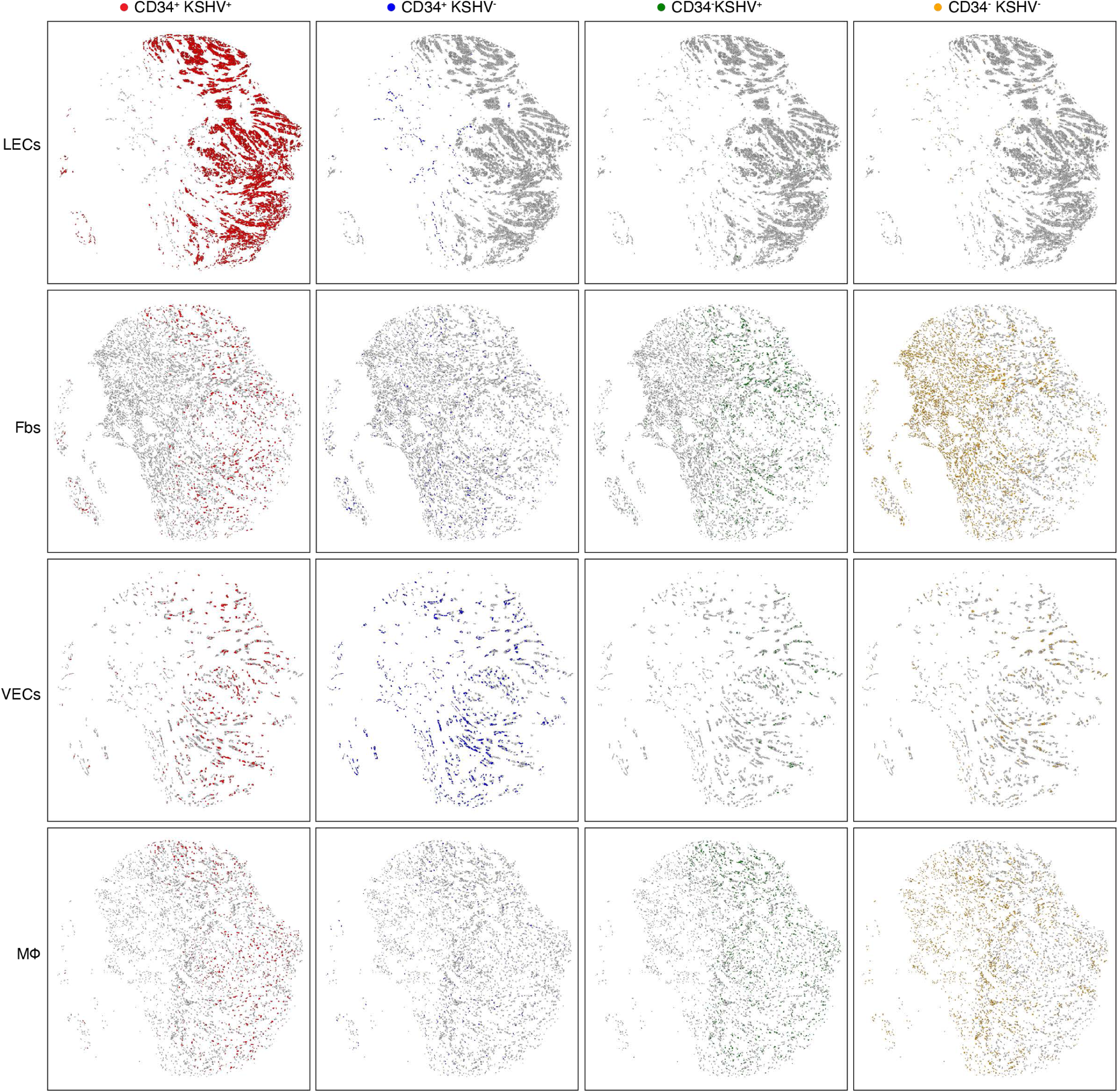
Spatial maps of major cell types Kaposi’s sarcoma by KSHV infection status and CD34 status. (A-D) Cell types include lymphatic endothelial cells (LECs, A), fibroblasts (Fbs, B), vascular endothelial cells (VECs, C), and macrophages (Mφs, D).

**Figure S8.**
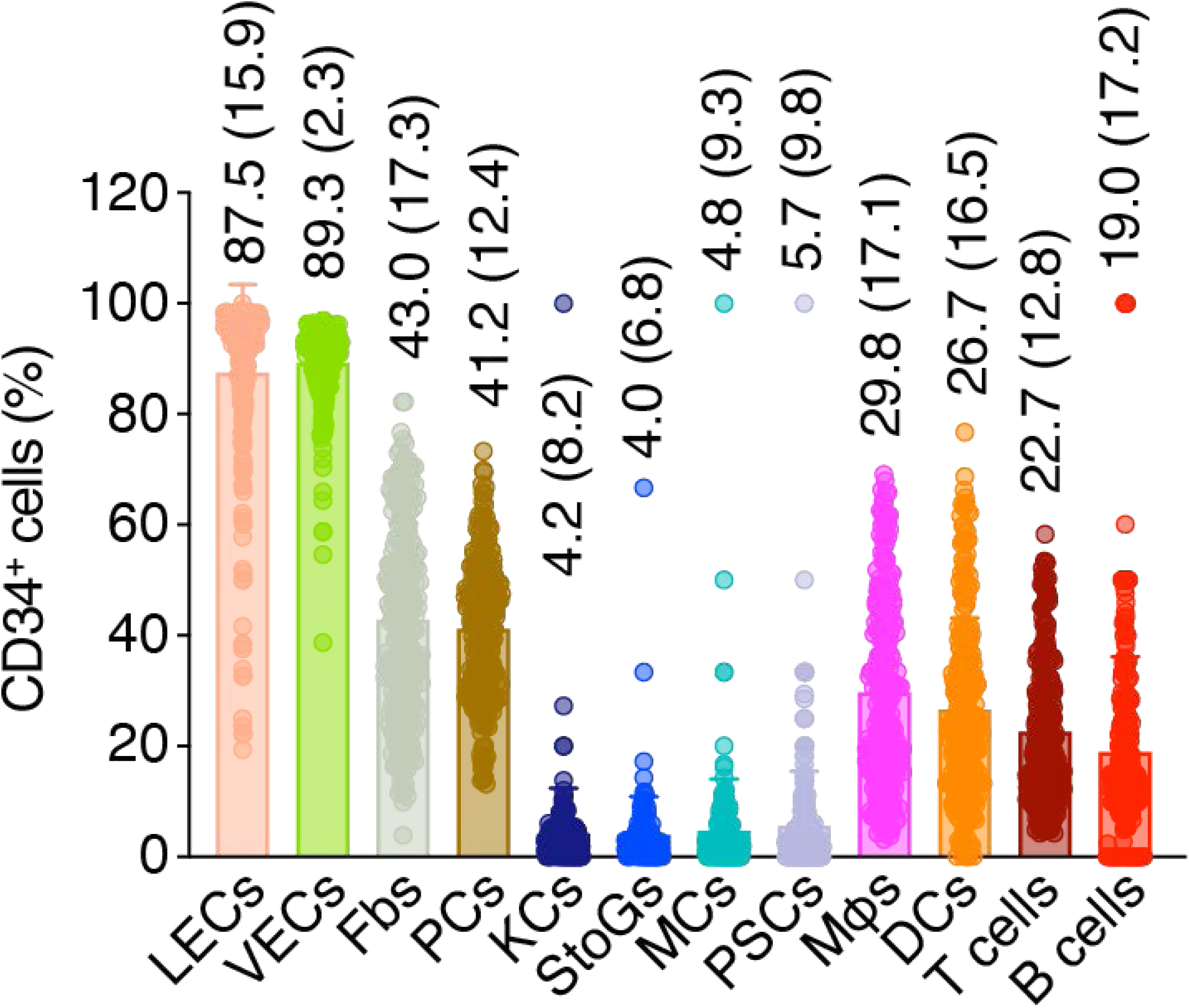
Scatter bar plot showing the percentages of cells expressing CD34 marker in different cell types in Kaposi’s sarcoma. Bars represent mean percentages and standard deviations.

**Figure S9.**
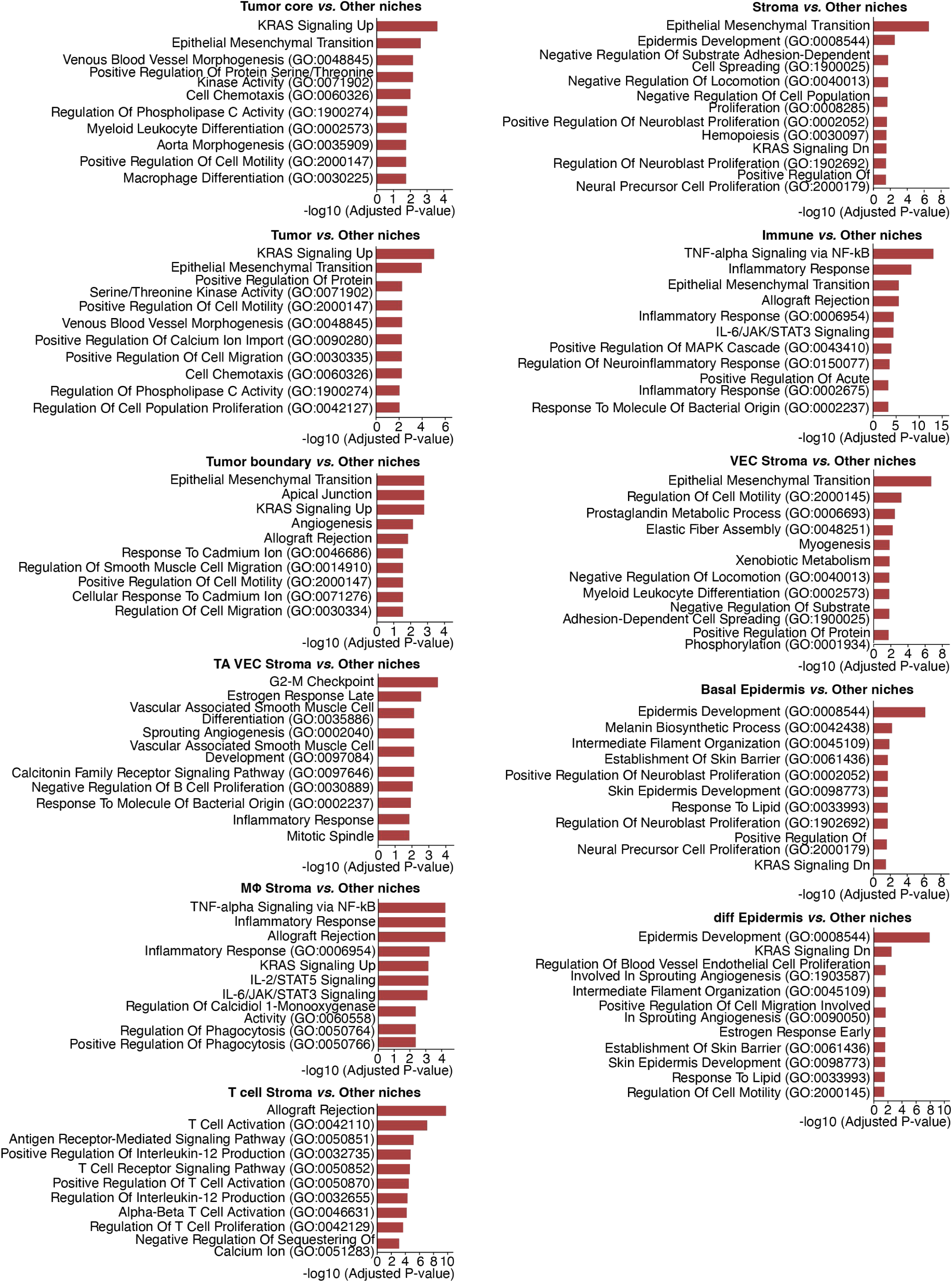
Enrichment pathways revealed in the identified niches in Kaposi’s sarcoma.

**Figure S10.**
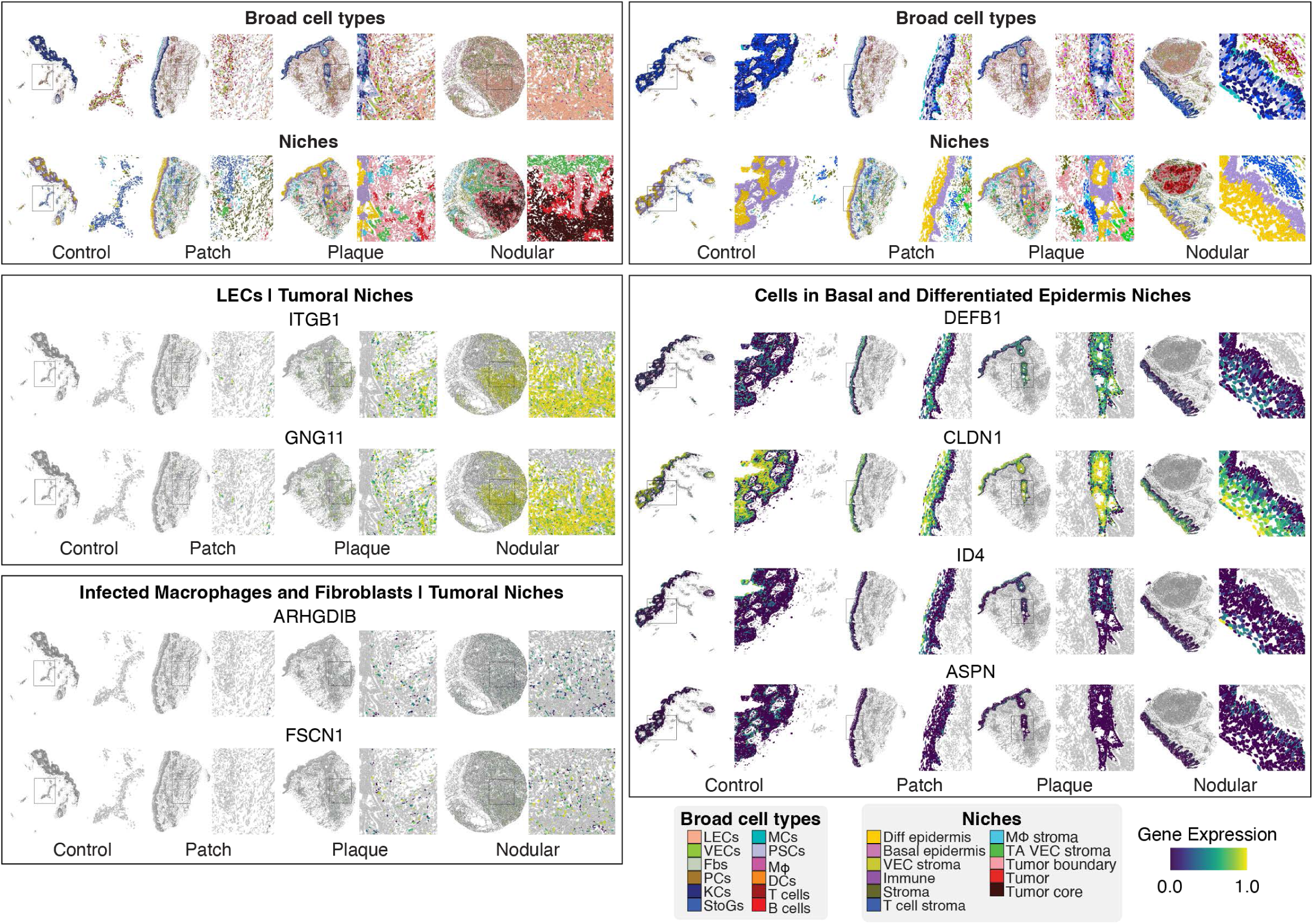
Representative spatial maps of selected spatial single-cell features by Kaposi’s sarcoma stage.

**Figure S11.**
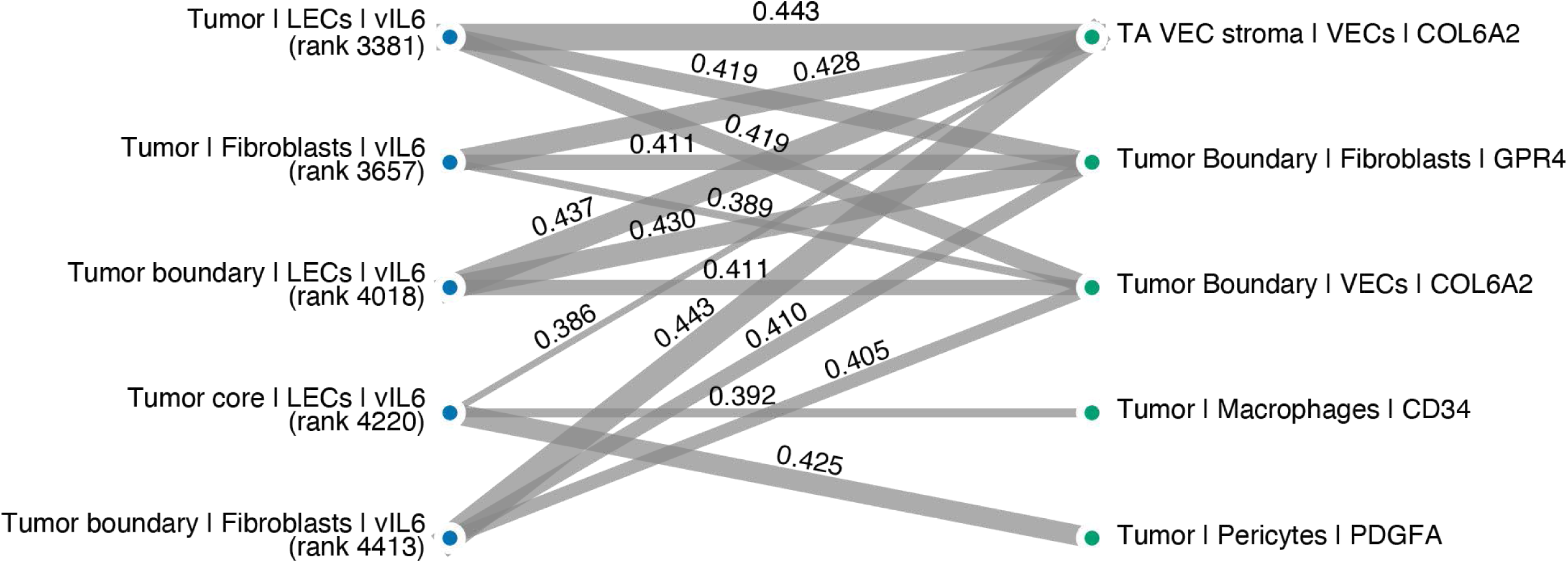
Five highest ranked vIL6 features and their most correlated top predictive spatial single-cell features in Kaposi’s sarcoma. Pearson correlations are shown on top of the edges.

## References

1 Chang, Y. et al. Identification of herpesvirus-like DNA sequences in AIDS-associated Kaposi’s sarcoma. Science 266, 1865–1869, doi:10.1126/science.7997879 (1994).

2 Jary, A. et al. Kaposi’s Sarcoma-Associated Herpesvirus, the Etiological Agent of All Epidemiological Forms of Kaposi’s Sarcoma. Cancers (Basel) 13, doi:10.3390/cancers13246208 (2021).

3 Dzobo, K. The Role of Viruses in Carcinogenesis and Molecular Targeting: From Infection to Being a Component of the Tumor Microenvironment. OMICS 25, 358–371, doi:10.1089/omi.2021.0052 (2021).

4 Gramolelli, S. & Ojala, P. M. Kaposi’s sarcoma herpesvirus-induced endothelial cell reprogramming supports viral persistence and contributes to Kaposi’s sarcoma tumorigenesis. Curr Opin Virol 26, 156–162, doi:10.1016/j.coviro.2017.09.002 (2017).

5 van der Meulen, E., Anderton, M., Blumenthal, M. J. & Schafer, G. Cellular Receptors Involved in KSHV Infection. Viruses 13, doi:10.3390/v13010118 (2021).

6 Zhang, W. & Gao, S. J. Exploitation of Cellular Cytoskeletons and Signaling Pathways for Cell Entry by Kaposi’s Sarcoma-Associated Herpesvirus and the Closely Related Rhesus Rhadinovirus. Pathogens 1, 102–127, doi:10.3390/pathogens1020102 (2012).

7 Douglas, J. L., Gustin, J. K., Moses, A. V., Dezube, B. J. & Pantanowitz, L. Kaposi Sarcoma Pathogenesis: A Triad of Viral Infection, Oncogenesis and Chronic Inflammation. Transl Biomed 1 (2010).

8 Riva, G., Barozzi, P., Torelli, G. & Luppi, M. Immunological and inflammatory features of Kaposi’s sarcoma and other Kaposi’s sarcoma-associated herpesvirus/human herpesvirus 8-associated neoplasias. AIDS Rev 12, 40–51 (2010).

9 He, M. et al. Molecular Biology of KSHV in Relation to HIV/AIDS-Associated Oncogenesis. Cancer Treat Res 177, 23–62, doi:10.1007/978-3-030-03502-0_2 (2019).

10 Grossmann, C., Podgrabinska, S., Skobe, M. & Ganem, D. Activation of NF-kappaB by the latent vFLIP gene of Kaposi’s sarcoma-associated herpesvirus is required for the spindle shape of virus-infected endothelial cells and contributes to their proinflammatory phenotype. J Virol 80, 7179–7185, doi:10.1128/JVI.01603-05 (2006).

11 Lee, M. S. et al. Human Mesenchymal Stem Cells of Diverse Origins Support Persistent Infection with Kaposi’s Sarcoma-Associated Herpesvirus and Manifest Distinct Angiogenic, Invasive, and Transforming Phenotypes. mBio 7, e02109–02115, doi:10.1128/mBio.02109-15 (2016).

12 Qian, L. W., Xie, J., Ye, F. & Gao, S. J. Kaposi’s sarcoma-associated herpesvirus infection promotes invasion of primary human umbilical vein endothelial cells by inducing matrix metalloproteinases. J Virol 81, 7001–7010, doi:10.1128/JVI.00016-07 (2007).

13 Xie, J., Pan, H., Yoo, S. & Gao, S. J. Kaposi’s sarcoma-associated herpesvirus induction of AP-1 and interleukin 6 during primary infection mediated by multiple mitogen-activated protein kinase pathways. J Virol 79, 15027–15037, doi:10.1128/JVI.79.24.15027-15037.2005 (2005).

14 Ye, F. C. et al. Kaposi’s sarcoma-associated herpesvirus promotes angiogenesis by inducing angiopoietin-2 expression via AP-1 and Ets1. J Virol 81, 3980–3991, doi:10.1128/JVI.02089-06 (2007).

15 Morini, M. et al. Kaposi’s sarcoma cells of different etiologic origins respond to HIV-Tat through the Flk-1/KDR (VEGFR-2): relevance in AIDS-KS pathology. Biochem Biophys Res Commun 273, 267–271, doi:10.1006/bbrc.2000.2941 (2000).

16 Simonart, T. et al. Expression of the fibroblast/macrophage marker 1B10 by spindle cells in Kaposi’s sarcoma lesions and by Kaposi’s sarcoma-derived tumor cells. J Cutan Pathol 29, 72–78, doi:10.1034/j.1600-0560.2002.290202.x (2002).

17 Cancian, L., Hansen, A. & Boshoff, C. Cellular origin of Kaposi’s sarcoma and Kaposi’s sarcoma-associated herpesvirus-induced cell reprogramming. Trends Cell Biol 23, 421–432, doi:10.1016/j.tcb.2013.04.001 (2013).

18 Jones, T. et al. Direct and efficient cellular transformation of primary rat mesenchymal precursor cells by KSHV. J Clin Invest 122, 1076–1081, doi:10.1172/JCI58530 (2012).

19 Li, Y. et al. Evidence for Kaposi Sarcoma Originating from Mesenchymal Stem Cell through KSHV-induced Mesenchymal-to-Endothelial Transition. Cancer Res 78, 230–245, doi:10.1158/0008-5472.CAN-17-1961 (2018).

20 Lacunza, E. et al. Unveiling the role of KSHV-infected human mesenchymal stem cells in Kaposi’s sarcoma initiation. J Med Virol 96, e29684, doi:10.1002/jmv.29684 (2024).

21 Carroll, P. A., Brazeau, E. & Lagunoff, M. Kaposi’s sarcoma-associated herpesvirus infection of blood endothelial cells induces lymphatic differentiation. Virology 328, 7–18, doi:10.1016/j.virol.2004.07.008 (2004).

22 Hong, Y. K. et al. Lymphatic reprogramming of blood vascular endothelium by Kaposi sarcoma-associated herpesvirus. Nat Genet 36, 683–685, doi:10.1038/ng1383 (2004).

23 Wang, H. W. et al. Kaposi sarcoma herpesvirus-induced cellular reprogramming contributes to the lymphatic endothelial gene expression in Kaposi sarcoma. Nat Genet 36, 687–693, doi:10.1038/ng1384 (2004).

24 O’Connell, K. M. Kaposi’s sarcoma: histopathological study of 159 cases from Malawi. J Clin Pathol 30, 687–695, doi:10.1136/jcp.30.8.687 (1977).

25 Nadji, M., Morales, A. R., Ziegles-Weissman, J. & Penneys, N. S. Kaposi’s sarcoma: immunohistologic evidence for an endothelial origin. Arch Pathol Lab Med 105, 274–275 (1981).

26 Beckstead, J. H., Wood, G. S. & Fletcher, V. Evidence for the origin of Kaposi’s sarcoma from lymphatic endothelium. Am J Pathol 119, 294–300 (1985).

27 Rutgers, J. L. et al. The expression of endothelial cell surface antigens by AIDS-associated Kaposi’s sarcoma. Evidence for a vascular endothelial cell origin. Am J Pathol 122, 493–499 (1986).

28 Choi, D. et al. The Lymphatic Cell Environment Promotes Kaposi Sarcoma Development by Prox1-Enhanced Productive Lytic Replication of Kaposi Sarcoma Herpes Virus. Cancer Res 80, 3130–3144, doi:10.1158/0008-5472.CAN-19-3105 (2020).

29 Regezi, J. A. et al. Human immunodeficiency virus-associated oral Kaposi’s sarcoma. A heterogeneous cell population dominated by spindle-shaped endothelial cells. Am J Pathol 143, 240–249 (1993).

30 Tse, D. & Stan, R. V. Morphological heterogeneity of endothelium. Semin Thromb Hemost 36, 236–245, doi:10.1055/s-0030-1253447 (2010).

31 Bielenberg, D. R. & D’Amore, P. A. Angiogenesis: Biology and Pathology, Second Edition. Cold Spring Harb Perspect Med 15, doi:10.1101/cshperspect.a041779 (2025).

32 Yang, Y. et al. CXCL12-CXCR4/CXCR7 Axis in Cancer: from Mechanisms to Clinical Applications. Int J Biol Sci 19, 3341–3359, doi:10.7150/ijbs.82317 (2023).

33 Gomez-Herranz, M., Taylor, J. & Sloan, R. D. IFITM proteins: Understanding their diverse roles in viral infection, cancer, and immunity. J Biol Chem 299, 102741, doi:10.1016/j.jbc.2022.102741 (2023).

34 Galati, D. et al. Immunotherapies Targeting CD123 and CD303: A New Frontier in Treating Blastic Plasmacytoid Dendritic Cell Neoplasm. Int J Mol Sci 26, doi:10.3390/ijms26062732 (2025).

35 Khachigian, L. M. et al. Transcriptional regulation of vascular smooth muscle cell proliferation, differentiation and senescence: Novel targets for therapy. Vascul Pharmacol 146, 107091, doi:10.1016/j.vph.2022.107091 (2022).

36 Ye, F., Lei, X. & Gao, S. J. Mechanisms of Kaposi’s Sarcoma-Associated Herpesvirus Latency and Reactivation. Adv Virol 2011, doi:10.1155/2011/193860 (2011).

37 Komaki, S., Inagaki, T., Kumar, A. & Izumiya, Y. The Role of vIL-6 in KSHV-Mediated Immune Evasion and Tumorigenesis. Viruses 16, doi:10.3390/v16121900 (2024).

38 Bu, W. et al. Identification of direct transcriptional targets of the Kaposi’s sarcoma-associated herpesvirus Rta lytic switch protein by conditional nuclear localization. J Virol 82, 10709–10723, doi:10.1128/JVI.01012-08 (2008).

39 Hu, D. et al. Induction of Kaposi’s Sarcoma-Associated Herpesvirus-Encoded Viral Interleukin-6 by X-Box Binding Protein 1. J Virol 90, 368–378, doi:10.1128/JVI.01192-15 (2016).

40 Chatterjee, M., Osborne, J., Bestetti, G., Chang, Y. & Moore, P. S. Viral IL-6-induced cell proliferation and immune evasion of interferon activity. Science 298, 1432–1435, doi:10.1126/science.1074883 (2002).

41 Breiteneder-Geleff, S. et al. Angiosarcomas express mixed endothelial phenotypes of blood and lymphatic capillaries: podoplanin as a specific marker for lymphatic endothelium. Am J Pathol 154, 385–394, doi:10.1016/S0002-9440(10)65285-6 (1999).

42 Goerdt, S. & Sorg, C. Endothelial heterogeneity and the acquired immunodeficiency syndrome: a paradigm for the pathogenesis of vascular disorders. Clin Investig 70, 89–98, doi:10.1007/BF00227347 (1992).

43 Kaaya, E. E. et al. Heterogeneity of spindle cells in Kaposi’s sarcoma: comparison of cells in lesions and in culture. J Acquir Immune Defic Syndr Hum Retrovirol 10, 295–305 (1995).

44 Ribatti, D., Tamma, R., Ruggieri, S., Annese, T. & Crivellato, E. Surface markers: An identity card of endothelial cells. Microcirculation 27, e12587, doi:10.1111/micc.12587 (2020).

45 Purushothaman, P., Uppal, T., Sarkar, R. & Verma, S. C. KSHV-Mediated Angiogenesis in Tumor Progression. Viruses 8, doi:10.3390/v8070198 (2016).

46 Diaz-Flores, L. et al. Delimiting CD34+ Stromal Cells/Telocytes Are Resident Mesenchymal Cells That Participate in Neovessel Formation in Skin Kaposi Sarcoma. Int J Mol Sci 24, doi:10.3390/ijms24043793 (2023).

47 Gurzu, S., Ciortea, D., Munteanu, T., Kezdi-Zaharia, I. & Jung, I. Mesenchymal-to-endothelial transition in Kaposi sarcoma: a histogenetic hypothesis based on a case series and literature review. PLoS One 8, e71530, doi:10.1371/journal.pone.0071530 (2013).

48 Pyakurel, P. et al. Lymphatic and vascular origin of Kaposi’s sarcoma spindle cells during tumor development. Int J Cancer 119, 1262–1267, doi:10.1002/ijc.21969 (2006).

49 Chen, Z. & Zhao, B. The role of tumor-associated macrophages in HPV induced cervical cancer. Front Immunol 16, 1586806, doi:10.3389/fimmu.2025.1586806 (2025).

50 Khan, M. N. et al. Tumor-associated macrophages and CD8+ T cells: dual players in the pathogenesis of HBV-related HCC. Front Immunol 15, 1472430, doi:10.3389/fimmu.2024.1472430 (2024).

51 Ding, C. & Peng, H. Minimum redundancy feature selection from microarray gene expression data. J Bioinform Comput Biol 3, 185–205, doi:10.1142/s0219720005001004 (2005).

52 Bettuzzi, T., Lebbe, C. & Grolleau, C. Modern Approach to Manage Patients With Kaposi Sarcoma. J Med Virol 97, e70294, doi:10.1002/jmv.70294 (2025).

53 Staskus, K. A. et al. Kaposi’s sarcoma-associated herpesvirus gene expression in endothelial (spindle) tumor cells. J Virol 71, 715–719, doi:10.1128/JVI.71.1.715-719.1997 (1997).

54 Morris, V. A. et al. The KSHV viral IL-6 homolog is sufficient to induce blood to lymphatic endothelial cell differentiation. Virology 428, 112–120, doi:10.1016/j.virol.2012.03.013 (2012).

55 Aoki, Y. et al. Angiogenesis and hematopoiesis induced by Kaposi’s sarcoma-associated herpesvirus-encoded interleukin-6. Blood 93, 4034–4043 (1999).

56 Fielding, C. A. et al. Viral IL-6 blocks neutrophil infiltration during acute inflammation. J Immunol 175, 4024–4029, doi:10.4049/jimmunol.175.6.4024 (2005).

57 Vart, R. J. et al. Kaposi’s sarcoma-associated herpesvirus-encoded interleukin-6 and G-protein-coupled receptor regulate angiopoietin-2 expression in lymphatic endothelial cells. Cancer Res 67, 4042–4051, doi:10.1158/0008-5472.CAN-06-3321 (2007).

58 Zhu, X. et al. Synergy between Kaposi’s sarcoma-associated herpesvirus (KSHV) vIL-6 and HIV-1 Nef protein in promotion of angiogenesis and oncogenesis: role of the AKT signaling pathway. Oncogene 33, 1986–1996, doi:10.1038/onc.2013.136 (2014).

59 MacPhail, L. A., Dekker, N. P. & Regezi, J. A. Macrophages and vascular adhesion molecules in oral Kaposi’s sarcoma. J Cutan Pathol 23, 464–472, doi:10.1111/j.1600-0560.1996.tb01436.x (1996).

60 Ramos da Silva, S., et al. Broad Severe Acute Respiratory Syndrome Coronavirus 2 Cell Tropism and Immunopathology in Lung Tissues From Fatal Coronavirus Disease 2019. J Infect Dis 223, 1842–1854, doi:10.1093/infdis/jiab195 (2021).

61 Gao, S. J., Deng, J. H. & Zhou, F. C. Productive lytic replication of a recombinant Kaposi’s sarcoma-associated herpesvirus in efficient primary infection of primary human endothelial cells. J Virol 77, 9738–9749, doi:10.1128/jvi.77.18.9738-9749.2003 (2003).

62 Brulois, K. F. et al. Construction and manipulation of a new Kaposi’s sarcoma-associated herpesvirus bacterial artificial chromosome clone. J Virol 86, 9708–9720, doi:10.1128/JVI.01019-12 (2012).

63 Gao, S. J. et al. Seroconversion to antibodies against Kaposi’s sarcoma-associated herpesvirus-related latent nuclear antigens before the development of Kaposi’s sarcoma. N Engl J Med 335, 233–241, doi:10.1056/NEJM199607253350403 (1996).

64 Ester, M., Kriegel, H.-P., Sander, J. & Xu, X. W. A density-based algorithm for discovering clusters in large spatial databases with noise. KDD 96, 226–231 (1996).

65 Abdi, H. & Williams, L. J. Principal component analysis. Wiley interdisciplinary Reviews: Computational Statistics 2, 433–459 (2010).

66. 66 Nolet, C. et al. Accelerating single-cell genomic analysis with GPUs. bioRxiv 2022.05.26.493607, doi: 10.1101/2022.1105.1126.493607 (2022).

67 Ootomo, H. et al. CAGRA: Highly Parallel Graph Construction and Approximate Nearest Neighbor Search for GPUs. 2024 IEEE 40th International Conference on Data Engineering (ICDE), Utrecht, Netherlands, 2024 doi: 10.1109/ICDE60146.2024.00323, 4236-4247 (2024).

68 Traag, V. A., Waltman, L. & van Eck, N. J. From Louvain to Leiden: guaranteeing well-connected communities. Sci Rep 9, 5233, doi:10.1038/s41598-019-41695-z (2019).

69 Ding, J. et al. Systematic comparison of single-cell and single-nucleus RNA-sequencing methods. Nat Biotechnol 38, 737–746, doi:10.1038/s41587-020-0465-8 (2020).

70 Murtagh, F. & Contreras, P. Algorithms for hierarchical clustering: an overview. WIREs Data Mining and Knowledge Discovery 2, 86–97 (2012).

71 Das, A. et al. Molecular and immune signatures, and pathological trajectories of fatal COVID-19 lungs defined by in situ spatial single-cell transcriptome analysis. J Med Virol 95, e29009, doi:10.1002/jmv.29009 (2023).

72 Hartigan, J. A. & Wong, M. A. A K-Means Clustering Algorithm. Applied Statistics 28, 100–108 (1979).

73. Getis, A. Spatial Autocorrelation. In: Fischer, M., Getis, A. (eds) Handbook of Applied Spatial Analysis. Springer, Berlin, Heidelberg 10.1007/978-3-642-03647-7_14 (2010).

74 Moon, K. R. et al. Visualizing structure and transitions in high-dimensional biological data. Nat Biotechnol 37, 1482–1492, doi:10.1038/s41587-019-0336-3 (2019).

75 Street, K. et al. Slingshot: cell lineage and pseudotime inference for single-cell transcriptomics. BMC Genomics 19, 477, doi:10.1186/s12864-018-4772-0 (2018).

76 Van den Berge, K., et al. Trajectory-based differential expression analysis for single-cell sequencing data. Nat Commun 11, 1201, doi:10.1038/s41467-020-14766-3 (2020).

77. Gillies, S.,> et al. Shapely (2.1.1). Zenodo 10.5281/zenodo.15463269 (2025).

78 Campello, R. J. G. B., Moulavi, D. & Sander, J. Density-Based Clustering Based on Hierarchical Density Estimates. Advances in Knowledge Discovery and Data Mining **Conference paper**, 160–172 (2013).

79 Berg, M., Cheong, O., Kreveld, M. & Overmars, M. Convex Hulls. In: Computational Geometry. Springer, Berlin, Heidelberg. 10.1007/978-3-540-77974-2_11, 243-258 (2008).

80 Dice, L. R. Measures of the amount of ecologic association between species. Ecology 26, 297–302 (1945).

81. Chen, T. Q. & Guestrin, C. XGBoost: A Scalable Tree Boosting System. KDD ’16: Proceedings of the 22nd ACM SIGKDD International Conference on Knowledge Discovery and Data Mining 10.1145/2939672.293978, 785-794 (2016).

82 Radovic, M., Ghalwash, M., Filipovic, N. & Obradovic, Z. Minimum redundancy maximum relevance feature selection approach for temporal gene expression data. BMC Bioinformatics 18, 9, doi:10.1186/s12859-016-1423-9 (2017).

